# Mechanisms of enhanced or impaired DNA target selectivity driven by protein dimerization

**DOI:** 10.1101/2025.02.18.638941

**Authors:** Mankun Sang, Gabriel Au, Margaret E. Johnson

## Abstract

Successful DNA transcription demands coordination between proteins that bind DNA while simultaneously binding to one another to form dimers or higher-order complexes. For proteins with numerous DNA targets throughout the genome, measurements that report on their dwell time or occupancy thus represent a convolution over a population interacting with specific DNA, nonspecific DNA, or protein partners on DNA. Dimerization is known to add contacts that can help a single protein to stably bind DNA. However, we show here that dimerization can also impair measured dwell times and occupancy on target sequences because the population redistributes across DNA. We combine mass-action kinetic models of pairwise reversible reactions between proteins and DNA with theory and spatial stochastic simulations to isolate the role of dimerization on observed DNA dwell times, occupancy, and spatial distribution of proteins on DNA. Three key themes emerge: (i) Protein-protein interactions, in addition to protein-DNA interactions, can localize a protein to DNA, and relative binding rates can thus widely tune dwell times. (ii) Dimensional reduction achieved through nonspecific binding and subsequent 1D diffusion controls the order-of-magnitude of enhancements despite nucleosome barriers. (iii) Dimerization enhances selectivity for locally clustered targets and often impairs binding to widely-spaced targets by sequestration. Compared with ChIP-seq data, our model explains how the distribution of the essential GAF protein throughout the genome is highly selective for clustered targets due to protein interactions. This model framework predicts when even weak dimerization can redistribute and stabilize proteins on DNA as a necessary part of transcription.

**Significance:** Transcription of DNA into RNA relies on a globally orchestrated but microscopically stochastic recruitment of multiple proteins to targeted DNA sequences. DNA binding is only a first step. Here we show how proteins with many targets throughout the genome can use protein-protein dimerization to not only help proteins bind DNA, but also to redistribute the population genome-wide, significantly enhancing or impairing dwell times and target occupancies dependent on target spacing. Using theory and simulations along with experimental comparison, outcomes depend on protein-to-target ratios, dimensional gains of 1D searchers, and relative binding rates. Our relatively simple formulas predict how these factors yield negligible or dramatic changes in target selection, with direct mapping to experimental measurements on essential transcription factors.

## Introduction

The transcription of DNA into RNA is essential for all living systems and relies on the concerted action of a variety of DNA-binding proteins[1-4]. While some of these proteins are highly specialized and recognize ∼1 target in the genome, here we focus on ‘generalist’ proteins, motivated by the essential GAGA Factor GAF[5], that bind specific DNA sequences or motifs that are widely distributed in the genome. These proteins may therefore have multiple ‘choices’ about where or when to bind, and key metrics of proper function include dwell times[1, 6] and occupancies on DNA[7, 8]. These multi-domain proteins can arrive at DNA sites as monomers [5, 9-11], dynamically recruited not only via specific or nonspecific DNA interactions[12-14] but via protein-protein interactions[12-14]. Experimental measurements that report on the lifetime and occupancy of populations of such multi-domain proteins on DNA thus reflect the combined effects of both DNA and protein interactions[15-17]. Two-state models of a protein on DNA typically assume specific and nonspecific binding modes [6, 18, 19]. However, without incorporating protein-protein interactions, these models cannot predict responses to mutations in protein interacting domains[20] or responses to expression level changes that must impact dimerization. A key insight from our modeling approach is that protein localization and immobilization on DNA can arise primarily from protein-protein interactions, without requiring direct DNA binding. Our focus here is to systematically isolate and understand how dimerization will quantitatively enhance or impair DNA target occupancy, dwell time, and selectivity towards clustered targets. We define a minimal model of reversible protein dimerization, incorporating DNA binding and diffusion in 3D and 1D. Using mass-action kinetics, theory, and particle-based reaction-diffusion simulations, we show how diverse outcomes emerge dependent on the ratio of proteins to targets, dimensional gains of 1D interactions, and relative binding rates.

Our model explicitly captures a key feature of many DNA-binding proteins: their nonspecific association with the charged DNA backbone[21], which facilitates 1D diffusion[18]. This sliding or hopping along DNA accelerates a protein search for a target sequence, a mechanism exploited by prokaryotic [22-24] and eukaryotic proteins[25], despite the latter’s frequent interruption along the 1D DNA path by nucleosomes[26]. Nonspecific association causes a longer lifetime and stronger affinity to DNA as compared to binding isolated target sequences[26, 27]. When two proteins localize to DNA, their dimerization becomes a 1D search as long as at least one partner can slide. Like on 2D membranes, this dimensional reduction will typically increase the collision probability compared to 3D [28], promoting significantly more stable and long-lived dimer and membrane- or DNA-bound populations at equilibrium[29, 30].

Building from the seminal work of Berg and Von Hippel[6, 22], our model allows nonspecifically adhered proteins to encounter and bind DNA targets and other 1D-associated proteins using 1D rates. To support reversible binding and retain detailed balance, we assume collisions are not always productive for binding. This is consistent with evidence of proteins sliding over targets without binding[24, 31], and with protein-protein association rates, which are rarely diffusion-limited[32]. With both 1D and 3D association in our model, the ratio of a protein’s local volume *V* to the accessible DNA length *L* between nucleosomes represents a critical factor controlling lifetimes and occupancy, but we will consider this fixed based on the Drosophila nucleus and genome size at *V*/*L*=1000 nm^2^ (see SX.B in SI). The *V*/*L* ratio has units of area and demands a corresponding ratio of on-rates, 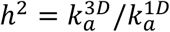. The magnitude of *h*^2^ will depend on the specific binding pair, but we will constrain it to the 1-1000 nm^2^ range, which is reasonable when localization to low dimensional spaces entropically restricts configurations without significant allosteric effects[33-35]. Collectively we define a dimensionality factor,

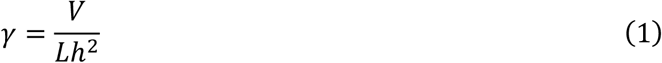

that here is greater than 1, which means the bound state is more stable relative to monomers in 1D compared to the 3D system[29].

Although dimerization between transcription factors is known to bring together DNA-binding domains that effectively switch-on DNA binding[36] (e.g. bZIP monomers[37, 38]), our model encompasses a much broader range of mechanisms for dimerization to impact target selectivity. We account for the fact that genome-wide measurements report on a population of competing proteins, not single molecules, and establish regimes where dimerization impairs target binding via sequestration, and where dimerization has minimal impact on mean dwell times despite altering the distribution of proteins on and off DNA. Critically, we quantify how dimerization (or higher-order oligomerization) significantly enhances selectivity for clustered targets, beyond what’s expected based solely on an increased density of target sites. While clustered targets in the genome (commonly found in promoters and enhancers[39, 40]) are known to enhance transcription factor binding[39-41], even monomers will naturally select for a cluster over a single target due to the higher target density. However, we here quantify how protein-protein interactions can dramatically enhance dwell times on DNA and show that this population effect diminishes for widely spaced targets[42, 43]. For the oligomer-forming GAF protein, ChIP-seq data supports selectivity for clustered targets in *Drosophila* [20, 44], and our model directly explains how protein-protein interactions are responsible for enhancing these localization inhomogeneities throughout DNA. By isolating the impact of dimerization from the stabilizing effects of specific and nonspecific DNA binding, our model provides a way to improve interpretation of single-particle tracking measurements and predict how both DNA-binding and protein-protein interaction domains control DNA target selection.

### Models and Theory Model Description

Our model contains two components: proteins and DNA. Protein species *P* can diffuse and have three interaction sites, one binds another protein partner, one nonspecifically binds to DNA, and one binds to specific DNA sites. Instead of a continuous sequence, DNA is a 1D line composed of two discrete species, specific sites *S* and nonspecific sites *N* (Fig. S1). While nonspecific sites are implicitly assumed to diffuse along the line, specific sites remain stationary. Discrete, diffusing nonspecific sites allow our (nonspatial) mass-action kinetic models to describe protein localization to DNA and the ability to slide (diffuse) along the backbone in 1D to encounter the immobile *S* sites. To be concrete, our model parameters are based on GAF from Drosophila[20]. We determine the number of *N* sites based on a line density of 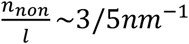 (Fig. S1). A key assumption of most results is that diffusion is not limiting, hence measured timescales are controlled by populations and rate constants. This is a reasonable approximation given our domain sizes and supported by our spatial simulations that relax this assumption: we assume 1D stretches are relatively short (∼60nm), because we consider nucleosomes as blocks to 1D diffusion. Our spatial stochastic, particle-based reaction-diffusion simulations [45] similarly treat nonspecific sites as (explicitly) diffusing particles.

The number of specific targets *S* in a segment of DNA (between 2 nucleosomes or ∼100 basepairs) is highly variable throughout the genome. For example, GAF has ∼10^5^ segments and ∼15,000 target motifs (see SX.C, SX.E in SI). So excepting the final CHiP-seq comparison, we model DNA segments with discrete integer copies of sites, either *S* = 0 or *S* = 2. We thus define models representing these distinct different DNA environments (Fig. 1c), and solve for the local equilibrium of these microenvironments, before considering competition between them. For our ‘DNA’ model, *S* = 0. For the ‘DNA + targ′ model, *S* = 2, but the sites are not adjacent, and thus a single protein dimer cannot bind both targets simultaneously. For the ‘DNA + clusTarg′ model, *S* = 2 and they are clustered, allowing a single dimer to bind both sites simultaneously. All three models have the same segment length, same *N* + *S*, and the same *V*/*L* ratio of 1000nm^2^ for Drosophila (see SX.B in SI for justification). For concentrations, we thus typically use [*S*] = 0 or 158 µM, and [*N*] = 948.9 or 791 µM.

**Figure 1.**
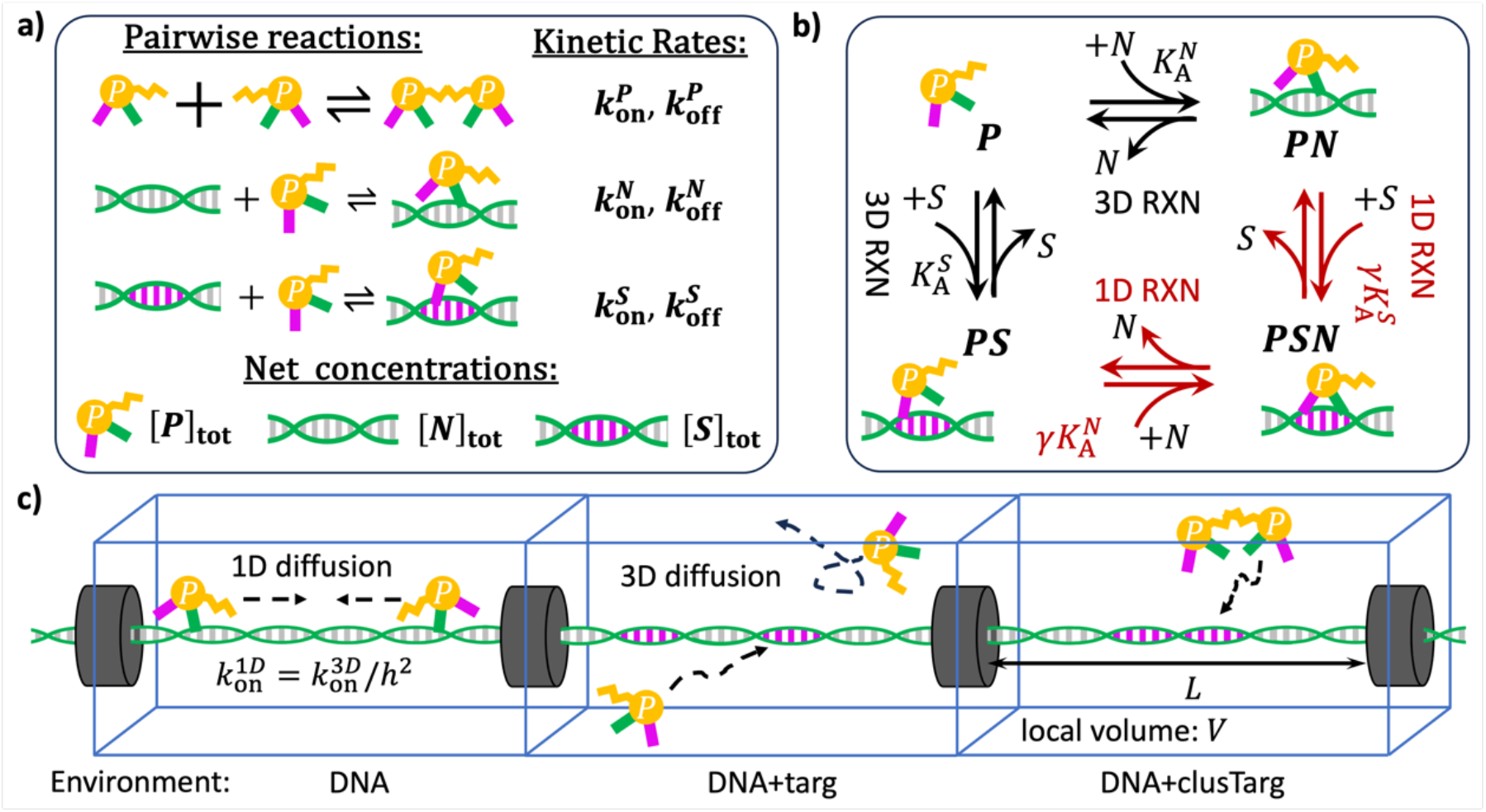
Microstate model of pairwise interactions for proteins and DNA in distinct DNA microenvironments. a) The fundamental pairwise reactions between proteins (yellow), nonspecific/random DNA sequences (green) and specific target DNA sequences (magenta). Each target only supports a single monomer contact. Average dwell times and occupancies will depend on rates 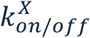 and species concentrations. b) The reaction network for monomers including 3D reactions (black arrows) and 1D reactions (red arrows) enforces detailed balance. *PSN* labels proteins that formed both target and nonspecific contacts. These state definitions enable transit between *PN* and *PS* without returning to the 3D bulk, and to conserve mass. The appearance of *γ* indicates a 1D reaction, as the partition coefficient 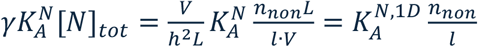 is independent of *V*. c) Three environments capture distinct distributions of targets (magenta bases) on DNA. The black cylinders are nucleosomes. Proteins bound to nonspecific DNA sequences perform 1D diffusion, but proteins bound to targets are fixed.

Our model has 7 rate parameters. Six of the rate parameters are from the fundamental pairwise or bimolecular interactions between our species: protein-protein dimerization *P* + *P* ⇌ *PP* with rates 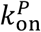 and 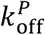, protein nonspecific binding to DNA (*P* + *N* ⇌ *PN*) with rates 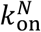 and 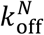, and protein binding to specific DNA (*P* + *S* ⇌ *PS*) with rates 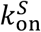 and 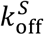 (Fig. 1a). We use macroscopic rates that incorporate diffusion times [46], and we show using particle-based reaction-diffusion simulations with intrinsic rates 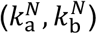 that this approximation is valid in the rate regimes we study (binding is not diffusion-limited) and in the spatial domains we consider (protein diffusion is fast enough to keep the system well-mixed) (Fig. S2). We note that recruitment fractions to DNA and target occupancy are independent of diffusion and kinetic rates because we operate at equilibrium, and 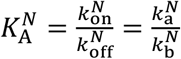 [46]. Because each protein *P* has three distinct and non-competing interaction sites, reactions with *P, S*, and *N* sites combine to form multiple states, and in our micro-kinetic framework, we enumerate all of these states, with some of these reactions occurring in 1D. Shown in Fig. 1b, the monomer *P* can bind to DNA using both specific and nonspecific sites, denoted as state *PSN*. Here, either the nonspecific or specific site may bind DNA first, followed by the other. The distinction between state *PSN* and *PS* maintains detailed balance since proteins can both slide onto *S* in 1D or directly bind from 3D. From a molecular perspective, the *PS* + *N* reaction is like a conformational change[47, 48], or unimolecular reaction. However, since the *N* species can diffuse, the change must be represented in our model via a bimolecular reaction.

Dimensional reduction, more specifically the transformation of 3D interactions to a 1D space, must change the *i*. system size and *ii*. reaction on-rate. As noted in the introduction, for system size we have (*i*) *V*/*L* and for rates (*ii*), the 1D on-rate can be defined as,

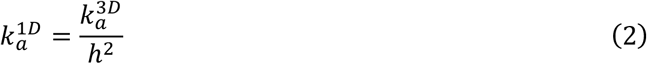

where *h*^2^ is the 7^th^ rate parameter (besides the 6 described in Fig. 1a). The off rates vary at most by a scalar, 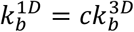 since they do not involve a search. For spatial simulations, these two aspects of dimensional reduction are both explicitly captured. For our coupled nonspatial rate equations, we need to track all species in the same units, and we choose to use 3D units, e.g. [*N*] has units of copies/volume. For all 1D association reactions, we therefore must scale the reaction rate by *γ* (Eq. 1) to capture both the change in reactant concentrations (*V*/*L*) and the change in on-rates (*h*^2^) (see SI in SI for detailed derivation). To minimize additional parameters, we assume that the off-rates are the same from 3D to 1D, 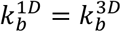 [49, 50], and we use the same value of *h*^2^ for all 1D reactions (see SX.B in SI for further details).

To briefly contrast our three microenvironments (Fig 1c), ‘DNA′ model contains 6 species or microstates: [*P*], [*PP*], [*N*], [*PN*], [*PPN*], [*PNPN*], that participate in 6 reactions (Fig. S3). For ‘DNA + targ′ model, in addition to the 6 species, it contains an additional 7 species: [*S*], [*PS*], [*PSN*], [*PPSN*], [*PPS*], [*PNPS*], [*PNPSN*] and 21 total reactions (Fig. S4, SI SV.D). *PNPS* is a dimer where each constituent monomer binds DNA once. *PPSN* is also a dimer that forms two DNA interactions, but both are on the same monomer. For ‘DNA + clusTarg′ model with target clusters indicated by *S*_2_, there are 10 additional species beyond the 6 species in model DNA, and 27 total reactions (Fig. S5, SI SV.E).

### Two-state model for calculating the mean dwell time

Our three model environments all contain multiple protein-DNA bound microstates (e.g. *PN, PS*, and *PSN*) and unbound states (e.g. *P* and *PP*). To define convenient metrics that compare between models and capture what is typically measured in experiment, we map these microstates to a two-state model. We assume an equilibrium steady state, where the flux in and out of each pairwise binding reaction is equivalent, e.g. 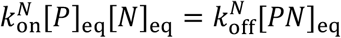 (all rates and concentrations are converted to 3D units by default). All metrics can ultimately be written as functions of our known parameters 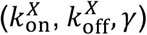 and equilibrium monomer species, [*P*], [*N*], and [*S*]_eq_. The equilibrium populations result from coupled nonlinear equations, and therefore must generally be solved for numerically, but we will assume nonspecific sites in excess, [*N*]_eq_∼[*N*]_tot_, and in some relevant regimes we can approximate [*P*]_eq_ and [*S*]_eq_ to get closed-form expressions (see SI SVI).

To measure dwell time in the bound state of this two-state model, we use the following expression (see SI SIII):

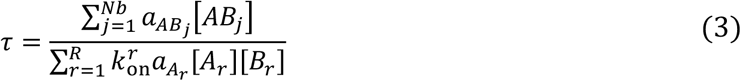

where the numerator sums over all *Nb* bound microstates involving the protein *A* with partner *B* being DNA sites (e.g. *S* and *N*) or proteins on DNA (e.g. *PN*), with stoichiometry *a*_*A*B_. The denominator sums over all *R* reactions that generate flux into these bound microstates from an unbound protein. We show in SIX in SI that calculating *τ* via the survival time distribution *S*(*t*) (see Fig. S6cd for examples of *S*(*t*)) produces the same analytical result as Eq. (3) under steady-state approximations, with the two-state method via Eq. (3) significantly simplifying the analysis.

We define the recruitment fraction of proteins on DNA by,

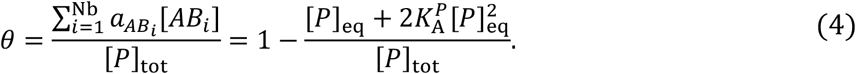

For DNA segments with targets (DNA + targ, DNA + clusTarg), we also quantify the fraction of specific DNA targets occupied by proteins, or target occupancy,

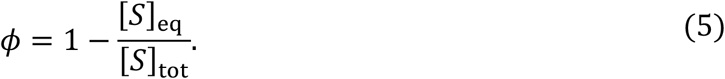

*ϕ* varies from 0 (all targets are free) to 1 (all targets are occupied), with a maximal value for a population of 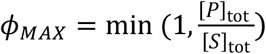.

## Results

### Driven by 1D reactions, dimerization enhances recruitment but does not always increase dwell time on nonspecific DNA

We first consider the simplest DNA environment where proteins can only bind DNA nonspecifically, as it illustrates how dimensional reduction can enhance DNA localization properties. For the dwell time, we find that counterintuitively, dimerization can both enhance and impair lifetimes relative to monomers. The mean dwell time for monomers is 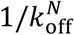. When proteins can also dimerize, the mean dwell time can be written as:

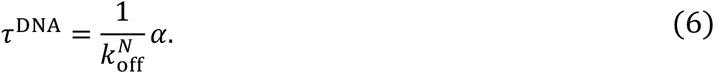

Where the scalar *α* > 1 indicates dimerization has lengthened the dwell time, and vice-versa. For proteins that form irreversible dimers, dimerization always lengthens dwell time (see SVI.B in SI):

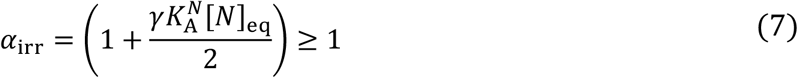

(Fig. 2d). This maximal increase can be as large as 3 orders-of-magnitude depending on the 3D partition coefficient to DNA, 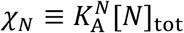 (where [*N*]_eq_ ≅ [*N*]_tot_), which is transformed to its 1D equivalent by *γ*. This key factor *γ*χ_*N*_ is equivalent to 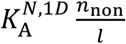 (see SI SII), where 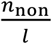 is the line density of *N* sites and 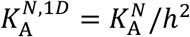 is the 1D affinity for DNA when the protein is localized by a partner to a 1D search (Fig 2). Since we can assume 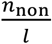 is constant, a higher *γ*χ_*N*_ means a stronger affinity for DNA during a 1D search. In the more general case where protein-protein interactions are reversible, we find (see SVI.B in SI),

**Figure 2.**
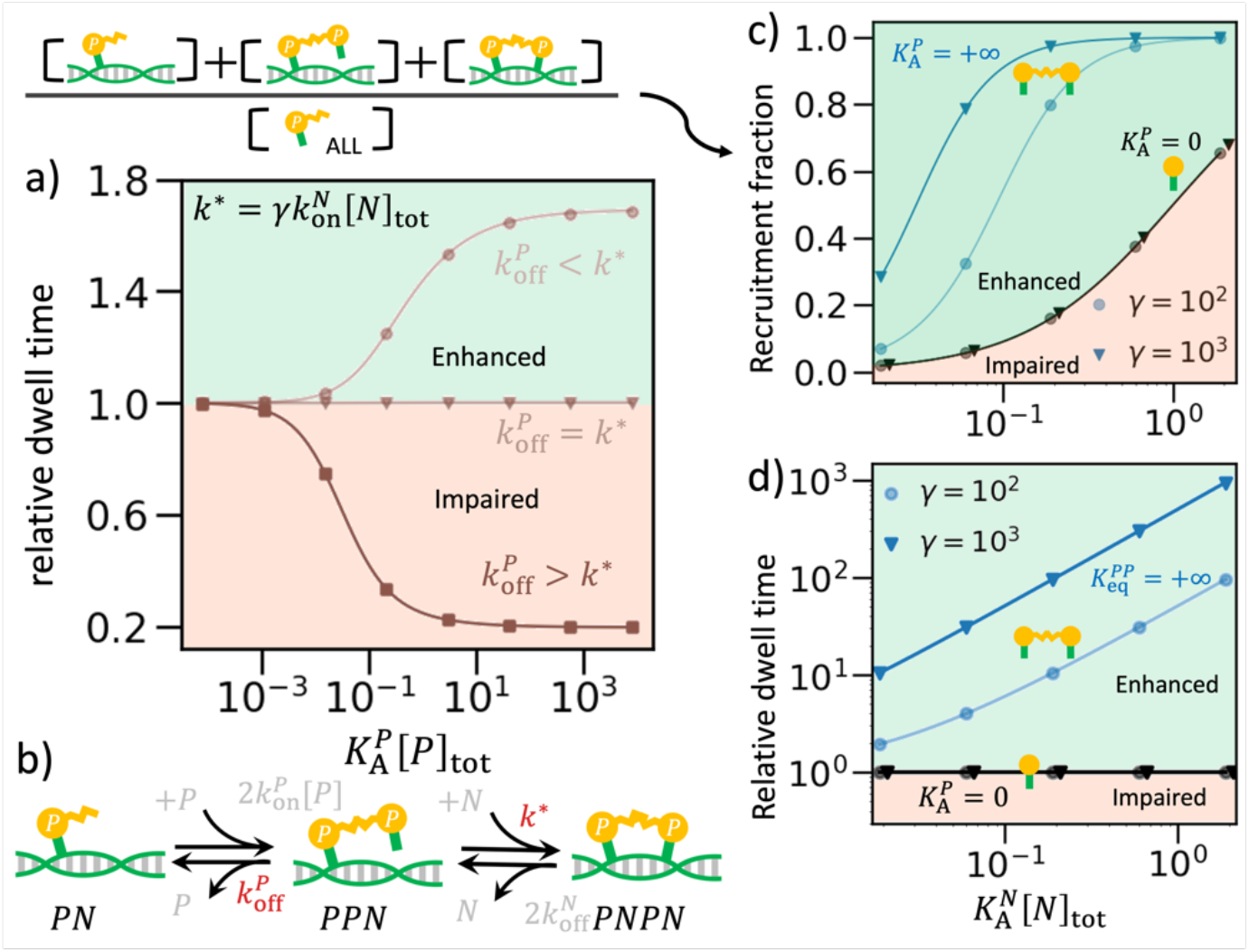
Dimerization enhances recruitment, but can either lengthen or shorten dwell times on nonspecific DNA. In all plots, points are numerical simulations (see SXII.A in SI). a) The dwell time can lengthen due to dimerization (circles), where we define 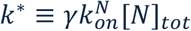. No change occurs when 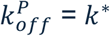 (triangles), and thus are impaired when 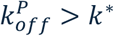 (squares). The degree of enhancement or reduction depends on the partitioning strength of dimerization 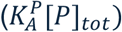. Here 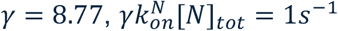, and [*P*] = 7.9 *μM*. From light to dark, 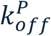 is 0.1 *s*^−1^ (circle), 1 *s*^−1^ (triangle), and 10 *s*^−1^(square). Curves are analytical theory given by Eq. (32’) in SI. b) This system does not include any targets. A protein can be colocalized with DNA by associating an already DNA-bound protein (*P* + *PN* → *PPN*) and then leave the DNA environment with a rate 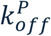 if it does not first bind the DNA with rate *k*^*^. c) The recruitment fraction depends on the strength of nonspecific partitioning, 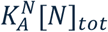 for both monomers (black curves: Eq. (34) in SI) and dimers (blue curves: Eq. 36 in SI). Dimers have better recruitment than monomers, with largest improvement when 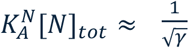, and with larger *γ* = 1000 (triangles) improving recruitment relative to *γ* =100 (circles). d) The dwell time of proteins on DNA relative to the monomer value, 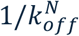, does not change for monomers, as expected (black curves: 1). For irreversible dimers (blue curves: Eq. (35) in SI), orders-of-magnitude increases in dwell time are possible for larger *γ* and more stable partitioning to DNA.

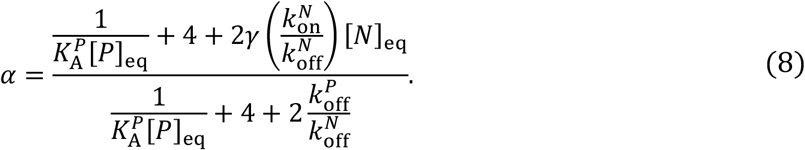

Now we see sometimes *α* < 1, or a shortening of dwell time due to dimerization. This occurs if

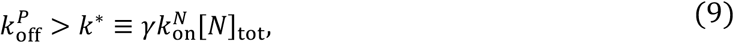

that is, for fast 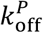 or slow values of 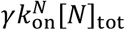 (Fig. 2a). The origin is that multi-domain proteins are still observed to be ‘DNA bound’ even if they are localized to DNA only by another bound protein (Fig. 2b). The lifetimes of these DNA-co-localized proteins can be controlled by the dimer lifetime if it is shorter than the time to form a second protein-DNA bond. Under this condition, stronger dimer affinity increases the fraction of these co-localized proteins that have a short dwell time. Although regimes of impairment are less common than enhancement (nonspecific binding is typically shorter lived than dimer binding, 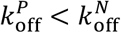), it establishes that co-localizing proteins to an observed rather than explicit DNA-bound state impacts measured timescales.

Unlike the dwell time, the recruitment fraction to DNA, *θ*^*DNA*^, will always increase due to dimerization as (Fig. 2c, SVII.C in SI):

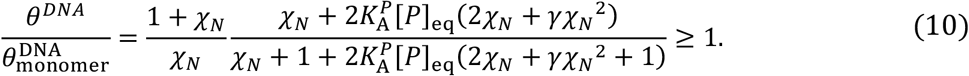

As expected, no enhancement occurs with 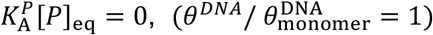. Maximal enhancement occurs for the irreversible dimer, 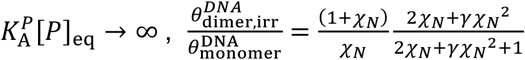(see SVII.C in SI). Here, larger *γ* always drives larger enhancement, as 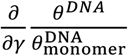 is always positive. However, the enhancement is not monotonic with χ_*N*_ (assuming a fixed *γ* > 2), and the biggest enhancement occurs at an intermediate value χ_*N*_* found by solving, 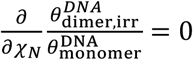:

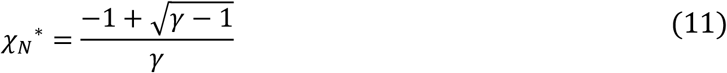

For stronger partitioning, χ_*N*_ > χ_*N*_^*^, monomers do not need ‘help’ to be recruited to DNA, whereas for weaker partitioning, dimerization cannot rescue very weak DNA binding (Fig. 2c). The maximal recruitment fraction can increase by up to 2∼16-fold dependent on *γ*, with 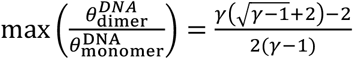. As discussed above, a higher *γ* must follow from stronger 1D affinities 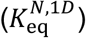 that stabilize dimers on DNA by binding a second *N* site.

### With separated targets, dimerization always enhances recruitment but not always dwell time or target occupancy

With the addition of specific target sites to the DNA, we first assert that targets are sufficiently separated such that a protein dimer can only bind with one target (Fig. 3). In this environment, *without nonspecific binding* and given the same total protein concentration, dimers can only impair target occupancy (*ϕ*) relative to monomers. This is because in this purely 3D binding problem, dimers sequester one protein to be incapable of target binding (see SVIII in SI for details). With nonspecific binding, dimerization can both enhance and impair *ϕ* (Fig. 3d). When targets are in excess of proteins ([S]_tot_ > [P]_tot_) we can show (see SI SVII.A) that the occupancy of irreversible dimers can drop below that of monomers due to sequestration, or 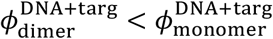 (Fig. 3a). This occurs when the partition coefficient to target sites χ_*S*_ is stronger than a critical value χ^*^:

**Figure 3.**
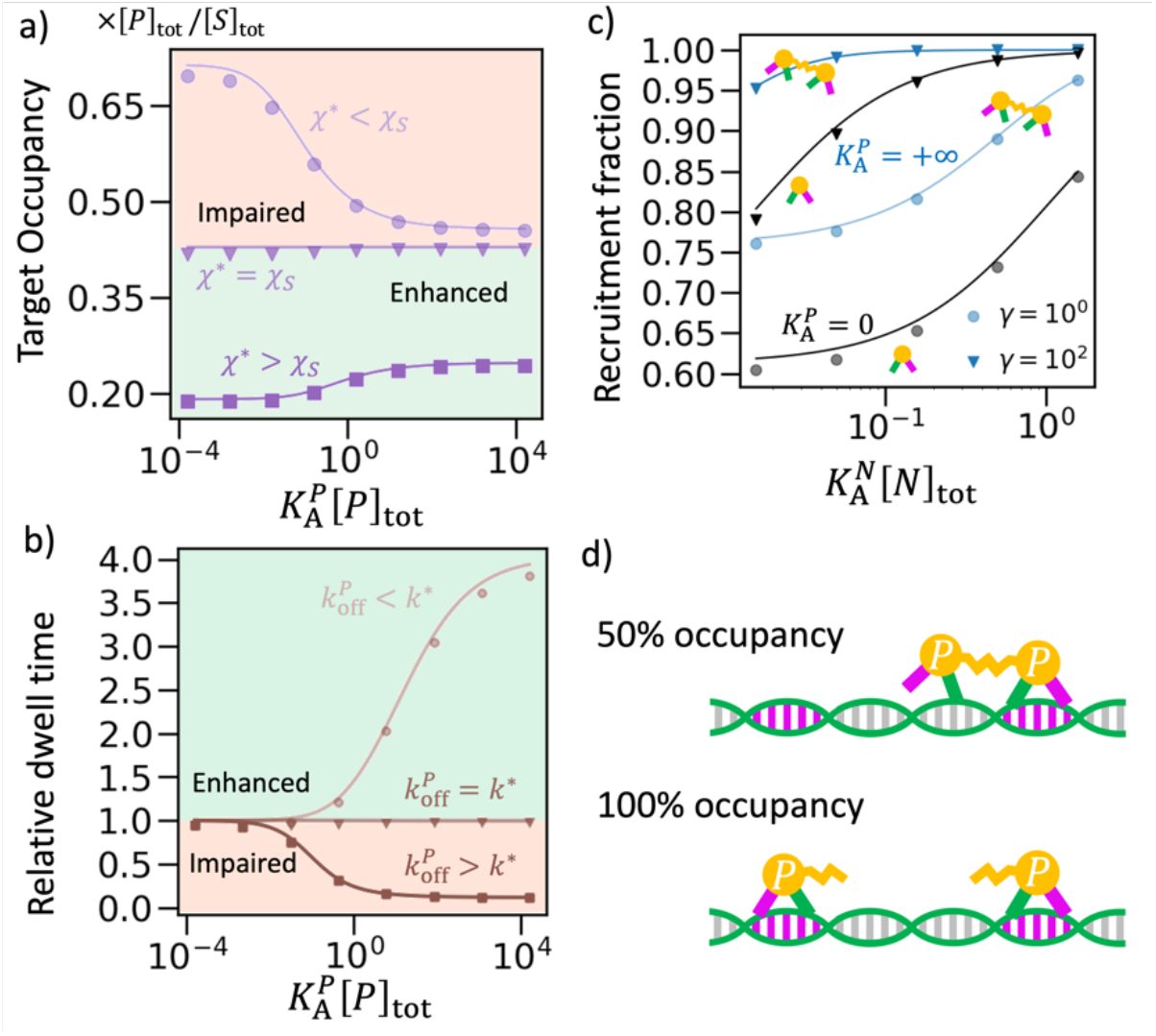
For proteins binding separated targets, dimerization impairs proteins that are already strong target binders. In this figure, [*P*]_*tot*_ = 0.1[*S*]_*tot*_ and numerical results (points) are obtained from solving the ODEs (see SXII.A in SI). a) With χ^*^ = 0.11, the target occupancy is impaired when it’s dominated by direct protein-target binding, namely χ_*S*_ > χ^*^. When χ_*S*_ = χ^*^, our theory predicts unchanged target occupancy with different 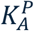. However, numerical results show a minimal enhancement to occupancy due to our approximation [*S*]_*eq*_∼[*S*]_*tot*_. When χ_*S*_ < χ^*^, dimerization enhances occupancy by forming a second nonspecific contact. The analytical results (curves) are given by Eq. (39) in SI. Note that the occupancy is shown here as a fraction of the maximum possible, *ϕ*_*MAX*_ = [*P*]_*tot*_/[*S*]_*tot*_. b) With *k*^*^ = 0.1*s*, dimerization can impair 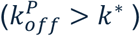 the relative dwell time, 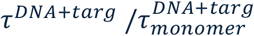, due to fast dissociation of co-localized proteins. Dimerization enhances 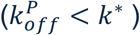 the dwell time when the co-localized protein prefers forming protein-DNA contacts. The analytical results (curves) are given by Eq. (37) and (40) in SI. The x-axis increases due to faster 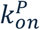, with constant [*P*]_*tot*_. c) Dimerization 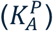, 3D-to-1D binding 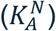, and dimensionality effect (*γ*) strictly enhance the recruitment fraction. Due to target binding, more than 60% of protein monomers are already bound on DNA. The analytical results (curves) are given by Eq (43) in SI (monomer) and Eq. (47) in SI (irreversible dimer). d) One dimer in environment *DNA* + *targ* can only bind with one target and the target-bound protein is stabilized at the cost of the partner finding a target.

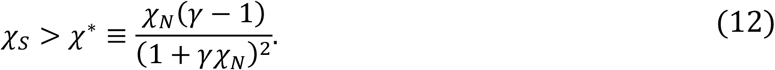

In this limit, the target occupancy is dominated by direct target binding, with minimal impact from nonspecific contacts and 1D sliding, *γ*χ_*N*_. In the other limit, when χ_*S*_ < χ^*^, monomers have low target occupancy so forming dimers improves target binding via nonspecific contacts (Fig. 3a). These same trends continue even for higher protein concentrations ([P]_tot_∼[S]_tot_) in numerical simulations (Fig S7), although the transition point is not as simple as Eq. (12).

Dimerization also may enhance or impair the mean dwell time, *τ*^DNA+targ^, which is quite sensitive to dimer lifetime, 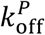. Irreversible dimers 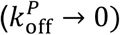 always enhance dwell times. To characterize reversible dimers, we first note that the dwell time of a monomer on a DNA target, 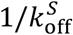, is lengthened by the adjacent nonspecific DNA, 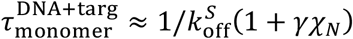, assuming specific binding is much more frequent than nonspecific binding (see SVI.C in SI). With the addition of dimerization, the dwell time is impaired relative to the monomer if the co-localized protein has transient dimer binding, or when 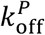 is too fast. This critical rate threshold can be approximated under the assumption of χ_*S*_ ≫ χ_*N*_ to predict impaired dwell times when (see SVII.B in SI):

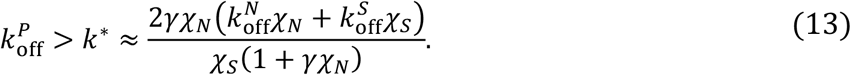

As *k*^*^ shrinks, dimerization therefore becomes more likely to impair dwell times; this occurs for stronger target partitioning χ_*S*_, since *∂k*^*^/*∂*χ_*S*_ < 0. In this regime, like occupancy, the monomer dwell times are dominated by direct target binding and are thus long. When dwell time is controlled by slow target bound states, dimers with transient protein-protein bonds impair lifetimes. In contrast, the larger *k*^*^ becomes, the more likely dimerization will enhance dwell time 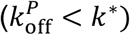. Towards this end, we can show that increases in *γ* will always increase *k*^*^ (*∂k*^*^/*∂γ* > 0), as will faster DNA dissociation, either 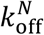 or 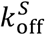. When 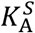 and 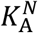 are constants, faster 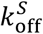 or 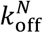 mean faster 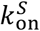 or 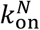, respectively, which helps the co-localized proteins bind with DNA. Finally, the recruitment fraction of proteins to DNA is always higher with dimers than monomers (see Fig. S7, SVII.C in SI). The relative gain achieved due to dimerization is higher for weaker nonspecific interactions (Fig 3c).

### Clustered targets cooperate with dimerization to offer dramatic increases in protein-DNA stability

When two targets are spatially adjacent, a dimer can bind both targets, and thus dimerization and target binding mutually enhance each other (Fig. 4c). A dimer that recruits to a target site through one protein (*PPS*_2_) co-localizes the other protein for rapid association to the second target site (*PPS*_2_ → *PSPS*). These fully-target-bound dimers are very stable since their dissociation off the target requires breaking two bonds (*P*-*S* and *P*-*P*), significantly increasing target occupancy (Fig. 4a). We can also show that recruitment to DNA always increases, 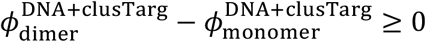 (see SVII.C in SI). Dimerization almost always increases dwell time as well (Fig. 4b). We can calculate the critical rate *k*^*^ for dimer lifetime 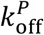 such that dimerization *increases* the dwell time if (see Eq. 55’ in SI):

**Figure 4.**
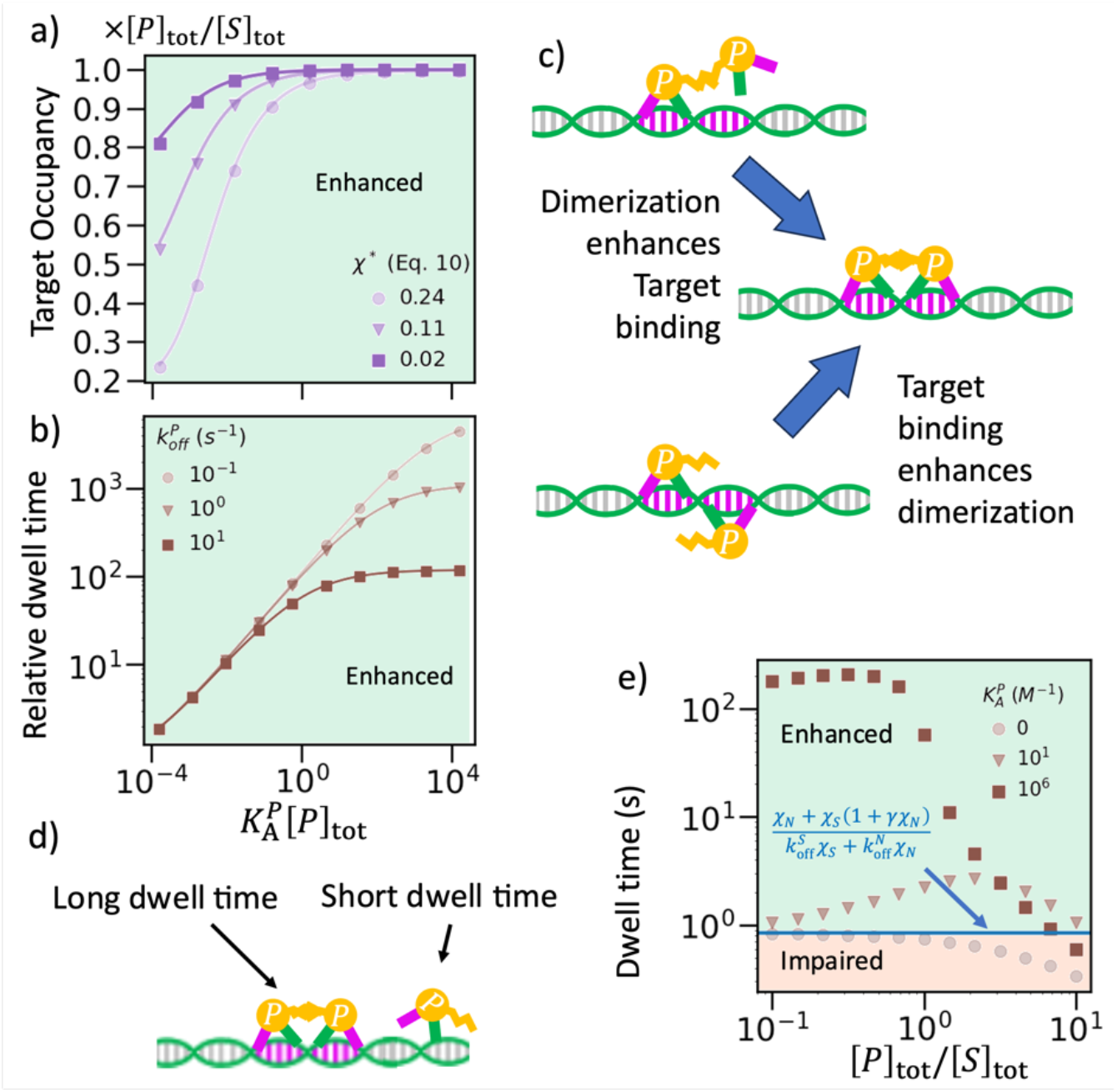
Emergent cooperativity between dimers and clustered targets enhances target occupancy and nearly always increases dwell time. In this figure, [*P*]_*tot*_ = 0.1[*S*]_*tot*_ ([*P*]_*tot*_ = [*S*]_*tot*_ has the same trend, see Fig S8) and numerical results (points) are obtained by solving ODEs (see SXII.A) a) The target occupancy always benefits from dimerization since it no longer sequesters proteins from targets. Theoretical results (curves) are given by Eq. (50) in SI. Occupancy is here again relative to the maximum *ϕ*_*MAX*_ = [*P*]_*tot*_/[*S*]_*tot*_, and the x-axis is increased via 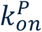. b) The relative dwell time 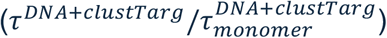 benefits from dimerization. Because of the positive feedback between dimerization and target binding, 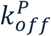 must be very fast before dimerization acts to impair the dwell time in this model. Theoretical results (curves) are given by Eq. (48) and (41) in SI. c) Dimerization and target binding enhance each other to produce an effective positive feedback arising strictly from the co-localization. d) Once all target sites are occupied, excess proteins can only bind DNA nonspecifically. These excess proteins have short dwell times, which reduces the population-averaged dwell time. e) The absolute dwell time is compared between monomers (tan circles), weak dimers (brown triangles) and strong dimers (dark brown squares), as the ratio of proteins-to-targets increase, showing nonmonotonic behavior for dimers. The blue line (Eq. (48) in SI) is the theoretical dwell time of monomers when DNA sites are in excess. Monomer dwell times drop below this theoretical approximation at high ratios of proteins-to-targets because they run out of specific targets and bind nonspecifically.

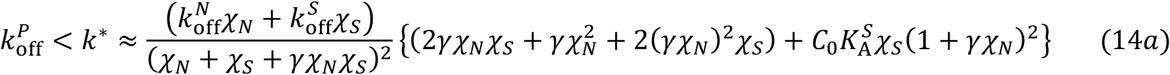

Assuming χ_*S*_ ≫ χ_*N*_ and 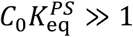, we have,

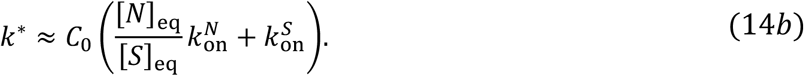

Since *C*_0_ = 1 M and physiologically we estimate 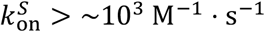, we have a lower bound that 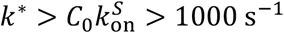. We see that *k*^*^ is already quite high, and nonspecific binding only makes it higher. It requires therefore a very short-lived protein-protein interaction to see 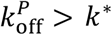, meaning that only in rare cases will dimerization impair dwell time.

For this clustered target environment, we assessed the enhancement strength as the ratio of proteins relative to targets ([*P*]_tot_/[*S*]_tot_) increases from ≪ 1 to 10. With more proteins, the target occupancy monotonically increases (Fig. S8d). However, the dwell time is non-monotonic for dimers. For monomers, excess proteins can only reduce the mean dwell time, as binding nonspecifically introduces short-lived contacts with the DNA (Fig. 4e). In contrast, mean dwell time for dimers initially benefits from increasing protein concentration as dimers helps stabilize target binding, but will ultimately decrease with excess proteins for the same reason as monomers (Fig. 4e). Once proteins are in large excess of targets (>10x), dimerization provides fairly negligible increases to dwell times relative to monomers. The dwell time maxima corresponds with the same intermediate ratio of [*P*]_tot_/[*S*]_tot_ where target occupancy achieves its maximal *gain* and not maximal *occupancy* (Fig. S8d in SI), illustrating the trade-off between increasing protein occupancy on targets versus increasing nonspecifically bound proteins.

### Dimerization generates selectivity of proteins for clustered targets on DNA

We here combine two DNA microenvironments, one with a pair of clustered targets and one with a pair of separated targets, to calculate the spatial selectivity of proteins in this inhomogeneous environment theoretically (see SXI in SI) and compare with spatial stochastic simulations (Fig. 5a). We quantify the selectivity for environment A vs B as the ratio of recruitment fractions to each DNA segment, 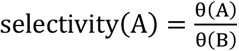, which is > 1 when proteins select for *A*. By keeping all parameters identical in both environments except the separation between two targets, we see that monomers show no selectivity for *A* clustered vs B separated targets, in theory or explicit spatial simulations (Fig. 5c). For even weak dimers, however, with 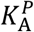 in mM^−1^ scale, we see a dramatic increase in protein selectivity for the clustered targets (Fig. 5bc). The selectivity increases as expected for stronger dimers (Fig. 5b). However, these simple monotonic trends are not retained when protein copies match target copies, or 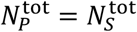. In this case, once the clustered target is occupied, proteins/dimers remaining in solution will now bind the separated targets. While there is still selectivity for the clustered targets, it is reduced with stronger dimerization as once θ(A) → 1 on the clustered segment, dimerization can now only increase θ(B) (Fig. 5b). Lastly, we note that the particle-based stochastic NERDSS simulations[45] show a similar trend in selectivity to the theory, but with some deviations particularly for strong dimers (Fig. 5b). This is primarily driven by the discrete individual molecules tracked in the NERDSS simulations, whereas in the continuum analytical theory, we allow fractional distributions of proteins throughout the system. By explicitly capturing spatial heterogeneity and diffusion, the NERDSS simulations show an even stronger switch-like behavior in selectivity for clustered sites as dimerization strength increases above ∼ 10^2^ M^−1^.

**Figure 55.**
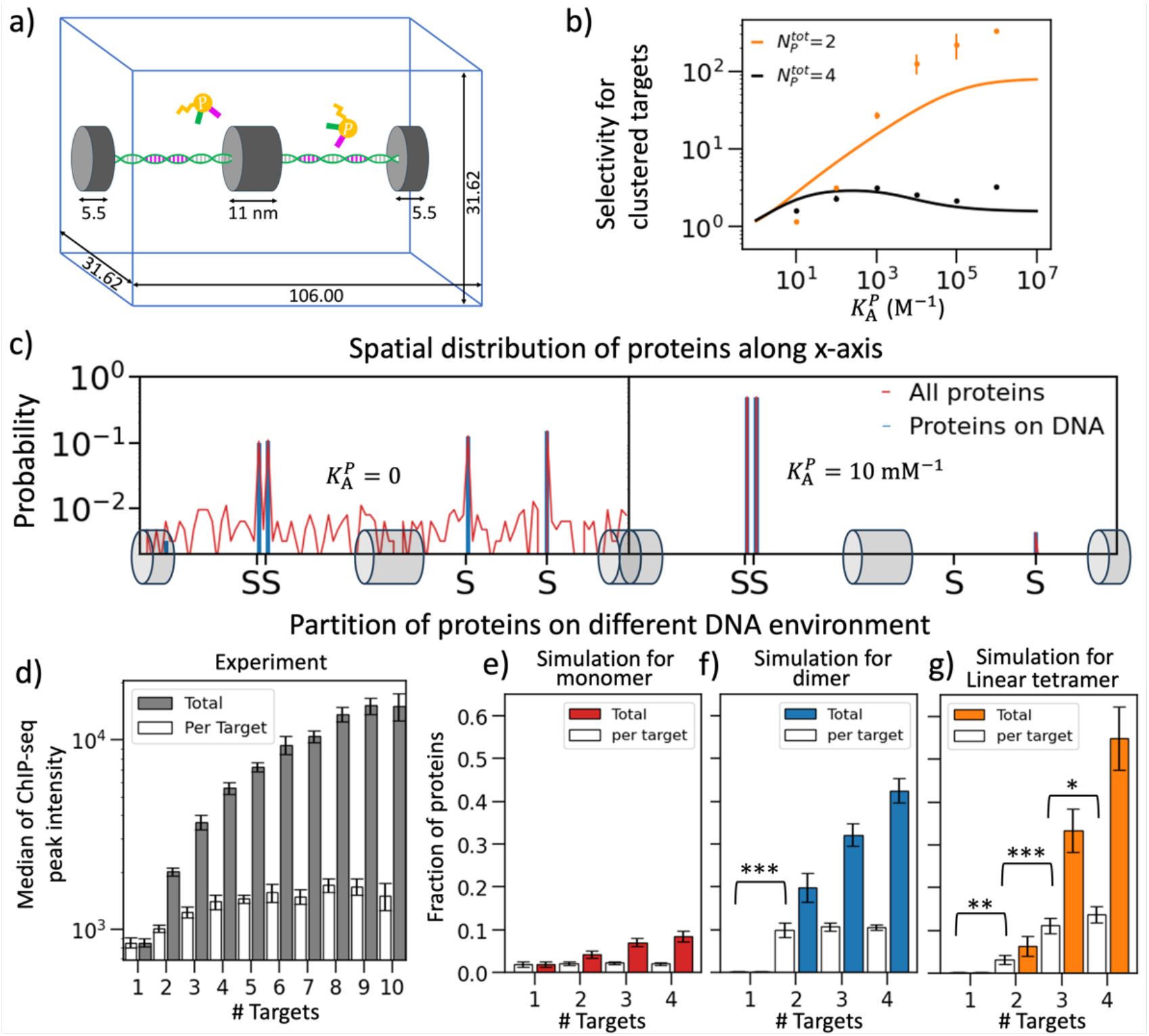
Dimers and oligomers select for targets based on their spatial separation. a) System set-up for stochastic particle-based NERDSS simulations to quantify selectivity of proteins for two clustered targets (left side of nucleosome) vs two separated targets (right side). b) As dimerization strength increases, two proteins become highly selective for clustered targets (orange), while 4 proteins have weaker selectivity (black). Points from NERDSS (see SXII.E in SI), and lines are calculated using an ODE model with 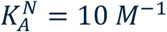 and 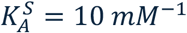 (see SI section SXI). c) Averaged spatial distribution of proteins from equilibrated NERDSS simulations (SXII.E in SI) with 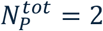 with no dimerization 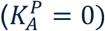 on the left, showing no selection for clustered targets, and with weak dimers on the right with high selectivity 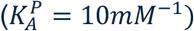. Blue histograms indicate proteins positioned on DNA, whereas red curves are all proteins (3D and 1D). Both are normalized by 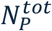. d) The ChIP-seq peaks of GAF [44] to sequences containing more targets have a higher intensity (dark gray bars). Targets (GAGAG motifs) are counted as unique when they do not overlap. The per-target intensity also increases, demonstrating a nonlinear dependence on number of targets (white bars). The standard errors are estimated by 100 bootstraps of ChIP-seq peaks at each target number. e), f), g) NERDSS simulation results for proteins binding to DNA where one segment has a single target, one segment has 2 targets, one segment has 3 targets, and one segment has 4 targets (see SXII.F in SI) e) Monomers do not show enhanced selectivity for targets arrayed in clusters (white bars) f) Dimerization helps proteins select DNA segments with a cluster of 2 or more targets (white bars) g) Tetramerization helps proteins select DNA with a cluster of 4 targets more strongly than smaller clusters. Diffusion constants of protein monomers are *D*_1*D*_ = 0.6 *nm*^2^/*μs* and 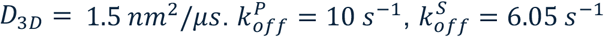, and 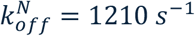.

### Simulation results reproduce ChIP-seq data on oligomer selectivity for clustered targets

The GAF protein forms higher-order oligomers of up to 6-mers and possibly larger [20, 51, 52] and has thousands of targets in the genome (Fig S10). *In vivo*, GAF occupancy as measured via ChIP-seq [44] is higher at clustered targets [20] (Fig. 5d, SI). Through spatial simulations, we therefore challenged our proteins to now select between targets organized as a single target, cluster of two targets, or four-in-a-row clusters (Fig. 5d). We inevitably expect more proteins at the 4-site cluster compared to the 1-site cluster, and indeed if the proteins do not dimerize, their occupancy is increased by exactly 4-fold (Fig. 5e). More generally, the occupancy *ϕ*_*n*_ on a cluster of *n* targets is 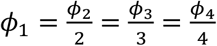 for monomers (Fig. S9). However, when we allow dimerization, we now see effectively no proteins binding to the 1-site cluster, and an increasing fraction from 2 to 4 sites. Normalizing by *n*, the dimer is clearly selective for a cluster of size 2 over 1, whereas *n* = 4 only increases due to a higher target density, 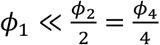, producing a plateau (Fig. 5f). This result implies that proteins that form a tetrameric oligomer would be maximally selective for a cluster of 4 targets, and indeed this is what we recover with our model, 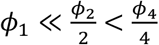 (Fig. 5g, Fig. S9).

These model results are consistent with the GAF ChIP-seq results, as GAF occupancy is higher in genome regions with clusters of larger *n* target sites, even after normalizing the intensity by *n*, i.e. 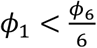 (Fig. 5d). This normalized intensity even shows a plateau at ∼*n* = 6 for GAF 6-mers. This result is only consistent with the *in-silico* oligomerization experiment and not the monomer behavior. We also observe the *in-silico* selectivity appears to be higher than GAF (Fig. 5fg), but this makes sense when considering the frequency of distinct clusters. Our simulated clusters (*n* =1,2,3 or 4 sites per DNA segment) were present at equal amounts, to simplify the simulation set-up (Fig. S10). In contrast, for GAF targets, cluster abundances are highly non-uniform (Fig. S10-d,e); the majority of target sequences appear alone (*n* = 1), and more than 90% are present at *n* < 3 [53]. The numerous sequences with small target clusters compete for proteins and impair the overall selectivity for sequences with large clusters but do not eliminate it, as we verify using simulations (Fig. S10fg). Hence the GAF shows remarkable selectivity for these large but infrequent clusters over single-target sites.

## DISCUSSION

Our results show that dimerization between DNA-binding proteins will only universally enhance DNA target occupancy and selectivity when the target sites are clustered to the same or higher degree than the protein dimer. Thus, although dimerization always increases the recruitment fraction (SVII.C in SI), driving more proteins onto DNA from solution, the intuition that these proteins are more effective at occupying DNA targets oversimplifies the role of population dynamics (Fig. 6). A critical control parameter for all models is the strength of 1D partitioning to the nonspecific DNA, or 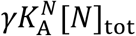. Without nonspecific binding, all trends are relatively straightforward—for isolated targets, dimerization impairs, and for clustered targets, it enhances (SVIII in SI). With nonspecific binding, a long-lived dimer (slow 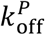) that binds an isolated single target will enhance its own dwell time on DNA via nonspecific interactions, but by sequestering proteins from distant targets, the population can report a reduction in the mean dwell time (Fig. 6). A short-lived dimer (fast 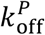) can also reduce the mean dwell time by recruiting proteins to DNA via transient protein interactions. Mutations that eliminate protein-protein interactions [20] can thus have unpredictable effects on dwell time. For isolated targets, dimerization can also simultaneously impair dwell time while still enhancing occupancy (black star in Fig. 6). While this may seem counterintuitive, it points to distinct driving factors for these two metrics; dimers can increase flux onto target DNA (higher effective on-rate), despite shortening the mean lifetime of the population across DNA (higher effective off-rate). Such isolated target motifs are common in the genome for proteins like GAF [53], and with a sufficiently transient protein-protein bond lifetime and moderate target binding, dimerizing proteins can occupy these isolated motifs more frequently but for short intervals (Fig. 6), supporting ‘trials’ of binding that may help ultimately identify clustered targets.

**Figure 6 6.**
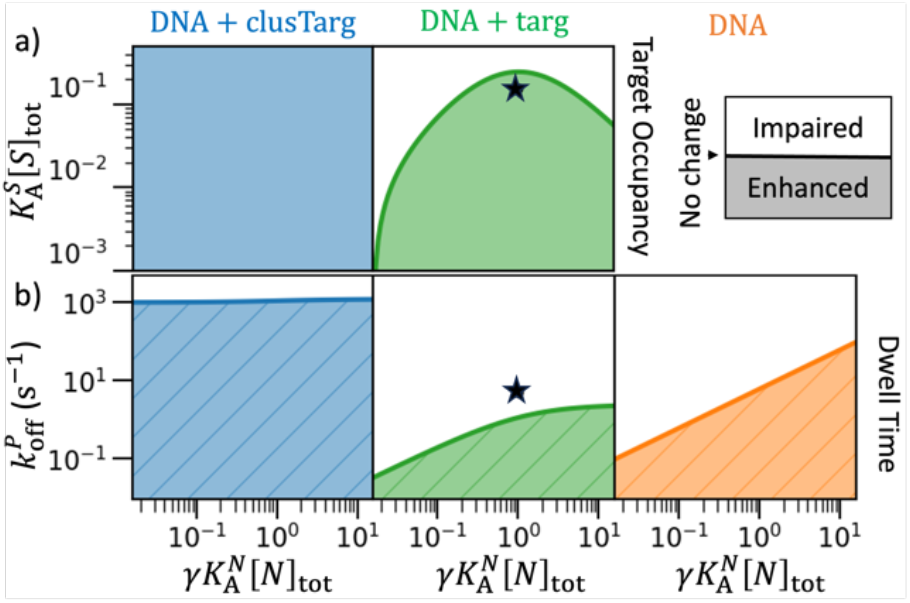
Phase diagrams summarize when dimerization enhances or impairs dwell times and occupancy relative to monomers. The solid lines in all plots represent no change between dimers and monomers. Colored regions where dimerization leads to enhancement (legend). Left column (blue) is for the *DNA* + *clusTarg* system, middle column (green) is for the *DNA* + *targ* system, right column (orange) is for bare DNA. DNA*DNA*. The x-axes measure response to increasing nonspecific binding and dimensional enhancement. a) Target occupancy is measured as the y-axis increases propensity for specific binding. Orange system has no targets. For *DNA* + *targ* (green), the green line is not monotonic with stronger 1D partitioning, illustrating sequestration to nonspecific DNA (green line Eq. 12)). With clustered targets (blue), stronger dimerization always increases target occupancy. The black stars in a) and b) label the same condition, illustrating how occupancy can increase despite impaired dwell time. b) Dwell time is shown as rate of protein-protein dissociation, 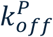 increases. In environment *DNA* + *clusTarg* (blue), only very short dimer lifetimes impair occupancy. In environment *DNA* + *targ* (green), dimerization frequently impairs the dwell time. For bare *DNA*, dimerization also impairs the dwell time when 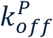 is fast (orange). Eq.(9) for orange edge, Eq. (13) for green edge, and Eq. (14) for blue edge). Other parameters in SI.

By extending beyond previous quantitative models for predicting DNA dwell time and occupancy via the explicit addition of dimerization, we significantly expand the state-space (up to 16 states) beyond the monomer bound to specific or nonspecific DNA (3-4 states) [22, 23], providing a new framework for *quantitatively* interpreting *in vitro* DNA binding experiments on multi-domain proteins [54]. With a thermodynamically consistent framework, our steady-state equations for dwell times and occupancies are relatively interpretable and transferrable binding parameters (e.g. 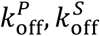) can be extracted from data despite the convolution of both protein and DNA contributions. Our application of the model to the GAF system has integrated out more accurate representation of the *in vivo* environment, where the genome organization is both structured and dynamic[55-57], mechanics of DNA contributes to affinities[58, 59], nonspecific regions can induce variable association strengths[60], and macromolecular interactions induce higher-order assemblies[61] or liquid-like droplets[52]. Further, we exclude energy-consuming processes that, like dimerization, can non-monotonically enhance the response to concentration variations[62]. Nonetheless, because these additional sources of variability exist throughout the genome, the data and our model clearly demonstrate that even weak protein-protein interactions can dramatically enhance selectivity for clustered targets, which is particularly notable when they are infrequent in the genome as is the case for GAF (Fig S10). This produces what is effectively a spectrum of affinities for different regions of the genome [63] that is not a linear function of the target cluster size but rather a nonlinear function of (also) the protein-protein affinity. Predicting this emergent spectrum for oligomers from the fundamental pairwise interactions, including both 3D and 1D association, is a key outcome of our model (Fig. 5).

For generalized transcription factors with widespread target sites, one expects selectivity for the sequences of highly-expressed genes to be higher to enhance transcription[64], and that should fundamentally depend on collectively stronger affinities for DNA emerging not just from sequence recognition but via protein self-assembly[17, 65]. Dimerization is then an essential building block for identifying the correct sequence for transcription with the right frequency, but our work shows that it must be tuned to help rather than hinder target binding. This tunability can be encoded in nonspecific interactions or the 1D protein-protein binding rates, particularly the *h*^2^ factor from *γ*, in addition to the 3D rates. While we here considered *h*^2^ to span 1000-fold variation, the significance of this 3D-to-1D binding parameter in controlling DNA interactions urges more careful quantification, as it has not been measured, and similar to membrane-associated interactions[66], allosteric effects could make assembly much more (or less) favorable in 1D vs 3D [35]. With expansions of this model to include heterogeneity in protein interactions and diverse components, we predict stoichiometry then provides further nonlinear tunability for which genes are ‘switched on’ for transcription either constitutively or in response to stimulus. This increasing design space for parameters will support more effective classification of DNA sequences, like a deeper neural network, with assemblies of diverse DNA-binding domains improving selectivity for longer sequence tracts of distinct motif clusters. The models used here will help predict how competition and cooperation between binding partners control selectivity as it varies nonlinearly with concentrations, binding rates, and spatial localization throughout the genome, which is necessary to understand proper transcriptional control [67].

## Supplemental Information

Supplementary Information contains SI Text, 10 SI Figures and 7 SI Tables. Dataset S1 is a spreadsheet enumerating all parameters used for plotted data, and code is in the github repo: https://github.com/sangmk/dimerEnhanceProteinDNA

## Acknowledgements

MEJ gratefully acknowledges support from an NIH U01DK127432 Award. We thank Profs Carl Wu, Taekjip Ha, and Yaojun Zhang, along with Dr. Xiaona Tang for helpful discussions on the project.

