## Supplemental Information for "Mechanisms of enhanced or impaired DNA target selectivity driven by protein dimerization"

**This PDF file includes:**

Supporting text

Tables S1 to S7

Figures S1 to S10

Legends for Dataset S1

Legends for Software S1

SI references

**Other supporting materials for this manuscript include the following:**

Dataset S1 (“Parameters_rateEquations.xlsx”)

Software S1 (github repo)

### Supplemental Methods

#### SI. Background on the dimensionality factor and its emergence in non-spatial ODE models.

To solve for the model illustrated in Fig. 1b nonspatially, such that all species can be converted between one another, we must choose what units we will use (or copy numbers) for concentrations. The association constant is in general a ratio of the forward and reverse rates: $K_{\mathrm{A}}^{S} = \frac{k_{\mathrm{on}}^{S}}{k_{\mathrm{off}}^{S}}$, where we leave out the dimension label for simplicity for all 3D reactions. We report all association constants with units/dimensions (e.g. $V$ in 3D) so the standard state does not need to be inferred. The binding equilibrium for a species in 3D concentration, for example, $[PS] = N_{PS}/V$ where $N_{PS}$ are copy numbers of $PS$, is given by $\frac{N_{PS}V}{N_P N_S} = \frac{k_{\mathrm{on}}^{S}}{k_{\mathrm{off}}^{S}}$. In these 3D units, the association rate has units of $V/\mathrm{time}$ and dissociation has units of $1/\mathrm{time}$. For the 1D reactions, however, we have that,

$$\frac{N_{PSN}L}{N_{PN}N_S} = \frac{k_{\mathrm{on}}^{1D,S}}{k_{\mathrm{off}}^{1D,S}} \tag{1}$$

Where association and dissociation rates have units of $L/\mathrm{time}$ and $1/\mathrm{time}$ respectively. To estimate 1D rates for a binding pair given a known 3D rate, one can consider the proteins as switching from diffusion to target recognition[1], with the protein re-orienting on the target sequence[2, 3]. To maintain generality, we note a corresponding characterization of 3D vs 2D rates and affinities using theory[4], simulation[5, 6], and experiment[7, 8]. If the change in equilibrium affinities results purely from entropic restrictions to configurations [7, 8], then the corresponding length scale $h$ is at the molecular or nanometer scale. It is important to note that it is not true that $\frac{N_{PSN}V}{N_{PN}N_S} = \frac{k_{on}^{3D,S}}{k_{\mathrm{off}}^{3D,S}}$, as this relation asserts that the copies are performing a 3D search in a volume $V$. Instead, to put the Eq. SI.1 in volume units, they must be re-scaled by multiplying both sides by the dimensionality factor (Eq. 1 of the main text):

$$\gamma = \frac{V}{Lh^2} \tag{2}$$

The rates are related to one another via,

$$k_{\mathrm{on}}^{1D} = \frac{k_{\mathrm{on}}^{3D}}{h'^2} \tag{3}$$

$$k_b^{1D} = ck_b^{3D} \tag{3'}$$

where $h'^2$ has units of area, and the off-rates can be distinct in 3D vs 1D by a scalar, $c$. The association constants are thus related by $h^2 = ch'^2$,

$$K_A^{1D} = \frac{K_A^{3D}}{h^2} \tag{4}$$

By multiplying both sides in Eq. 1 by $V/L$ and using Eq. 2, 3, and 4, we find:

$$\frac{N_{PSN}V}{N_{PN}N_S} = \frac{Vk_{\mathrm{on}}^{3D,S}/h'^2}{Lck_{\mathrm{off}}^{3D,S}} = \frac{Vk_{\mathrm{on}}^{3D,S}}{Lh^2k_{\mathrm{off}}^{3D,S}} = \gamma K_A^{3D,S} \tag{5}$$

In this way, we can see that the bimolecular equilibria between the 1D species is related to the 3D association constant and the 3D concentrations via the factor $\gamma$.

Equivalently, if we consider the reaction rate equation for $PN + S \rightleftharpoons PSN$, which is in 1D,

$$\frac{dN_{PSN}}{Ldt} = -\frac{N_{PSN}k_{\mathrm{off}}^{1D,S}}{L} + \frac{k_{\mathrm{on}}^{1D,S}N_{PN}N_S}{L^2} \tag{6}$$

If we plug in Eq. 3 and multiply both sides of Eq. 6 by $L/V$, we get $\frac{dN_{PSN}}{Vdt} = -\frac{N_{PSN}ck_{\mathrm{off}}^{3D,S}}{V} + \frac{ck_{\mathrm{on}}^{3D,S}N_{PN}N_S}{Lh^2V}$. Finally, this equation will be solved in fully 3D units by multiplying the 3rd term by $V/V$ and using Eq. 2, giving,

$$\frac{dN_{PSN}}{Vdt} = -\frac{N_{PSN}ck_{\mathrm{off}}^{3D,S}}{V} + \frac{\gamma ck_{\mathrm{on}}^{3D,S}N_{PN}N_S}{V^2}, \tag{6'}$$

which is exactly the equation in volume units, with the association step multiplied by the $\gamma$, and we allow a rescaling between 3D and 1D of the dissociation rate as well, noting that the $\gamma$ used here is based on the association constant Eq. 4, not the forward rate, hence the factor of $c$ also showing up in the association rate.

To track the time-dependent dynamics of proteins with DNA using the well-mixed approximation, we must solve the coupled nonlinear ODEs for the 6 volume species:

$$\begin{aligned}
\frac{d[P]}{dt} &= -k_{\mathrm{on}}^{S}[P][S] + k_{\mathrm{off}}^{S}[PS] - k_{\mathrm{on}}^{N}[P][N] + k_{\mathrm{off}}^{N}[PN] \\
\frac{d[S]}{dt} &= -k_{\mathrm{on}}^{S}[P][S] + k_{\mathrm{off}}^{S}[PS] - \gamma c_2 k_{\mathrm{on}}^{S}[PN][S] + c_2 k_{\mathrm{off}}^{S}[PSN] \\
\frac{d[N]}{dt} &= -k_{\mathrm{on}}^{N}[P][N] + k_{\mathrm{off}}^{N}[PN] - \gamma c k_{\mathrm{on}}^{N}[PS][N] + c k_{\mathrm{off}}^{N}[PSN] \\
\frac{d[PS]}{dt} &= k_{on}^{PS}[P][S] - k_{\mathrm{off}}^{S}[PS] - \gamma c k_{\mathrm{on}}^{N}[PS][N] + c k_{\mathrm{off}}^{N}[PSN] \\
\frac{d[PN]}{dt} &= k_{\mathrm{on}}^{N}[P][N] - k_{\mathrm{off}}^{N}[PN] - \gamma c_2 k_{\mathrm{on}}^{S}[PN][S] + c_2 k_{\mathrm{off}}^{S}[PSN] \\
\frac{d[PSN]}{dt} &= \gamma c k_{\mathrm{on}}^{N}[PS][N] - c k_{\mathrm{off}}^{N}[PSN] + \gamma c_2 k_{\mathrm{on}}^{S}[PN][S] - c_2 k_{\mathrm{off}}^{S}[PSN]
\end{aligned} \tag{7}$$

where $c$ and $c_2$ are the 3D-to-1D scalers for $k_{\text{off}}^{N}$ and $k_{\text{off}}^{S}$, respectively. Because of the thermodynamic cycles in this model (Fig. 1 in main text), we note that the magnitude of $h^2$ must be the same for both 1D binding events, although the on/off rates can be uniformly scaled from one reaction to the other. For simplicity, our models assume $c = c_2 = 1$, or the off rates are the same in 1D and 3D.

#### SII. Role of changes to $V/L$ on the system behavior

Increases in $\gamma$ will follow from either a decrease in $h^2$ or an increase in the ratio of $V/L$. The effects are not equivalent; $h^2$ is independent of all other system parameters, but $V/L$ is not, so for this section only we assume $h^2$ is fixed. A somewhat subtle point is that increases to the volume, while increasing $\gamma$, must also change either relative protein to DNA site copy numbers, or relative protein to DNA site concentrations. To illustrate, we consider two possible outcomes where $V$ increases but $L$ is unchanged. I) We increase the volume but assume the protein concentration stays the same. We therefore have an increase in protein copy numbers $P$, but without a corresponding increase in nonspecific sites $N$, since $N = \frac{n_{\text{non}}}{l} L$, and the density of nonspecific sites, $\frac{n_{\text{non}}}{l}$, is physically constant. Thus, the concentration of nonspecific sites has diluted via the volume increase ($[P]_{\text{tot}}$ is the same, but $[N]_{\text{tot}}$ decreases). II) We increase the volume and assume all copy numbers remained the same. We now retain the same ratio of $P/N$, but both species are diluted (both $[P]_{\text{tot}}$ and $[N]_{\text{tot}}$ decrease).

#### SIII. Background and definitions of two-state metrics for dwell time given our microkinetic models

All our models contain multiple protein-DNA bound states (e.g. $PN$, $PS$, and $PSN$) and unbound states (e.g. $P$ and $PP$). To define convenient metrics that compare between models and capture what is typically measured in experiment, we can map these many microkinetic states to a two-state model. In our primary two-state model, we compare proteins associate to DNA (including both specifically and nonspecifically bound proteins) vs proteins dissociate from DNA. We are interested in the timescales and occupancies at an equilibrium steady state, where the flux in and out of each pairwise binding reaction is equivalent, e.g. $k_{\text{on}}^{N}[P]_{\text{eq}}[N]_{\text{eq}} = k_{\text{off}}^{N}[PN]_{\text{eq}}$. The sum over sets of pairwise reactions is thus also equal. To maintain generality, here we denote proteins in unbound states as $A_i$ and proteins in bound states as $AB_j$. Each $A_i$ or $AB_j$ state has proteins with $a_{A_i}$ or $a_{AB_j}$ stoichiometry, respectively. The proteins in $\{A_i\}$ can transit to $\{AB_j\}$ by binding with $B_k$ which can be DNA sites (e.g. $S$ and $N$) or proteins on DNA (e.g. $PN$). Then, we can express the equilibrium flux as,

$$\sum_{r=1}^{R} a_{A_r} k_{\text{on}}^{r} [A_r][B_r] = \sum_{r=1}^{R} a_{AB_r} k_{\text{off}}^{r} [AB_r] \tag{7}$$

where $R$ sums over sets of pairwise reactions $A_r + B_r \rightleftharpoons AB_r$ that include transitions between the two states. Any reaction where the association constant $K_{\text{A}}^{r}$ (or association rate $k_{\text{on}}^{r}$) is zero must be left out of the sums.

We can further define an effective two-state association constant and corresponding two-state rate constants,

$$\frac{k_{\text{on,2state}}}{k_{\text{off,2state}}} = K_{A,\text{2state}} = \frac{\sum_{j=1}^{Nb} a_{AB_j}[AB_j]}{\sum_{r=1}^{R} a_{A_r}[A_r][B_r]} \tag{8}$$

The sum $Nb$ in the numerator is over all molecules in the bound state. The summation in the denominator over $R$ involves pair-wise reactions directly transitioning from unbound state to bound state. We note that these pairwise reactions must have a non-zero affinity (or $k_{\text{on}}$), otherwise they should be excluded from the sum. The sums in the numerator and denominator need not be over the same reactions. For example, the state $PNPN$ is part of the bound states in the numerator, but its formation does not contribute to the denominator because it cannot be directly transitioned into from the unbound state in a single step. The dwell time in the bound state of this two-state model is the inverse of this effective off-rate,

$$\tau = \frac{1}{k_{\text{off,2state}}} = \frac{\sum_{j=1}^{Nb} a_{AB_j}[AB_j]}{k_{\text{on,2state}} \sum_{r=1}^{R} a_{A_r}[A_r][B_r]}. \tag{9}$$

However, the denominator here requires a definition of the 2-state on-rate. A self-consistent definition for the on-rate is as a weighted average over all pairwise flux into the bound state,

$$k_{\text{on,2state}} = \frac{\sum_{r=1}^{R} k_{\text{on}}^{r} a_{A_r}[A_r][B_r]}{\sum_{r=1}^{R} a_{A_r}[A_r][B_r]}, \tag{10}$$

and thus,

$$k_{\text{off,2state}} = \frac{\sum_{r=1}^{R} k_{\text{off}}^{r} a_{A_r}[AB_r]}{\sum_{j=1}^{Nb} a_{AB_j}[AB_j]}. \tag{11.b}$$

Which recovers $K_{\text{A,2state}} = k_{\text{on,2state}}/k_{\text{off,2state}}$ in Eq. 9 when using Eq. 8 with the same $R$ set in both $k_{\text{off,2state}}$ and $k_{\text{on,2state}}$ . From Eq. 10, we now obtain for the mean dwell time,

$$\tau = \frac{\sum_{j=1}^{Nb} a_{AB_j}[AB_j]}{\sum_{r=1}^{R} k_{\text{on}}^{r} a_{A_r}[A_r][B_r]} \tag{11}$$

We note one technical point that when the two-state definition separates proteins in a free state (in solution) and a DNA bound state (via binding $[N]$ or $[S]$ or a protein on DNA), one must take care in taking the limiting cases for $k_{\text{on,2state}}$ or $K_{\text{A,2state}}$. They both reach the correct limiting behavior when $[N] \to 0$ or $[S] \to 0$, that $k_{\text{on,2state}}$ becomes $k_{\text{on}}^{S}$ or $k_{\text{on}}^{N}$, respectively, and $K_{\text{A,2state}}$ becomes $K_{\text{A}}^{S}$ or $K_{\text{A}}^{N}$, as expected for a monomer binding (no dimers). However, taking the limit of $K_{\text{A}}^{N} = 0$ is not the same as setting $[N] = 0$ if the denominator assumes that $P + N$ is still a valid path to the DNA bound state. Hence the point we made above, that if one of the DNA binding affinities is set to zero, the sums over $R$ should exclude reaction paths that have a $K_{\text{A}}^{X} = 0$ or a $k_{\text{on}}^{X} = 0$. Perhaps more simply, if $K_{\text{A}}^{N} = 0$, also set $[N] = 0$, and similarly for $K_{\text{A}}^{S}$. The dwell time is robustly accurate in both limits because of the appearance of $k_{\text{on}}^{X}$ in the $R$ sums.

#### SIV. Exact definitions for our models with protein dimerization in all 3 environments: 1.DNA, 2.DNA+targ, and 3.DNA+clusTarg

##### *SIV.A. Dwell time*

These formulas are used to generate the exact dwell times from the numerical simulations at equilibrium, shown in our plots as points. Here we consider dwell times when proteins can form dimers and use superscripts to identify our three different model environments, we have $\tau^{\text{DNA}}$, $\tau^{\text{DNA+targ}}$, and $\tau^{\text{DNA+clusTarg}}$.

For $\tau^{\text{DNA}}$, the sums in Eq. 12 are numerator:

$$\sum_{j=1}^{Nb} a_{AB_j}[AB_j] = [PN]_{\text{eq}} + 2\big([PPN]_{\text{eq}} + [PNPN]_{\text{eq}}\big), \tag{12}$$

And denominator:

$$\sum_{r=1}^{R} k_{\text{on}}^{r} a_{A_r}[A_r][B_r] = k_{\text{on}}^{N}[N]_{\text{eq}}\big([P]_{\text{eq}} + 2\cdot 2[PP]_{\text{eq}}\big) + 2k_{\text{on}}^{P}[PN]_{\text{eq}}[P]_{\text{eq}}. \tag{13}$$

For $\tau^{\text{DNA+targ}}$, the sums in Eq. 12 are numerator:

$$\begin{aligned}\sum_{j=1}^{Nb} a_{AB_j}[AB_j] = \big([PN]_{\text{eq}} + [PS]_{\text{eq}} + [PSN]_{\text{eq}}\big) \\ +2\big([PPN]_{\text{eq}} + [PPS]_{\text{eq}} + [PPSN]_{\text{eq}}\big) \\ +2\big([PNPN]_{\text{eq}} + [PSPN]_{\text{eq}} + [PNPSN]_{\text{eq}}\big),\end{aligned} \tag{14}$$

And denominator:

$$\sum_{r=1}^{R} k_{\mathrm{on}}^{r} a_{A_r}[A_r][B_r] = \left(2\cdot 2[PP]_{\mathrm{eq}} + [P]_{\mathrm{eq}}\right)\left(k_{\mathrm{on}}^{N}[N]_{\mathrm{eq}} + k_{\mathrm{on}}^{S}[S]_{\mathrm{eq}}\right)$$
$$+2[P]_{\mathrm{eq}} k_{\mathrm{on}}^{P}\left([PN]_{\mathrm{eq}} + [PS]_{\mathrm{eq}} + [PSN]_{\mathrm{eq}}\right). \tag{15}$$

For $\tau^{\mathrm{DNA+clusTarg}}$, the sums in Eq. 12 are numerator:

$$\sum_{j=1}^{Nb} a_{AB_j}[AB_j] = \left([PN]_{\mathrm{eq}} + [PS_2]_{\mathrm{eq}} + [PS_2N]_{\mathrm{eq}}\right)$$
$$+2\left([PPN]_{\mathrm{eq}} + [PPS_2]_{\mathrm{eq}}\right)$$
$$+2\left([PPS_2N]_{\mathrm{eq}} + [PS_2PN]_{\mathrm{eq}} + [PNPN]_{\mathrm{eq}} + [PNPS_2N]_{\mathrm{eq}}\right)$$
$$+2\left([PSPS]_{\mathrm{eq}} + [PSPSN]_{\mathrm{eq}} + [PSNPSN]_{\mathrm{eq}}\right), \tag{16}$$

And denominator:

$$\sum_{r=1}^{R} k_{\mathrm{on}}^{r} a_{A_r}[A_r][B_r] = \left(2\cdot 2[PP]_{\mathrm{eq}} + [P]_{\mathrm{eq}}\right)\left(k_{\mathrm{on}}^{N}[N]_{\mathrm{eq}} + 2k_{\mathrm{on}}^{S}[S_2]_{\mathrm{eq}}\right) \tag{17}$$
$$+2[P]_{\mathrm{eq}} k_{\mathrm{on}}^{P}\left([PN]_{\mathrm{eq}} + [PS_2]_{\mathrm{eq}} + [PSN]_{\mathrm{eq}}\right).$$

See section SXII.A for numerically evaluating these equations.

##### *SIV.B. Recruitment fraction*

Recall Eq. 3 in the main text

$$\theta = 1 - \frac{[P]_{\mathrm{eq}} + 2K_{\mathrm{eq}}^{PP}[P]_{\mathrm{eq}}^{2}}{[P]_{\mathrm{tot}}} = \frac{\sum_{i=1}^{\mathrm{Nb}} a_{AB_i}[AB_i]}{[P]_{\mathrm{tot}}}. \tag{18}$$

Although the middle equation is a simple way for calculating the recruitment fraction, to uncover the property of each environment, we need to express the recruitment fraction in the format of the right-hand side of Eq. 19 that is complicated but informative,

For $\theta^{\mathrm{DNA}}$, $\sum_{i=1}^{\mathrm{Nb}} a_{AB_i}[AB_i]$ is given by Eq. 13.

For $\theta^{\mathrm{DNA+targ}}$, $\sum_{i=1}^{\mathrm{Nb}} a_{AB_i}[AB_i]$ is given by Eq. 15.

For $\theta^{\mathrm{DNA+clusTarg}}$, $\sum_{i=1}^{\mathrm{Nb}} a_{AB_i}[AB_i]$ is given by Eq. 17.

See section SXII.A for numerically evaluating these equations.

##### *SIV.C. Target occupancy*

Similarly, although Eq. 4 in the main text gives a simple equation that $\phi = 1 - \frac{[S]_{\text{eq}}}{[S]_{\text{tot}}}$, we can evaluate free $[S]_{\text{eq}}$ from knowing the bound states. For separated targets (environment DNA+targ), we have,

$$\phi^{\text{DNA+targ}} = \frac{[PS]_{\text{eq}} + [PSN]_{\text{eq}} + [PPS]_{\text{eq}} + [PPSN]_{\text{eq}} + [PSPN]_{\text{eq}} + [PNPSN]_{\text{eq}}}{[S]_{\text{tot}}}. \quad (19)$$

For clustered targets (environment DNA+clusTarg), we have,

$$\begin{aligned}\phi^{\text{DNA+clusTarg}} &= \frac{[PS_2]_{\text{eq}} + [PS_2N]_{\text{eq}}}{[S]_{\text{tot}}} \\ &+2\frac{[PPS_2]_{\text{eq}} + [PPS_2N]_{\text{eq}} + [PS_2PN]_{\text{eq}} + [PNPS_2N]_{\text{eq}}}{[S]_{\text{tot}}} \\ &+2\frac{[PSPS]_{\text{eq}} + [PSPSN]_{\text{eq}} + [PSNPSN]_{\text{eq}}}{[S]_{\text{tot}}}\end{aligned} \quad (20)$$

See section SXII.A for numerically evaluating these equations.

### SV. Equilibrium equations, propensity functions, and rate equations for three environments.

#### SV.A. Propensity functions.

We use propensity functions to describe the probability of reactions. These propensity functions are used in Gillespie methods (see SXII.B) to decide which reactions occurs and used to express rate equations in a compact way. For example, for the monomer system shown in Fig 1 of the main text there are 4 reversible reactions, $P + N \leftrightharpoons PN$, $P + S \leftrightharpoons PS$, $PN + S \leftrightharpoons PSN$, and $PS + N \leftrightharpoons PSN$, and 6 species $P$, $N$, $S$, $PN$, $PS$, and $PSN$. The propensities are, $p_{P+N\to PN} = k_{\text{on}}^{N}[P][N]$, $p_{PN\to P+N} = k_{\text{off}}^{N}[PN]$, $p_{P+S\to PS} = k_{\text{on}}^{S}[P][S]$, $p_{PS\to P+S} = k_{\text{off}}^{S}[PS]$, $p_{PN+S\to PSN} = \gamma k_{\text{on}}^{N}[PN][S]$, $p_{PSN\to PN+S} = k_{\text{off}}^{N}[PSN]$, $p_{PS+N\to PSN} = \gamma k_{\text{on}}^{N}[PS][N]$, and $p_{PSN\to PS+N} = k_{\text{off}}^{N}[PSN]$.

Using these propensities, the rate equations can be rewritten as, $\frac{d}{dt}\begin{pmatrix}[P]\\ [N]\\ [S]\\ [PN]\\ [PS]\\ [PSN]\end{pmatrix} =$

$$\begin{pmatrix}-1 & -1 & 0 & 1 & 0 & 0\\ 1 & 1 & 0 & -1 & 0 & 0\\ -1 & 0 & -1 & 0 & 1 & 0\\ 1 & 0 & 1 & 0 & -1 & 0\\ 0 & 0 & -1 & -1 & 0 & 1\\ 0 & 0 & 1 & 1 & 0 & -1\\ 0 & -1 & 0 & 0 & -1 & 1\\ 0 & 1 & 0 & 0 & 1 & -1\end{pmatrix}^T \cdot \begin{pmatrix}p_{P+N\rightarrow PN}\\ p_{PN\rightarrow P+N}\\ p_{P+S\rightarrow PS}\\ p_{PS\rightarrow P+S}\\ p_{PN+S\rightarrow PSN}\\ p_{PSN\rightarrow PN+S}\\ p_{PS+N\rightarrow PSN}\\ p_{PSN\rightarrow PS+N}\end{pmatrix}.$$

Generally, by denoting the reaction matrix $\boldsymbol{M}$, the state vector $\boldsymbol{x}$, and the propensity vector $\boldsymbol{p}$, the nonlinear rate equations can be written as,

$$\frac{d\boldsymbol{x}}{dt} = \boldsymbol{M} \cdot \boldsymbol{p}(\boldsymbol{x}) \tag{21}$$

The reaction matrix $\boldsymbol{M}$'s dimension is $\text{Number of species} \times \text{Number of propensities}$. Since $\text{Number of propensities}$ can be very large as the reaction network gets bigger, we write $\boldsymbol{M}$ in its transpose form $\boldsymbol{M}^T$ for easy viewing. The elements in $\boldsymbol{M}$ represent the change of concentration due to reactions. E.g. The concentration of species $i$ increases $\boldsymbol{M}_{ij}$ units due to one unit of reaction $j$ happens.

#### SV.B. Stoichiometry factors.

Because of self-interactions and degeneracies, there are factors of $2$ and $1/2$ appearing in some pairwise reactions in our model. For any reaction, suppose the products have $m$ configurations and reactants have $n$ configurations. Then, its $K_{\mathrm{A}}$ should contain a factor of $m/n$, which is required for detailed balance and ensure all pairwise equilibria are met at equilibrium. One example, reaction $PN + P \leftrightharpoons PPN$ bases on the pairwise reaction $P + P \leftrightharpoons PP$ whose association constant is $K_{\mathrm{A}}^{P}$. Since $PPN$ has two configurations that either of two proteins can bind nonspecifically ($PPN$ and $PNP$), the association constant of $PN + P \leftrightharpoons PPN$ should be $2K_{\mathrm{A}}^{P}$. Another example, reaction $PPN + N \leftrightharpoons PNPN$ bases on the pairwise reaction $P + N \leftrightharpoons PN$ whose association constant is $K_{\mathrm{A}}^{N}$. Since $PPN + N \leftrightharpoons PNPN$ happens in 1D (Eq. 5) and $PPN$ has two configurations, its association constant is $\frac{\gamma K_{\mathrm{A}}^{N}}{2}$.

Stoichiometry factors also exist in propensity functions and thus rate equations. They depend on the possible reaction pathways. For reactions like $PN + P \leftrightharpoons PPN$ and

$PP + S \leftrightharpoons PPS$, there are two possible association pathways to one configuration, and the forward reactions have a stoichiometry factor 2. For reactions like $PPN + N \leftrightharpoons PNPN$, there are two $N$ sites available for dissociation, and the backward reaction has a stoichiometry factor 2. For reactions like $PPN + S \leftrightharpoons PPSN$ and $PPN + S \leftrightharpoons PNPS$, although there are multiple configurations in both reactants and products, they only have 1 reaction pathways since $PPSN$ and $PNPS$ are two species. The $S$ site must bind with the designated protein, and therefore the stoichiometry factors are 1.

#### SV.C. Environment DNA

In this environment, a protein ($P$) can bind nonspecific DNA ($N$) and can form dimers. The system contains 6 species (Fig. S2), $[P], [PP], [N], [PN], [PPN], [PNPN]$, with 6 pairwise reactions (Table S2). Using $[P]_{\text{eq}}$, $[N]_{\text{eq}}$, $K_{\text{A}}^{P}$, and $K_{\text{A}}^{N}$, the equilibrium concentrations are,

$$\begin{gathered}[PP]_{\text{eq}} = K_A^P[P]_{\text{eq}} \\ [PN]_{\text{eq}} = K_A^N[N]_{\text{eq}}[P]_{\text{eq}} \\ [PPN]_{\text{eq}} = 2K_A^N[N]_{\text{eq}}K_{\text{A}}^P[P]_{\text{eq}}^2 \\ [PNPN]_{\text{eq}} = \gamma\left(K_A^N[N]_{\text{eq}}\right)^2 K_A^P[P]_{\text{eq}}^2\end{gathered} \tag{22}$$

In this environment, we solve the reaction system,

$$\frac{d\boldsymbol{x}^{\text{DNA}}}{dt} = \boldsymbol{M}^{\text{DNA}} \cdot \boldsymbol{p}^{\text{DNA}}(\boldsymbol{x}^{\text{DNA}}) \tag{23}$$

where the state vector is,

$$\boldsymbol{x}^{\text{DNA}} = \begin{pmatrix}[P] \\ [PP] \\ [N] \\ [PN] \\ [PPN] \\ [PNPN]\end{pmatrix}. \tag{24a}$$

The propensity vector is,

$$\boldsymbol{p}^{\text{DNA}}(\boldsymbol{x}^{\text{DNA}}) = \begin{pmatrix} p_{P+P\to PP} \\ p_{PP\to P+P} \\ p_{P+PN\to PPN} \\ p_{PPN\to P+PN} \\ p_{P+N\to PN} \\ p_{PN\to P+N} \\ p_{PP+N\to PPN} \\ p_{PPN\to PP+N} \\ p_{PPN+N\to PNPN} \\ p_{PNPN\to PPN+N} \\ p_{PN+PN\to NPPN} \\ p_{PNPN\to PN+PN} \end{pmatrix}. \tag{24b}$$

The reaction matrix is,

$$(\boldsymbol{M}^{\text{DNA}})^T = \begin{pmatrix} -2 & 1 & 0 & 0 & 0 & 0 \\ 2 & -1 & 0 & 0 & 0 & 0 \\ -1 & 0 & 0 & -1 & 1 & 0 \\ 1 & 0 & 0 & 1 & -1 & 0 \\ -1 & 0 & -1 & 1 & 0 & 0 \\ 1 & 0 & 1 & -1 & 0 & 0 \\ 0 & -1 & -1 & 0 & 1 & 0 \\ 0 & 1 & 1 & 0 & -1 & 0 \\ 0 & 0 & -1 & 0 & -1 & 1 \\ 0 & 0 & 1 & 0 & 1 & -1 \\ 0 & 0 & 0 & -2 & 0 & 1 \\ 0 & 0 & 0 & 2 & 0 & -1 \end{pmatrix}. \tag{24c}$$

#### SV.D. Environment DNA + targ

In this environment, a protein ($P$) can bind nonspecific DNA ($N$) and targets ($S$) and can form dimers. The system contains 13 species (Fig. S3), $[P]$, $[PP]$, $[N]$, $[PN]$, $[PPN]$, $[PNPN]$, $[PS]$, $[PSN]$, $[PPS]$, $[PNPS]$, $[PPSN]$, $[PNPSN]$, $[S]$, with 21 pairwise reactions (Table S3). DNA + targ includes the original 6 reactions and an additional 15 reactions from monomer addition reactions (e.g. $P + PSN \rightleftharpoons PPSN$), two-two reactions (e.g. $PN + PS \rightleftharpoons PNPS$), and two-three reactions (e.g. $PSN + PN \rightleftharpoons PNPSN$) (Figure S3). Using $[P]_{\text{eq}}$, $[N]_{\text{eq}}$, $[S]_{\text{eq}}$, $K_{\text{A}}^{P}$, $K_{\text{A}}^{N}$, and $K_{\text{A}}^{S}$, the equilibrium concentrations are,

$$
\begin{gathered}
[PP]_{\text{eq}} = K_{\text{A}}^{P}[P]_{\text{eq}} \\
[PN]_{\text{eq}} = K_{\text{A}}^{N}[N]_{\text{eq}}[P]_{\text{eq}} \\
[PPN]_{\text{eq}} = 2K_{\text{A}}^{N}[N]_{\text{eq}}K_{A}^{P}[P]_{\text{eq}}^{2} \\
[PNPN]_{\text{eq}} = \gamma\left(K_{A}^{N}[N]_{\text{eq}}\right)^{2}K_{A}^{P}[P]_{\text{eq}}^{2} \\
[PS]_{\text{eq}} = K_{\text{A}}^{S}[S]_{\text{eq}}[P]_{\text{eq}} \\
[PSN]_{\text{eq}} = \gamma K_{\text{A}}^{N}[N]_{\text{eq}}K_{A}^{S}[S]_{\text{eq}}[P]_{\text{eq}} \\
[PPS]_{\text{eq}} = 2K_{A}^{S}[S]_{\text{eq}}K_{\text{A}}^{P}[P]_{\text{eq}}^{2} \\
[PNPS]_{\text{eq}} = 2\gamma K_{\text{A}}^{N}[N]_{\text{eq}}K_{\text{A}}^{S}[S]_{\text{eq}}K_{A}^{P}[P]_{\text{eq}}^{2} \\
[PPSN]_{\text{eq}} = 2\gamma K_{\text{A}}^{N}[N]_{\text{eq}}K_{\text{A}}^{S}[S]_{\text{eq}}K_{A}^{P}[P]_{\text{eq}}^{2} \\
[PNPSN]_{\text{eq}} = 2\left(\gamma K_{\text{A}}^{N}[N]_{\text{eq}}\right)^{2}K_{A}^{S}[S]_{\text{eq}}K_{\text{A}}^{P}[P]_{\text{eq}}^{2}
\end{gathered} \tag{24}
$$

In this environment, we solve the reaction system,

$$
\frac{d\boldsymbol{x}^{\text{DNA+targ}}}{dt} = \boldsymbol{M}^{\text{DNA+targ}} \cdot \boldsymbol{p}^{\text{DNA+targ}}\left(\boldsymbol{x}^{\text{DNA+targ}}\right) \tag{25}
$$

where the state vector is,

$$
\boldsymbol{x}^{\text{DNA+targ}} = \begin{pmatrix} \boldsymbol{x}^{\text{DNA}} \\ [S] \\ [PS] \\ [PSN] \\ [PPS] \\ [PNPS] \\ [PPSN] \\ [PNPSN] \end{pmatrix}. \tag{26a}
$$

The propensity vector is,

$$
\boldsymbol{p}^{\mathrm{DNA+targ}}(\boldsymbol{x}^{\mathrm{DNA+targ}}) = \begin{pmatrix} \boldsymbol{p}^{\mathrm{DNA}}(\boldsymbol{x}^{\mathrm{DNA+targ}}) \\ p_{P+S\to PS} \\ p_{PS\to P+S} \\ p_{PP+S\to PPS} \\ p_{PPS\to PP+S} \\ p_{PS+P\to PPS} \\ p_{PPS\to PS+P} \\ p_{PSN+P\to PPSN} \\ p_{PPSN\to PSN+P} \\ p_{PS+N\to PSN} \\ p_{PSN\to PS+N} \\ p_{PPS+N\to PNPS} \\ p_{PNPS\to PPS+N} \\ p_{PPS+N\to PPSN} \\ p_{PPSN\to PPS+N} \\ p_{PNPS+N\to PNPSN} \\ p_{PNPSN\to PNPS+N} \\ p_{PPSN+N\to PNPSN} \\ p_{PNPSN\to PPSN+N} \\ p_{PN+S\to PSN} \\ p_{PSN\to PN+S} \\ p_{PPN+S\to PNPS} \\ p_{PNPS\to PPN+S} \\ p_{PPN+S\to PPSN} \\ p_{PPSN\to PPN+S} \\ p_{PNPN+S\to PNPSN} \\ p_{PNPSN\to PNPN+S} \\ p_{PN+PS\to PNPS} \\ p_{PNPS\to PN+PS} \\ p_{PSN+PN\to PNPSN} \\ p_{PNPSN\to PSN+PN} \end{pmatrix}. \qquad (26b)
$$

The reaction matrix is,

$$
(\boldsymbol{M}^{\text{DNA+targ}})^T = \begin{pmatrix}
\multicolumn{6}{c}{(\boldsymbol{M}^{\text{DNA}})^T} & \multicolumn{7}{c}{\mathbf{0}} \\
-1 & 0 & 0 & 0 & 0 & 0 & -1 & 1 & 0 & 0 & 0 & 0 & 0 \\
1 & 0 & 0 & 0 & 0 & 0 & 1 & -1 & 0 & 0 & 0 & 0 & 0 \\
0 & -1 & 0 & 0 & 0 & 0 & -1 & 0 & 0 & 1 & 0 & 0 & 0 \\
0 & 1 & 0 & 0 & 0 & 0 & 1 & 0 & 0 & -1 & 0 & 0 & 0 \\
-1 & 0 & 0 & 0 & 0 & 0 & 0 & -1 & 0 & 1 & 0 & 0 & 0 \\
1 & 0 & 0 & 0 & 0 & 0 & 0 & 1 & 0 & -1 & 0 & 0 & 0 \\
-1 & 0 & 0 & 0 & 0 & 0 & 0 & 0 & -1 & 0 & 0 & 1 & 0 \\
1 & 0 & 0 & 0 & 0 & 0 & 0 & 0 & 1 & 0 & 0 & -1 & 0 \\
0 & 0 & -1 & 0 & 0 & 0 & 0 & -1 & 1 & 0 & 0 & 0 & 0 \\
0 & 0 & 1 & 0 & 0 & 0 & 0 & 1 & -1 & 0 & 0 & 0 & 0 \\
0 & 0 & -1 & 0 & 0 & 0 & 0 & 0 & 0 & -1 & 1 & 0 & 0 \\
0 & 0 & 1 & 0 & 0 & 0 & 0 & 0 & 0 & 1 & -1 & 0 & 0 \\
0 & 0 & -1 & 0 & 0 & 0 & 0 & 0 & 0 & -1 & 0 & 1 & 0 \\
0 & 0 & 1 & 0 & 0 & 0 & 0 & 0 & 0 & 1 & 0 & -1 & 0 \\
0 & 0 & -1 & 0 & 0 & 0 & 0 & 0 & 0 & 0 & -1 & 0 & 1 \\
0 & 0 & 1 & 0 & 0 & 0 & 0 & 0 & 0 & 0 & 1 & 0 & -1 \\
0 & 0 & -1 & 0 & 0 & 0 & 0 & 0 & 0 & 0 & 0 & -1 & 1 \\
0 & 0 & 1 & 0 & 0 & 0 & 0 & 0 & 0 & 0 & 0 & 1 & -1 \\
0 & 0 & 0 & -1 & 0 & 0 & -1 & 0 & -1 & 0 & 0 & 0 & 0 \\
0 & 0 & 0 & 1 & 0 & 0 & 1 & 0 & 1 & 0 & 0 & 0 & 0 \\
0 & 0 & 0 & 0 & -1 & 0 & -1 & 0 & 0 & 0 & 1 & 0 & 0 \\
0 & 0 & 0 & 0 & 1 & 0 & 1 & 0 & 0 & 0 & -1 & 0 & 0 \\
0 & 0 & 0 & 0 & -1 & 0 & -1 & 0 & 0 & 0 & 0 & 1 & 0 \\
0 & 0 & 0 & 0 & 1 & 0 & 1 & 0 & 0 & 0 & 0 & -1 & 0 \\
0 & 0 & 0 & 0 & 0 & -1 & -1 & 0 & 0 & 0 & 0 & 0 & 1 \\
0 & 0 & 0 & 0 & 0 & 1 & 1 & 0 & 0 & 0 & 0 & 0 & -1 \\
0 & 0 & 0 & -1 & 0 & 0 & 0 & -1 & 0 & 0 & 1 & 0 & 0 \\
0 & 0 & 0 & 1 & 0 & 0 & 0 & 1 & 0 & 0 & -1 & 0 & 0 \\
0 & 0 & 0 & -1 & 0 & 0 & 0 & 0 & -1 & 0 & 0 & 0 & 1 \\
0 & 0 & 0 & 1 & 0 & 0 & 0 & 0 & 1 & 0 & 0 & 0 & -1
\end{pmatrix}. \qquad (26c)
$$

#### SV.E. Environment DNA + clusTarg

To account for the clustering of the S sites, instead of tracking the concentration of $S$, we will track the concentration of $S_2$. The initial concentrations are therefore set to $[S_2] = [S]/2$ so that the number of sites is equivalent. This is important because $S$ is not a diffusible site, so we cannot consume it within bimolecular reactions where the other reactant also does not diffuse. There are 16 species in this model (Fig. S4): $[P]$, $[N]$, $[PP]$, $[PN]$, $[PPN]$, $[PNPN]$, $[S_2]$, $[PS_2]$, $[PS_2N]$, $[PPS_2]$, $[PPS_2N]$, $[PNPS_2]$, $[PNPS_2N]$, $[PSPS]$, $[PSPSN]$, $[PSNPSN]$. In our model, we forbid two proteins to be bound to both specific sites without being bound to each other. This eliminates recruitment of $P$ to $PS_2$ via DNA binding, it must arrive via a protein-protein interaction from 3D or 1D. 27 Pairwise reactions are listed in Table S4. Using $[P]_{\mathrm{eq}}$, $[N]_{\mathrm{eq}}$, $[S_2]_{\mathrm{eq}}$, $K_{\mathrm{A}}^{P}$, $K_{\mathrm{A}}^{N}$, and $K_{\mathrm{A}}^{S}$, the equilibrium concentrations are,

$$
\begin{gathered}
[PP]_{\mathrm{eq}} = K_{\mathrm{A}}^{P}[P]_{\mathrm{eq}} \\
[PN]_{\mathrm{eq}} = K_{A}^{N}[N]_{\mathrm{eq}}[P]_{\mathrm{eq}} \\
[PPN]_{\mathrm{eq}} = 2K_{\mathrm{A}}^{N}[N]_{\mathrm{eq}}K_{\mathrm{A}}^{P}[P]_{\mathrm{eq}}^{2} \\
[PNPN]_{\mathrm{eq}} = \gamma\left(K_{\mathrm{A}}^{N}[N]_{\mathrm{eq}}\right)^{2}K_{\mathrm{A}}^{P}[P]_{\mathrm{eq}}^{2} \\
[PS_2]_{\mathrm{eq}} = 2K_{\mathrm{A}}^{S}[S_2]_{\mathrm{eq}}[P]_{\mathrm{eq}} \\
[PS_2N]_{\mathrm{eq}} = 2\gamma K_{\mathrm{A}}^{N}[N]_{\mathrm{eq}}K_{\mathrm{A}}^{S}[S_2]_{\mathrm{eq}}[P]_{\mathrm{eq}} \\
[PPS_2]_{\mathrm{eq}} = 4K_{\mathrm{A}}^{S}[S_2]_{\mathrm{eq}}K_{\mathrm{A}}^{P}[P]_{\mathrm{eq}}^{2} \\
[PNPS_2]_{\mathrm{eq}} = 4\gamma K_{\mathrm{A}}^{N}[N]_{\mathrm{eq}}K_{\mathrm{A}}^{S}[S_2]_{\mathrm{eq}}K_{\mathrm{A}}^{P}[P]_{\mathrm{eq}}^{2} \\
[PPS_2N]_{\mathrm{eq}} = 4\gamma K_{\mathrm{A}}^{N}[N]_{\mathrm{eq}}K_{\mathrm{A}}^{S}[S_2]_{\mathrm{eq}}K_{\mathrm{A}}^{P}[P]_{\mathrm{eq}}^{2} \\
[PNPS_2N]_{\mathrm{eq}} = 4\left(\gamma K_{\mathrm{A}}^{N}[N]_{\mathrm{eq}}\right)^{2}K_{\mathrm{A}}^{S}[S_2]_{\mathrm{eq}}K_{\mathrm{A}}^{P}[P]_{\mathrm{eq}}^{2} \\
[PSPS]_{\mathrm{eq}} = 2K_{\mathrm{A}}^{S}C_0K_{\mathrm{A}}^{S}[S_2]_{\mathrm{eq}}K_{\mathrm{A}}^{P}[P]_{\mathrm{eq}}^{2} \\
[PSPSN]_{\mathrm{eq}} = 4K_{\mathrm{A}}^{S}C_0\gamma K_{\mathrm{A}}^{N}[N]_{\mathrm{eq}}K_{\mathrm{A}}^{S}[S_2]_{\mathrm{eq}}K_{\mathrm{A}}^{P}[P]_{\mathrm{eq}}^{2} \\
[PSNPSN]_{\mathrm{eq}} = 2K_{\mathrm{A}}^{S}C_0\left(\gamma K_{\mathrm{A}}^{N}[N]_{\mathrm{eq}}\right)^{2}K_{\mathrm{A}}^{S}[S_2]_{\mathrm{eq}}K_{\mathrm{A}}^{P}[P]_{\mathrm{eq}}^{2}
\end{gathered}
\tag{26}
$$

In this environment, we solve the reaction system,

$$
\frac{d\boldsymbol{x}^{\mathrm{DNA+clusTarg}}}{dt} = \boldsymbol{M}^{\mathrm{DNA+clusTarg}} \cdot \boldsymbol{p}^{\mathrm{DNA+clusTarg}}\left(\boldsymbol{x}^{\mathrm{DNA+clusTarg}}\right) \tag{27}
$$

where the state vector is,

$$
\boldsymbol{x}^{\mathrm{DNA+clusTarg}} = \begin{pmatrix} \boldsymbol{x}^{\mathrm{DNA}} \\ [S_2] \\ [PS_2] \\ [PS_2N] \\ [PPS_2] \\ [PNPS_2] \\ [PPS_2N] \\ [PNPS_2N] \\ [PSPS], \\ [PSPSN], \\ [PSNPSN] \end{pmatrix}. \tag{28a}
$$

The first 12 elements of $\boldsymbol{p}^{\mathrm{DNA+clusTarg}}$ are the same elements as $\boldsymbol{p}^{\mathrm{DNA}}$, and the others read,

$$
\boldsymbol{p}^{\mathrm{DNA+targ}}\left(\boldsymbol{x}^{\mathrm{DNA+clusTarg}}\right) = \begin{pmatrix}
\boldsymbol{p}^{\mathrm{DNA}}\left(\boldsymbol{x}^{\mathrm{DNA+clusTarg}}\right) \\
p_{P+S_2\to PS_2} \\
p_{PS_2\to P+S_2} \\
p_{PP+S_2\to PPS_2} \\
p_{PPS_2\to PP+S_2} \\
p_{PS_2+P\to PPS_2} \\
p_{PPS_2\to PS_2+P} \\
p_{PS_2N+P\to PPS_2N} \\
p_{PPS_2N\to PS_2N+P} \\
p_{PS_2+N\to PS_2N} \\
p_{PS_2N\to PS_2+N} \\
p_{PPS_2+N\to PNPS_2} \\
p_{PNPS_2\to PPS_2+N} \\
p_{PPS_2+N\to PPS_2N} \\
p_{PPS_2N\to PPS_2+N} \\
p_{PNPS_2+N\to PNPS_2N} \\
p_{PNPS_2N\to PNPS_2+N} \\
p_{PPS_2N+N\to PNPS_2N} \\
p_{PNPS_2N\to PPS_2N+N} \\
p_{PSPS+N\to PSPSN} \\
p_{PSPSN\to PSPS+N} \\
p_{PSPSN+N\to PSNPSN} \\
p_{PSNPSN\to PSPSN+N} \\
p_{PN+S_2\to PS_2N} \\
p_{PS_2N\to PN+S_2} \\
p_{PPN+S_2\to PNPS_2} \\
p_{PNPS_2\to PPN+S_2} \\
p_{PPN+S_2\to PPS_2N} \\
p_{PPS_2N\to PPN+S_2} \\
p_{PNPN+S_2\to PNPS_2N} \\
p_{PNPS_2N\to PNPN+S_2} \\
p_{PN+PS_2\to PNPS_2} \\
p_{PNPS_2\to PN+PS_2} \\
p_{PS_2N+PN\to PNPS_2N} \\
p_{PNPS_2N\to PS_2N+PN} \\
p_{PPS_2\to PSPS} \\
p_{PSPS\to PPS_2} \\
p_{PPS_2N\to PSPSN} \\
p_{PSPSN\to PPS_2N} \\
p_{PNPS_2\to PSPSN} \\
p_{PSPSN\to PNPS_2} \\
p_{PNPS_2N\to PSNPSN} \\
p_{PSNPSN\to PNPS_2N}
\end{pmatrix}. \qquad (28b)
$$

The first 12 rows of $\boldsymbol{M}^{\mathrm{DNA+clusTarg}}$ are $(\boldsymbol{M}^{\mathrm{DNA}} \quad \boldsymbol{0})$. The full reaction matrix is,

$$
\left(\boldsymbol{M}^{\text{DNA+clusTarg}}\right)^{T} =
\left(\begin{array}{cccccccccccccccc}
\multicolumn{6}{c}{\left(\boldsymbol{M}^{\text{DNA}}\right)^{T}} & \multicolumn{10}{c}{\mathbf{0}} \\
-1 & 0 & 0 & 0 & 0 & 0 & -2 & 1 & 0 & 0 & 0 & 0 & 0 & 0 & 0 & 0 \\
1 & 0 & 0 & 0 & 0 & 0 & 2 & -1 & 0 & 0 & 0 & 0 & 0 & 0 & 0 & 0 \\
0 & -1 & 0 & 0 & 0 & 0 & -2 & 0 & 0 & 1 & 0 & 0 & 0 & 0 & 0 & 0 \\
0 & 1 & 0 & 0 & 0 & 0 & 2 & 0 & 0 & -1 & 0 & 0 & 0 & 0 & 0 & 0 \\
-1 & 0 & 0 & 0 & 0 & 0 & 0 & -1 & 0 & 1 & 0 & 0 & 0 & 0 & 0 & 0 \\
1 & 0 & 0 & 0 & 0 & 0 & 0 & 1 & 0 & -1 & 0 & 0 & 0 & 0 & 0 & 0 \\
-1 & 0 & 0 & 0 & 0 & 0 & 0 & 0 & -1 & 0 & 0 & 1 & 0 & 0 & 0 & 0 \\
1 & 0 & 0 & 0 & 0 & 0 & 0 & 0 & 1 & 0 & 0 & -1 & 0 & 0 & 0 & 0 \\
0 & 0 & -1 & 0 & 0 & 0 & 0 & -1 & 1 & 0 & 0 & 0 & 0 & 0 & 0 & 0 \\
0 & 0 & 1 & 0 & 0 & 0 & 0 & 1 & -1 & 0 & 0 & 0 & 0 & 0 & 0 & 0 \\
0 & 0 & -1 & 0 & 0 & 0 & 0 & 0 & 0 & -1 & 1 & 0 & 0 & 0 & 0 & 0 \\
0 & 0 & 1 & 0 & 0 & 0 & 0 & 0 & 0 & 1 & -1 & 0 & 0 & 0 & 0 & 0 \\
0 & 0 & -1 & 0 & 0 & 0 & 0 & 0 & 0 & -1 & 0 & 1 & 0 & 0 & 0 & 0 \\
0 & 0 & 1 & 0 & 0 & 0 & 0 & 0 & 0 & 1 & 0 & -1 & 0 & 0 & 0 & 0 \\
0 & 0 & -1 & 0 & 0 & 0 & 0 & 0 & 0 & 0 & -1 & 0 & 1 & 0 & 0 & 0 \\
0 & 0 & 1 & 0 & 0 & 0 & 0 & 0 & 0 & 0 & 1 & 0 & -1 & 0 & 0 & 0 \\
0 & 0 & -1 & 0 & 0 & 0 & 0 & 0 & 0 & 0 & 0 & -1 & 1 & 0 & 0 & 0 \\
0 & 0 & 1 & 0 & 0 & 0 & 0 & 0 & 0 & 0 & 0 & 1 & -1 & 0 & 0 & 0 \\
0 & 0 & -1 & 0 & 0 & 0 & 0 & 0 & 0 & 0 & 0 & 0 & 0 & -1 & 1 & 0 \\
0 & 0 & 1 & 0 & 0 & 0 & 0 & 0 & 0 & 0 & 0 & 0 & 0 & 1 & -1 & 0 \\
0 & 0 & -1 & 0 & 0 & 0 & 0 & 0 & 0 & 0 & 0 & 0 & 0 & 0 & -1 & 1 \\
0 & 0 & 1 & 0 & 0 & 0 & 0 & 0 & 0 & 0 & 0 & 0 & 0 & 0 & 1 & -1 \\
0 & 0 & 0 & -1 & 0 & 0 & -2 & 0 & -1 & 0 & 0 & 0 & 0 & 0 & 0 & 0 \\
0 & 0 & 0 & 1 & 0 & 0 & 2 & 0 & 1 & 0 & 0 & 0 & 0 & 0 & 0 & 0 \\
0 & 0 & 0 & 0 & -1 & 0 & -2 & 0 & 0 & 0 & 1 & 0 & 0 & 0 & 0 & 0 \\
0 & 0 & 0 & 0 & 1 & 0 & 2 & 0 & 0 & 0 & -1 & 0 & 0 & 0 & 0 & 0 \\
0 & 0 & 0 & 0 & -1 & 0 & -2 & 0 & 0 & 0 & 0 & 1 & 0 & 0 & 0 & 0 \\
0 & 0 & 0 & 0 & 1 & 0 & 2 & 0 & 0 & 0 & 0 & -1 & 0 & 0 & 0 & 0 \\
0 & 0 & 0 & 0 & 0 & -1 & -2 & 0 & 0 & 0 & 0 & 0 & 1 & 0 & 0 & 0 \\
0 & 0 & 0 & 0 & 0 & 1 & 2 & 0 & 0 & 0 & 0 & 0 & -1 & 0 & 0 & 0 \\
0 & 0 & 0 & -1 & 0 & 0 & 0 & -1 & 0 & 0 & 1 & 0 & 0 & 0 & 0 & 0 \\
0 & 0 & 0 & 1 & 0 & 0 & 0 & 1 & 0 & 0 & -1 & 0 & 0 & 0 & 0 & 0 \\
0 & 0 & 0 & -1 & 0 & 0 & 0 & 0 & -1 & 0 & 0 & 0 & 1 & 0 & 0 & 0 \\
0 & 0 & 0 & 1 & 0 & 0 & 0 & 0 & 1 & 0 & 0 & 0 & -1 & 0 & 0 & 0 \\
0 & 0 & 0 & 0 & 0 & 0 & 0 & 0 & 0 & -1 & 0 & 0 & 0 & 1 & 0 & 0 \\
0 & 0 & 0 & 0 & 0 & 0 & 0 & 0 & 0 & 1 & 0 & 0 & 0 & -1 & 0 & 0 \\
0 & 0 & 0 & 0 & 0 & 0 & 0 & 0 & 0 & 0 & 0 & -1 & 0 & 0 & 1 & 0 \\
0 & 0 & 0 & 0 & 0 & 0 & 0 & 0 & 0 & 0 & 0 & 1 & 0 & 0 & -1 & 0 \\
0 & 0 & 0 & 0 & 0 & 0 & 0 & 0 & 0 & 0 & -1 & 0 & 0 & 0 & 1 & 0 \\
0 & 0 & 0 & 0 & 0 & 0 & 0 & 0 & 0 & 0 & 1 & 0 & 0 & 0 & -1 & 0 \\
0 & 0 & 0 & 0 & 0 & 0 & 0 & 0 & 0 & 0 & 0 & 0 & -1 & 0 & 0 & 1 \\
0 & 0 & 0 & 0 & 0 & 0 & 0 & 0 & 0 & 0 & 0 & 0 & 1 & 0 & 0 & -1
\end{array}\right). \qquad (28c)
$$

The blue elements in $\boldsymbol{M}^{\mathrm{DNA+clusTarg}}$ correspond to the new 10 elements in $\boldsymbol{x}^{\mathrm{DNA+clusTarg}}$ not appearing in $\boldsymbol{x}^{\mathrm{DNA}}$.

#### SVI. Approximations to derive analytical expressions for dwell times, recruitment fraction, and target occupancy in all environments

We first summarize the approach to simplify the exact expressions. The exact expressions can all be written as functions of known parameters and the equilibrium concentrations of free proteins and free DNA binding sites, or $[P]_{\mathrm{eq}}$, $[N]_{\mathrm{eq}}$, and $[S]_{\mathrm{eq}}$. In general, at equilibrium we end up with coupled quartic or quintic equations (see section SVI.A), such that we cannot solve $[P]_{\mathrm{eq}}$ exactly. To a good approximation, we can assert that $[N]_{\mathrm{eq}} \approx [N]_{\mathrm{tot}}$, as the nonspecific sites outnumber the protein copies. The primary unknowns are therefore $[P]_{\mathrm{eq}}$ and $[S]_{\mathrm{eq}}$. To recover useful analytical formulas, we will derive expressions at two limits: monomer ($K_{\mathrm{eq}}^{P} \rightarrow 0$) and irreversible dimer ($K_{\mathrm{A}}^{P} \rightarrow \infty$). We also derive approximations to $[P]_{\mathrm{eq}}$ that work when DNA target sites are also in excess, e.g. $[S]_{\mathrm{eq}} \sim [S]_{\mathrm{tot}}$. Our metrics also simplify under relevant regimes of strong or weak binding, which we use to compare if dimerization enhances or impairs our metrics. Key parameters are then $\chi_N = K_{\mathrm{A}}^{N}[N]_{\mathrm{tot}}$ , which is the constant partition coefficient for a protein from solution (3D) to nonspecific DNA, or the propensity for nonspecific binding. We similarly define the constant $\chi_S = K_{\mathrm{A}}^{S}[S]_{\mathrm{tot}}$, the 3D partition coefficient from solution to target DNA. The 1D partition coefficients follow from multiplication by $\gamma$.

##### SVI.A. General estimation of the equilibrium concentration of protein monomers in solution

Assuming $[S]_{\mathrm{eq}} = [S]_{\mathrm{tot}}$ and $[N]_{\mathrm{eq}} = [N]_{\mathrm{tot}}$, and denoting $\chi_S = K_{\mathrm{A}}^{S}[S]_{\mathrm{tot}}$ and $\chi_N = K_{\mathrm{A}}^{N}[N]_{\mathrm{tot}}$, we now only need to solve $[P]_{\mathrm{eq}}$ to obtain the equilibrium. By mass conservation, we have,

$$[P] + 2[PP] + \sum_{j=1}^{Nb} a_{AB_j}\left[AB_j\right] = [P]_{\mathrm{tot}} \tag{28}$$

where $\sum_{j=1}^{Nb} a_{AB_j}\left[AB_j\right]$ sums over all proteins bound on DNA, defined by Eqs. 13, 15, and 17. At equilibrium, the concentrations on the left hand can be replaced by the equilibrium equations defined in Eqs. 23, 25, and 27, respectively. Now, Eq. 29 is quadratic for $[P]_{\mathrm{eq}}$ and can be rewritten in the form of,

$$A \cdot \frac{[P]_{\mathrm{eq}}}{[P]_{\mathrm{tot}}} + 2 \cdot B \cdot K_A^P [P]_{\mathrm{tot}} \cdot \left(\frac{[P]_{\mathrm{eq}}}{[P]_{\mathrm{tot}}}\right)^2 - 1 = 0. \tag{29}$$

Its positive root is physiologically meaningful and thus,

$$[P]_{\text{eq}} = [P]_{\text{tot}} \cdot \frac{-B + \sqrt{B^2(K_A^P[P]_{\text{tot}})^2 + A}}{A} \tag{30}$$

For environment DNA, we have (from Eqs. 13 and 23),

$$\begin{cases} A^{\text{DNA}} = 1 + \chi_N \\ B^{\text{DNA}} = 1 + 2\chi_N + \gamma\chi_N^2 \end{cases}. \tag{31$a$}$$

For environment DNA + targ, we have (from Eqs. 15 and 25),

$$\begin{cases} A^{\text{DNA+targ}} = 1 + \chi_N + \chi_S + \gamma\chi_N\chi_S \\ B^{\text{DNA+targ}} = 1 + 2\chi_N + \gamma\chi_N^2 + 2\chi_S(1 + \gamma\chi_N)^2 \end{cases}. \tag{31$b$}$$

For environment DNA + clusTarg, we have (from Eqs. 17 and 27),

$$\begin{cases} A^{\text{DNA+clusTarg}} = 1 + \chi_N + \chi_S + \gamma\chi_N\chi_S \\ B^{\text{DNA+clusTarg}} = 1 + 2\chi_N + \gamma\chi_N^2 + \left(2\chi_S + K_{\text{A}}^S C_0\chi_S\right)(1 + \gamma\chi_N)^2 \end{cases}. \tag{31c}$$

Note that it is even not practical to solve $[P]_{\text{eq}}$ in the simplest DNA environment without assuming $[N]_{\text{eq}} = [N]_{\text{tot}}$, which requires to find the root of the top two coupled nonlinear equations:

$$\begin{cases} \left(1 + K_{\text{A}}^N[P]_{\text{eq}} + 2K_{\text{A}}^N K_{\text{A}}^P[P]_{\text{eq}}^2\right)[N]_{\text{eq}} + 2K_{\text{A}}^P[P]_{\text{eq}}^2\gamma\left(K_{\text{A}}^N[N]_{\text{eq}}\right)^2 = [N]_{\text{tot}} \\ [P]_{\text{eq}} = [P]_{\text{tot}} \cdot \dfrac{-B + \sqrt{B^2(K_{\text{A}}^P[P]_{\text{tot}})^2 + A}}{A} \\ A = 1 + K_{\text{A}}^N[N]_{\text{eq}} \\ B = 1 + 2K_{\text{A}}^N[N]_{\text{eq}} + \gamma\left(K_{\text{A}}^N[N]_{\text{eq}}\right)^2 \end{cases}. \tag{31d}$$

Therefore, our analytical approximations either rely on the assumptions of $[S]_{\text{eq}} = [S]_{\text{tot}}$ and $[N]_{\text{eq}} = [N]_{\text{tot}}$, or require considering all proteins are monomers or dimers, or both.

#### SVI.B Nonspecific binding only (environment DNA).

To derive closed-form analytical expressions for the dwell time and the recruitment fraction, we start by plugging the equilibrium equations (Eq. 23) into Eq. 12, 13, and 14, to get,

$$\tau^{\text{DNA}} = \frac{1}{k_{\text{off}}^N} \cdot \frac{\dfrac{1}{K_A^P[P]_{\text{eq}}} + 4 + 2\gamma K_A^N[N]_{\text{eq}}}{\dfrac{1}{K_A^P[P]_{\text{eq}}} + 4 + 2\dfrac{k_{\text{off}}^P}{k_{\text{off}}^N}}. \tag{31}$$

which can be evaluated numerically (see section SXII.A). In the same way, the recruitment fraction reads from equilibrium equations (Eq. 23), and Eq. 19 and 13.

$$\theta^{\mathrm{DNA}} = \frac{K_{\mathrm{A}}^{N}[N]_{\mathrm{eq}} + 2K_{A}^{P}[P]_{\mathrm{eq}}\left(2K_{A}^{N}[N]_{\mathrm{eq}} + \gamma\left(K_{A}^{N}[N]_{\mathrm{eq}}\right)^{2}\right)}{K_{A}^{N}[N]_{\mathrm{eq}} + 1 + 2K_{\mathrm{A}}^{P}[P]_{\mathrm{eq}}\left(2K_{\mathrm{A}}^{N}[N]_{\mathrm{eq}} + \gamma\left(K_{A}^{N}[N]_{\mathrm{eq}}\right)^{2} + 1\right)} \tag{32}$$

which can be evaluated numerically (see section SXII.A). Eq. 32 and 33 are exact. However, analytically evaluating them requires knowledge of $[P]_{\mathrm{eq}}$ that we can only estimate with Eq. 31 and Using $[N]_{\mathrm{eq}} = [N]_{\mathrm{tot}}$, such that Eq. 32 and 33 can be written as,

$$\tau^{\mathrm{DNA}} = \frac{1}{k_{\mathrm{off}}^{N}} \cdot \frac{\frac{1}{K_{A}^{P}[P]_{\mathrm{eq}}} + 4 + 2\gamma\chi_{N}}{\frac{1}{K_{\mathrm{A}}^{P}[P]_{\mathrm{eq}}} + 4 + 2\frac{k_{\mathrm{off}}^{P}}{k_{\mathrm{off}}^{N}}}, \tag{32'}$$

and,

$$\theta^{\mathrm{DNA}} = \frac{\chi_{N} + 2K_{\mathrm{A}}^{P}[P]_{\mathrm{eq}}(2\chi_{N} + \gamma{\chi_{N}}^{2})}{\chi_{N} + 1 + 2K_{\mathrm{A}}^{P}[P]_{\mathrm{eq}}(2\chi_{N} + \gamma{\chi_{N}}^{2} + 1)}, \tag{33'}$$

respectively.

*Limit of no protein-protein interactions (monomers):*

In the limit that there is no protein dimerization ($K_{\mathrm{A}}^{P} \to 0$), the dwell time is equivalent to the monomer dwell time, $\tau_{\mathrm{monomer}}^{\mathrm{DNA}} = 1/k_{\mathrm{off}}^{N}$ , as expected.

The recruitment fraction is also relatively simply defined as:

$$\theta_{\mathrm{monomer}}^{\mathrm{DNA}} = \frac{K_{A}^{N}[N]_{\mathrm{eq}}}{1 + K_{A}^{N}[N]_{\mathrm{eq}}} \approx \frac{\chi_{N}}{1 + \chi_{N}}, \tag{33}$$

so we see 50% recruitment when $\chi_{N} = 1$.

*Limit of irreversible protein dimers:*

At the irreversible protein-binding, $K_{\mathrm{A}}^{P} \to \infty$, or $k_{\mathrm{off}}^{P} \to 0$, the mean dwell time from Eq. 32 is,

$$\tau_{\mathrm{dimer,irr}}^{\mathrm{DNA}} = \frac{1}{k_{\mathrm{off}}^{N}} \cdot \left(1 + \frac{\gamma K_{A}^{N}[N]_{\mathrm{eq}}}{2}\right) \approx \frac{1}{k_{\mathrm{off}}^{N}} \cdot \left(1 + \frac{\gamma\chi_{N}}{2}\right). \tag{34}$$

Relative to monomer residence time, Eq. 35 derives,

$$\frac{\tau_{\mathrm{dimer,irr}}^{\mathrm{DNA}}}{\tau_{\mathrm{monomer}}^{\mathrm{DNA}}} \approx \left(1 + \frac{\gamma\chi_{N}}{2}\right). \tag{35b}$$

The recruitment fraction for dimers from Eq. 33 is:

$$\theta_{\text{dimer,irr}}^{\text{DNA}} = \frac{2K_{\text{A}}^{N}[N]_{\text{eq}} + \gamma\left(K_{\text{A}}^{N}[N]_{\text{eq}}\right)^2}{1 + 2K_{\text{A}}^{N}[N]_{\text{eq}} + \gamma\left(K_{\text{A}}^{N}[N]_{\text{eq}}\right)^2} \approx \frac{2\chi_N + \gamma{\chi_N}^2}{1 + 2\chi_N + \gamma{\chi_N}^2}. \tag{35}$$

As discussed in the main text, $\gamma\chi_N$ is the 1D partition coefficient of a protein to bind DNA, when held to a local 1D search by a partner.

#### SVI.C Approximate expressions for the dwell time, recruitment fraction, and target occupancy in environments with separated targets (DNA+targ)

Under the approximation of $[N]_{\text{eq}} = [N]_{\text{tot}}$ and $[S]_{\text{eq}} = [S]_{\text{tot}}$, we can approximate the dwell time, recruitment fraction, and target occupancy.

From Eq. 12, 15, 16, and 25, the dwell time reads,

$$\tau^{\text{DNA+targ}} = \frac{[P]_{\text{eq}}(\chi_N + \chi_S + \gamma\chi_S\chi_N) + 2K_A^P[P]_{\text{eq}}^2(2\chi_N + \gamma\chi_N^2 + 2\chi_S(1+\gamma\chi_N)^2)}{\left(4K_{\text{A}}^P[P]_{\text{eq}}^2 + [P]_{\text{eq}}\right)\left(k_{\text{off}}^N\chi_N + k_{\text{off}}^S\chi_S\right) + 2k_{\text{on}}^P[P]_{\text{eq}}^2(\chi_N + \chi_S + \gamma\chi_S\chi_N)} \tag{36}$$

From Eq. 19, 15, and 25, the recruitment fraction reads,

$$\theta^{\text{DNA+targ}} = \frac{[P]_{\text{eq}}(\chi_N + \chi_S + \gamma\chi_S\chi_N) + 2K_A^P[P]_{\text{eq}}^2(2\chi_N + \gamma\chi_N^2 + 2\chi_S(1+\gamma\chi_N)^2)}{[P]_{\text{tot}}} \tag{37}$$

From Eq. 20 and 25, the target occupancy reads,

$$\phi^{\text{DNA+targ}} = \frac{[P]_{\text{eq}}\chi_S(1+\gamma\chi_N) + 2K_{\text{A}}^P[P]_{\text{eq}}^2\chi_S(1+\gamma\chi_N)^2}{[S]_{\text{tot}}} \tag{38}$$

See section SXII.A for numerically solving these equations without approximations.

*Limit of no protein-protein interaction:*

In the limit of no protein binding, $K_{\text{A}}^{PP} \to 0$, or $k_{\text{on}}^P \to 0$, we have $\tau_{\text{monomer}}^{\text{DNA+targ}} = \tau_{\text{monomer}}^{\text{DNA+clusTarg}}$, as binding of monomers to targets is independent of the spatial separation from other targets in these models with no explicit diffusion. In other words, when proteins return to solution (3D), they are equally like to bind any of the target sites in our local environments. From Eq. 37, we have:

$$\tau_{\text{monomer}}^{\text{DNA+targ}} = \frac{K_{\text{A}}^S[S]_{\text{eq}} + K_{\text{A}}^N[N]_{\text{eq}} + \gamma K_A^N[N]_{\text{eq}}K_{\text{A}}^S[S]_{\text{eq}}}{k_{\text{on}}^S[S]_{\text{eq}} + k_{\text{on}}^N[N]_{\text{eq}}} \approx \tau_S\frac{\chi_S + \chi_N(1+\gamma\chi_S)}{\chi_S + \frac{k_{\text{off}}^N}{k_{\text{off}}^S}\chi_N}. \tag{39}$$

Where $\chi_S = K_{\text{A}}^S[S]_{\text{tot}}$ and $\tau_S = \frac{1}{k_{\text{off}}^S}$. The approximation on the right requires excess of both specific and nonspecific sites relative to proteins. Alternatively, we can approximate the exact expression in the regime where the protein binds to the targets more frequently than the nonspecific sites ($k_{\text{on}}^S[S]_{\text{eq}} \gg k_{\text{on}}^N[N]_{\text{eq}}$).

$$\tau_{\text{monomer}}^{\text{DNA+targ}} \approx \tau_S\left(1+\gamma K_{\text{A}}^{N}[N]_{\text{eq}}\right) \approx \tau_S(1+\gamma\chi_N) \tag{40}$$

The expression shows that dwell time for target bound proteins increases with $\gamma K_{\text{A}}^{N}[N]$, the 1D partition coefficient.

The recruitment fractions for these two conditions are also the same, and under the approximation of $[N]_{\text{eq}} = [N]_{\text{tot}}$ and $[S]_{\text{eq}} = [S]_{\text{tot}}$, we derive from Eq. 38:

$$\theta_{\text{monomer}}^{\text{DNA+targ}} = \theta_{\text{monomer}}^{\text{DNA+cluTarg}} \approx \frac{\chi_N + \chi_S(1+\gamma\chi_N)}{1+\chi_N+\chi_S(1+\gamma\chi_N)}. \tag{41}$$

The recruitment fraction always increases in the presence of nonspecific binding ($\frac{\partial \theta_{\text{monomer}}^{\text{DNA+targ}}}{\partial K_{\text{A}}^{N}} \geq 0$) as the addition of any new protein-DNA interactions without other changes can only reduce the number of proteins free in solution.

Similarly, they have the same target occupancy given by Eq. 39, $\phi_{\text{monomer}}^{\text{DNA+targ}} = \phi_{\text{monomer}}^{\text{DNA+clusTarg}} = \frac{K_{\text{A}}^{S}[S]_{\text{eq}}[P]_{\text{eq}}\left(1+\gamma K_{\text{A}}^{N}[N]_{\text{eq}}\right)}{[S]_{\text{tot}}}$. Since at the monomer limit $[P]_{\text{eq}} = [P]_{\text{tot}}(1-\theta)$, using Eq. 42, the above equation can be rewritten under the approximation of $[N]_{\text{eq}} = [N]_{\text{tot}}$ and $[S]_{\text{eq}} = [S]_{\text{tot}}$ as,

$$\phi_{\text{monomer}}^{\text{DNA+targ}} = \phi_{\text{monomer}}^{\text{DNA+clusTarg}} = \frac{(1+\gamma\chi_N)\chi_S}{1+\chi_N+\chi_S(1+\gamma\chi_N)} \cdot \frac{[P]_{\text{tot}}}{[S]_{\text{tot}}}. \tag{42}$$

##### *Limit of irreversible protein-protein interactions*

If proteins form irreversible dimers, for the mean dwell times of environment DNA + targ, assuming $K_{\text{A}}^{N}[N]_{\text{eq}} = \chi_N$ and $K_{\text{A}}^{S}[S]_{\text{eq}} = \chi_S$, Eq. 37 can be simplified as,

$$\tau_{\text{dimer.irr}}^{\text{DNA+targ}} \approx \frac{1}{k_{\text{off}}^{S}} \cdot \frac{2\chi_N + \gamma\chi_N^2 + 2\chi_S(1+\gamma\chi_N)^2}{2\left(k_{\text{on}}^{N}[N]_0 + k_{\text{on}}^{S}[S]_0\right)}. \tag{43}$$

To investigate how dimerization kinetics affects dwell time, we consider another case where the dimer association cannot be fully neglected. When dimerization is very strong but still reversible, where $1 \ll K_{\text{A}}^{PP}[P]_{\text{tot}} < \infty$, the mean dwell time for DNA + targ is governed by 1) DNA-bound dimers, 2) the binding of in-solution monomers ($P$) to DNA-bound monomers ($PN$, $PS$, and $PSN$), and 3) the binding of in-solution dimers ($PP$) to DNA,

$$\tau_{\text{rev-strong}}^{\text{DNA+targ}} = \frac{2\left([PPN]_{\text{eq}} + [PPS]_{\text{eq}} + [PPSN]_{\text{eq}} + [PSPN]_{\text{eq}} + [PNPN]_{\text{eq}} + [PNPSN]_{\text{eq}}\right)}{2\cdot 2[PP]_{\text{eq}}\left(k_{\text{on}}^{N}[N]_{\text{eq}} + k_{on}^{S}[S]_{\text{eq}}\right) + 2[P]_{\text{eq}}k_{\text{on}}^{P}\left([PN]_{\text{eq}} + [PS]_{\text{eq}} + [PSN]_{\text{eq}}\right)}.$$

Assuming $K_{\text{A}}^{N}[N]_{\text{eq}} = \chi_N$ and $K_{\text{A}}^{S}[S]_{\text{eq}} = \chi_S$, using Eq. 25, it can be rewritten as,

$$\tau_{\text{rev-strong}}^{\text{DNA+targ}} \approx \frac{2\chi_N + 2\chi_S + 4\gamma\chi_S\chi_N + \gamma\chi_N^2 + 2\gamma^2\chi_N^2\chi_S}{2\left(k_{\text{off}}^N\chi_N + k_{\text{off}}^S\chi_S\right) + k_{\text{off}}^P(\chi_N + \chi_S + \gamma\chi_N\chi_S)} \tag{44}$$

Returning to the irreversible dimer limit, under the approximation of $[N]_{\text{eq}} = [N]_{\text{tot}}$ and $[S]_{\text{eq}} = [S]_{\text{tot}}$, we ca also estimate recruitment fractions for environment DNA + targ according to Eq. 38,

$$\theta_{\text{dimer,irr}}^{\text{DNA+targ}} \approx \frac{2\chi_N + \gamma{\chi_N}^2 + 2\chi_S(1+\gamma\chi_N)^2}{1 + 2\chi_N + \gamma{\chi_N}^2 + 2\chi_S(1+\gamma\chi_N)^2}, \tag{45}$$

The target occupancy for environment DNA + targ is given by Eq. 39, $\phi_{\text{dimer,irr}}^{\text{DNA+targ}} = \frac{2\chi_S(1+\gamma\chi_N)^2[PP]_{\text{eq}}}{[S]_{\text{tot}}}$. Like the monomer limit, given by $[PP]_{\text{eq}} = [PP]_{\text{tot}}(1-\theta)$ and Eq. 46, we have,

$$\phi_{\text{dimer,irr}}^{\text{DNA+targ}} = \frac{2\chi_S(1+\gamma\chi_N)^2}{1 + 2\chi_N + \gamma{\chi_N}^2 + 2\chi_S(1+\gamma\chi_N)^2} \cdot \frac{[PP]_{\text{tot}}}{[S]_{\text{tot}}}, \tag{46}$$

#### SVI.D Approximate expressions for the dwell time, recruitment fraction, and target occupancy in environments with clustered targets (DNA+clusTarg)

In this model, we do not need to repeat the metrics in the limit of no dimerization, as the results for clustered targets produce the same result as separated targets. Under the approximation of $[N]_{\text{eq}} = [N]_{\text{tot}}$ and $[S]_{\text{eq}} = [S]_{\text{tot}}$, we can approximate the dwell time, recruitment fraction, and target occupancy.

From Eq. 12, 17, 18, and 27, the dwell time reads,

$$\tau^{\text{DNA+clusTarg}} = \frac{[P]_{\text{eq}}(\chi_N + \chi_S + \gamma\chi_S\chi_N) + 2K_A^P[P]_{\text{eq}}^2\left(2\chi_N + \gamma\chi_N^2 + (2\chi_S + K_A^S C_0\chi_S)(1+\gamma\chi_N)^2\right)}{\left(4K_A^P[P]_{\text{eq}}^2 + [P]_{\text{eq}}\right)\left(k_{\text{off}}^N\chi_N + k_{\text{off}}^S\chi_S\right) + 2k_{\text{on}}^P[P]_{\text{eq}}^2(\chi_N + \chi_S + \gamma\chi_S\chi_N)} \tag{47}$$

From Eq. 19, 17, and 27, the recruitment fraction reads,

$$\theta^{\text{DNA+clusTarg}} = \frac{[P]_{\text{eq}}(\chi_N + \chi_S + \gamma\chi_S\chi_N) + 2K_A^P[P]_{\text{eq}}^2\left(2\chi_N + \gamma\chi_N^2 + (2\chi_S + K_A^S C_0\chi_S)(1+\gamma\chi_N)^2\right)}{[P]_{\text{tot}}} \tag{48}$$

From Eq. 21, and 27, the target occupancy reads,

$$\phi^{\text{DNA+clusTarg}} = \frac{[P]_{\text{eq}}\chi_S(1+\gamma\chi_N) + 2K_A^P[P]_{\text{eq}}^2(2\chi_S + K_A^S C_0\chi_S)(1+\gamma\chi_N)^2}{[S]_{\text{tot}}} \tag{49}$$

See section SXII.A for numerically solving these equations without approximations.

*Limit of irreversible protein-protein dimerization:*

For the mean dwell time of environment DNA + clusTarg, assuming $K_{\mathrm{A}}^{N}[N]_{\mathrm{eq}} = \chi_N$ and $K_{\mathrm{A}}^{S}[S]_{\mathrm{eq}} = \chi_S$, Eq. 48 can be simplified as,

$$\tau_{\mathrm{dimer,irr}}^{\mathrm{DNA+clusTarg}} \approx \frac{2\chi_N + \gamma\chi_N^2 + 2\chi_S(1+\gamma\chi_N)^2 + C_0 K_A^S \chi_S \left(1+\gamma\chi_N\right)^2}{2\left(k_{\mathrm{on}}^{N}[N]_0 + k_{\mathrm{on}}^{S}[S]_0\right)} \qquad (50)$$

When dimerization is very strong but still reversible, where $1 \ll K_{\mathrm{A}}^{PP}[P]_{\mathrm{tot}} < \infty$, the mean dwell time for DNA + targ is governed by 1) DNA-bound dimers, 2) the binding of in-solution monomers ($P$) to DNA-bound monomers ($PN$, $PS_2$, and $PS_2N$), and 3) the binding of in-solution dimers ($PP$) to DNA, $\tau_{\mathrm{rev-strong}}^{\mathrm{DNA+clusTarg}} = \frac{2([PPN]_{\mathrm{eq}}+[PPS_2]_{\mathrm{eq}}+[PPS_2N]_{\mathrm{eq}}+[PS_2PN]_{\mathrm{eq}}+[PNPN]_{\mathrm{eq}}+[PNPS_2N]_{\mathrm{eq}}+[PSNPSN]_{\mathrm{eq}})}{2\cdot 2[PP]_{\mathrm{eq}}(k_{\mathrm{on}}^{N}[N]_{\mathrm{eq}}+2k_{on}^{S}[S_2]_{\mathrm{eq}})+2[P]_{\mathrm{eq}}k_{\mathrm{on}}^{P}([PN]_{\mathrm{eq}}+[PS_2]_{\mathrm{eq}}+[PS_2N]_{\mathrm{eq}})}$. Assuming $K_{\mathrm{A}}^{N}[N]_{\mathrm{eq}} = \chi_N$ and $K_{\mathrm{A}}^{S}[S]_{\mathrm{eq}} = \chi_S$, using Eq. 27, it can be rewritten as,

$$\tau_{\mathrm{rev-strong}}^{\mathrm{DNA+targ}} \approx \frac{2\chi_N + 2\chi_S + 4\gamma\chi_S\chi_N + \gamma\chi_N^2 + 2\gamma^2\chi_N^2\chi_S + C_0 K_A^S \chi_S \left(1+\gamma\chi_N\right)^2}{2\left(k_{\mathrm{off}}^{N}\chi_N + k_{\mathrm{off}}^{S}\chi_S\right) + k_{\mathrm{off}}^{P}(\chi_N + \chi_S + \gamma\chi_N\chi_S)} \qquad (51')$$

The recruitment fraction is simplified from Eq. 49,

$$\theta_{\mathrm{dimer,irr}}^{\mathrm{DNA+clusTarg}} \approx \frac{2\chi_N + \gamma\chi_N^2 + 2\chi_S(1+\gamma\chi_N)^2 + C_0 K_A^S \chi_S \left(1+\gamma\chi_N\right)^2}{1 + 2\chi_N + \gamma\chi_N^2 + 2\chi_S(1+\gamma\chi_N)^2 + C_0 K_A^S \chi_S \left(1+\gamma\chi_N\right)^2}, \qquad (52)$$

Considering $[PP]_{\mathrm{eq}} = [PP]_{\mathrm{tot}}(1-\theta)$, the target occupancy is given by Eq. 50 and 52:

$$\phi_{\mathrm{dimer,irr}}^{\mathrm{DNA+clusTarg}} \approx \frac{2[1 + C_0 K_A^S]\chi_S(1+\gamma\chi_N)^2}{1 + 2\chi_N + \gamma\chi_N^2 + 2\chi_S(1+\gamma\chi_N)^2 + C_0 K_A^S \chi_S (1+\gamma\chi_N)^2} \cdot \frac{[PP]_{\mathrm{tot}}}{[S]_{\mathrm{tot}}}. \qquad (53)$$

### SVII. Inequalities between monomer and dimer metrics

#### SVII.A Comparing target occupancy for DNA+targ environment between monomer and irreversible dimer.

Here we compare the target occupancy for the separated targets under the assumption that target sites and nonspecific sites are in excess of proteins: $K_{\mathrm{A}}^{N}[N]_{\mathrm{eq}} = \chi_N$ and $K_{\mathrm{A}}^{S}[S]_{\mathrm{eq}} = \chi_S$. We can then see that the dimer occupancy is lower, or $\phi_{\mathrm{dimer,irr}}^{\mathrm{DNA+targ}} < \phi_{\mathrm{monomer}}^{\mathrm{DNA+targ}}$, from Eqs. 43 and 47, when, $\frac{(1+\gamma\chi_N)^2\chi_S}{1+2\chi_N+\gamma\chi_N^2+2\chi_S(1+\gamma\chi_N)^2} < \frac{(1+\gamma\chi_N)\chi_S}{1+\chi_N+\chi_S(1+\gamma\chi_N)}$,

which can be simplified to,

$$\chi_S > \frac{\chi_N(\gamma-1)}{(1+\gamma\chi_N)^2}. \qquad (54)$$

If the specific sites are not in excess ($[S]_{\mathrm{tot}} \sim [P]_{\mathrm{tot}}$), then we cannot assume the same value of $[S]_{\mathrm{eq}}$ for the monomer and dimer states, so the inequality does not simplify readily.

#### SVII.B Comparing dwell time for DNA+targ environment between monomer and reversible dimer.

Here we compare the dwell time for the separated targets under the assumption that target sites and nonspecific sites are in excess of proteins: $K_{\mathrm{A}}^{N}[N]_{\mathrm{eq}}=\chi_N$ and $K_{\mathrm{A}}^{S}[S]_{\mathrm{eq}}=\chi_S$. We can then see that the dimer dwell time is equal, or $\tau_{\mathrm{rev-strong}}^{\mathrm{DNA+targ}}=\tau_{\mathrm{monomer}}^{\mathrm{DNA+targ}}$ when $k_{\mathrm{off}}^{P}=k^*$, where the critical rate $k^*$ is given by Eqs. 40 and 45,

$$k^* \approx \frac{\left(k_{\mathrm{off}}^{N}\chi_N+k_{\mathrm{off}}^{S}\chi_S\right)\left(2\gamma\chi_N\chi_S+\gamma\chi_N^2+2(\gamma\chi_N)^2\chi_S\right)}{(\chi_N+\chi_S+\gamma\chi_N\chi_S)^2}. \tag{55}$$

Assuming $\chi_S \gg \chi_N$, we have,

$$k^* \approx \frac{2\gamma\chi_N\left(k_{\mathrm{off}}^{N}\chi_N+k_{\mathrm{off}}^{S}\chi_S\right)}{\chi_S(1+\gamma\chi_N)}. \tag{56}$$

With larger values of $k^*$, dimerization is more likely to enhance dwell time, because we will have $k_{\mathrm{off}}^{P}<k^*$. This occurs with larger values of $\gamma$, and with larger values of $k_{\mathrm{off}}^{N}$ or $k_{\mathrm{off}}^{S}$, considering $K_{\mathrm{A}}^{N}$ or $K_{\mathrm{A}}^{S}$ is constant.

Using the same method, we get the critical rate for environment $\mathrm{DNA}+\mathrm{clusTarg}$, given by Eqs. 40 and 45',

$$k^* \approx \frac{k_{\mathrm{off}}^{N}\chi_N+k_{\mathrm{off}}^{S}\chi_S}{(\chi_N+\chi_S+\gamma\chi_N\chi_S)^2}\left(2\gamma\chi_N\chi_S+\gamma\chi_N^2+2(\gamma\chi_N)^2\chi_S+C_0K_A^S\chi_S(1+\gamma\chi_N)^2\right). \tag{57'}$$

#### SVII.C Recruitment fraction is always enhanced by dimerization

The recruitment fractions tend to have relatively straightforward dependencies, since it does not depend on kinetics. For environment DNA, since $N$ sites are usually in excess and we can safely assume $[N]_{\mathrm{tot}}=[N]_{\mathrm{eq}}$, we can simply compare Eq. 33 and 34 while assuming $\chi_N$ is a constant in both systems. It derives,

$$\frac{\theta^{DNA}}{\theta_{\mathrm{monomer}}^{\mathrm{DNA}}}=\frac{1+\chi_N}{\chi_N}\frac{\chi_N+2K_A^P[P]_{\mathrm{eq}}(2\chi_N+\gamma{\chi_N}^2)}{\chi_N+1+2K_A^P[P]_{\mathrm{eq}}(2\chi_N+\gamma{\chi_N}^2+1)}\geq 1, \tag{58}$$

whose upper limit is given by Eq. 33 and 36,

$$\frac{\theta_{\mathrm{dimer,irr}}^{DNA}}{\theta_{\mathrm{monomer}}^{\mathrm{DNA}}}=\frac{(1+\chi_N)}{\chi_N}\frac{2\chi_N+\gamma{\chi_N}^2}{2\chi_N+\gamma{\chi_N}^2+1} \tag{59}$$

As noted in the main text, this enhancement always increases with larger $\gamma$ from taking the derivative, although when $\gamma\leq 2$, the dimensionality effect is too small to enhance protein dimers binding DNA. More generally, for any $0<K_{\mathrm{A}}^{P}<\infty$, $\theta$ is simply the average of $\theta_{\mathrm{dimer,irr}}$ and $\theta_{\mathrm{monomer}}$. Thus, as long as $\theta_{\mathrm{dimer,irr}}>\theta_{\mathrm{monomer}}$, we know $\theta\geq\theta_{\mathrm{monomer}}$. In this section, we prove $\theta_{\mathrm{dimer,irr}}>\theta_{\mathrm{monomer}}$ always hold for three environments.

To compare between $\theta^{\text{DNA}}_{\text{dimer,irr}}$ and $\theta^{\text{DNA}}_{\text{monomer}}$, we calculate the numerator in $\theta^{\text{DNA}}_{\text{dimer,irr}} - \theta^{\text{DNA}}_{\text{monomer}}$ from Eq. 34 and 36, which is $\Delta = 2K_{\text{A}}^{N}[N]_{\text{eq,dimer}} + \gamma\left(K_{\text{A}}^{N}[N]_{\text{eq,dimer}}\right)^2 - K_{\text{A}}^{N}[N]_{\text{eq,monomer}}$. To prove $\Delta > 0$, we prove $\Delta \leq 0$ is impossible. Suppose $\Delta \leq 0$, by definition of the recruitment fraction fewer proteins bound to the DNA, then $[N]_{\text{eq,dimer}} > [N]_{\text{eq,monomer}}$ since $[N]_{\text{tot}}$ is the same in both systems. However, $[N]_{\text{eq,dimer}} > [N]_{\text{eq,monomer}}$ yields $\Delta > 0$, which contradicts with the hypothesis $\Delta \leq 0$. Therefore, it must be $\Delta > 0$. Dimerization thus strictly enhances recruitment fraction to bare DNA.

The idea behind this derivation is that when $[N]_{\text{eq}}$ does not change, $\theta^{\text{DNA}}_{\text{dimer,irr}} > \theta^{\text{DNA}}_{\text{monomer}}$ means $\theta^{\text{DNA}}_{\text{dimer,irr}} > \theta^{\text{DNA}}_{\text{monomer}}$ no matter what $[N]_{\text{eq}}$ really is. Similarly, when target binding is considered, $\theta_{\text{dimer,irr}} > \theta_{\text{monomer}}$ with the same $[N]_{\text{eq}}$ and $[S]_{\text{eq}}$. Therefore, there must be $\theta_{\text{dimer,irr}} > \theta_{\text{monomer}}$.

For environment DNA + targ, the numerator of $\theta^{\text{DNA+targ}}_{\text{dimer,irr}} - \theta^{\text{DNA+targ}}_{\text{monomer}}$ is $\Delta = 2K_{\text{A}}^{N}[N]_{\text{eq,dimer}} + \gamma\left(K_{\text{A}}^{N}[N]_{\text{eq,dimer}}\right)^2 + 2K_{\text{A}}^{S}[S]_{\text{eq,dimer}}\left(1 + \gamma K_{\text{A}}^{N}[N]_{\text{eq,dimer}}\right)^2 - \left\{K_{\text{A}}^{N}[N]_{\text{eq,monomer}} + K_{\text{A}}^{S}[S]_{\text{eq,monomer}}\left(1 + \gamma K_{\text{A}}^{N}[N]_{\text{eq,monomer}}\right)\right\}$, given by Eqs. 42 and 46. Obviously, $\Delta > 0$ with the same $[S]_{\text{eq}}$ and $[N]_{\text{eq}}$. Then, we can prove $\Delta \leq 0$ is impossible. Suppose $\Delta \leq 0$, fewer proteins bound to the DNA means either $[N]_{\text{eq,dimer}} > [N]_{\text{eq,monomer}}$ (proteins removed from $N$ sites) or $[S]_{\text{eq,dimer}} > [S]_{\text{eq,monomer}}$ (proteins removed from $S$ sites. It is also possible to remove proteins from both $S$ and $N$ sites), since $[N]_{\text{tot}}$ and $[S]_{\text{tot}}$ are the same in both systems. Either of these inequality yields $\Delta > 0$, which contradicts with the hypothesis of $\Delta \leq 0$. Therefore, it must be $\theta^{\text{DNA+targ}}_{\text{dimer,irr}} > \theta^{\text{DNA+targ}}_{\text{monomer}}$.

For environment DNA + clusTarg, the numerator of $\theta^{\text{DNA+clusTarg}}_{\text{dimer,irr}} - \theta^{\text{DNA+clusTarg}}_{\text{monomer}}$ is $2K_{\text{A}}^{N}[N]_{\text{eq,dimer}} + \gamma\left(K_{\text{A}}^{N}[N]_{\text{eq,dimer}}\right)^2 + 2K_{\text{A}}^{S}[S]_{\text{eq,dimer}}\left(1 + \gamma K_{\text{A}}^{N}[N]_{\text{eq,dimer}}\right)^2 + C_0 K_{A}^{S} K_{\text{A}}^{S}[S]_{\text{eq,dimer}}\left(1 + \gamma K_{\text{A}}^{N}[N]_{\text{eq,dimer}}\right)^2 - \left\{K_{\text{A}}^{N}[N]_{\text{eq,mmonomer}} + K_{\text{A}}^{S}[S]_{\text{eq,mmonomer}}\left(1 + \gamma K_{\text{A}}^{N}[N]_{\text{eq,mmonomer}}\right)\right\}$ (given by Eq. 42 and 52), which is positive with the same $[N]_{\text{eq}}$ and $[S]_{\text{eq}}$ and thus $\theta^{\text{DNA+clusTarg}}_{\text{dimer,irr}} > \theta^{\text{DNA+clusTarg}}_{\text{monomer}}$.

#### SVIII Target binding without nonspecific interactions

Here we consider limiting behavior when nonspecific binding is eliminated, or $[N]_{\text{tot}} = 0$. For binding with pure clustered targets, the only DNA bound states are then: $[PS_2]$, $[PPS_2]$, and $[PSPS]$. The mean dwell time is:

$$\tau^{\text{clusTarg}} = \frac{1 + 2K_{\text{A}}^{P}[P]_{\text{eq}}\left(2 + C_0 K_{\text{A}}^{S}\right)}{2k_{\text{off}}^{P}K_{\text{A}}^{P}[P]_{\text{eq}} + k_{\text{off}}^{S}\left(1 + K_{\text{A}}^{S}[P]_{\text{eq}} + 4K_{\text{A}}^{P}[P]_{\text{eq}}\right)}. \tag{59}$$

Compared with the DNA environment, the extra term $k_{\text{off}}^{S}K_{\text{A}}^{S}[P]_{\text{eq}}$ comes from monomers dissociating from target in the reaction $PPS_2 \rightarrow P + PS_2$. When there is no dimerization, the dwell time is simply $1/k_{\text{off}}^{S}$. For irreversible dimers, the dwell time is,

$$\tau_{\text{dimer}}^{\text{clusTarg}} = \frac{1}{k_{\text{off}}^{S}}\left(1 + \frac{C_0 K_{\text{A}}^{S}}{2}\right). \tag{60}$$

When proteins form strong dimers but can still dissociate, the dwell time is,

$$\tau_{\text{rev-strong}}^{\text{clusTarg}} = \frac{1}{k_{\text{off}}^{S}} \cdot \frac{2 + C_0 K_{\text{A}}^{S}}{2 + k_{\text{off}}^{P}/k_{\text{off}}^{S}}. \tag{61}$$

The target occupancy is:

$$\phi^{\text{clusTarg}} = \frac{K_{\text{A}}^{S}[S]_{\text{eq}}}{[S]_{\text{tot}}} \cdot \left([P]_{\text{eq}} + 2K_{\text{A}}^{P}\left([P]_{\text{eq}}\right)^2 + 2C_0 K_{\text{A}}^{S}K_{\text{A}}^{P}\left([P]_{\text{eq}}\right)^2\right). \tag{62}$$

For monomers, the occupancy follows $\phi_{\text{monomer}}^{\text{clusTarg}} = \frac{K_{\text{A}}^{S}[S]_{\text{eq}}}{1+K_{\text{A}}^{S}[S]_{\text{eq}}} \cdot \frac{[P]_{\text{tot}}}{[S]_{\text{tot}}}$ and for irreversible dimers, the occupancy is,

$$\phi_{\text{dimer}}^{\text{clusTarg}} = \frac{K_{\text{A}}^{S}[S]_{\text{eq}} + K_{\text{A}}^{S}[S]_{\text{eq}}C_0 K_{\text{A}}^{S}}{1 + 2K_{\text{A}}^{S}[S]_{\text{eq}} + K_{\text{A}}^{S}[S]_{\text{eq}}C_0 K_{\text{A}}^{S}} \cdot \frac{[P]_{\text{tot}}}{[S]_{\text{tot}}}. \tag{63}$$

For binding with separated targets, considering bound states $[PS]$ and $[PPS]$, the mean dwell time is:

$$\tau^{\text{targ}} = \frac{1 + 4K_{\text{A}}^{P}[P]_{\text{eq}}}{2k_{\text{off}}^{P}K_{\text{A}}^{P}[P]_{\text{eq}} + k_{\text{off}}^{S}\left(1 + 4K_{\text{A}}^{P}[P]_{\text{eq}}\right)}. \tag{64}$$

For monomers and irreversible dimers, the dwell times are the same, $1/k_{\text{off}}^{S}$, since they can only bind one target. For strong but reversible dimers,

$$\tau_{\text{rev-strong}}^{\text{targ}} = \frac{1}{k_{\text{off}}^{S}} \cdot \frac{2}{2 + \frac{k_{\text{off}}^{P}}{k_{\text{off}}^{S}}}. \tag{65}$$

The target occupancy for this condition is:

$$\phi^{\text{targ}} = \frac{K_{\text{A}}^{S}[S]_{\text{eq}}}{[S]_{\text{tot}}} \cdot \left([P]_{\text{eq}} + 2K_{\text{A}}^{P}\left([P]_{\text{eq}}\right)^2\right). \tag{66}$$

The limits for monomers and irreversible dimers are,

$$\phi_{\text{monomer}}^{\text{targ}} = \frac{K_{\text{A}}^{S}[S]_{\text{eq}}}{1 + K_{\text{A}}^{S}[S]_{\text{eq}}} \cdot \frac{[P]_{\text{tot}}}{[S]_{\text{tot}}}, \tag{67}$$

and,

$$\phi_{\text{dimer}}^{\text{targ}} = \frac{K_A^S[S]_{\text{eq}}}{1 + 2K_A^S[S]_{\text{eq}}} \cdot \frac{[P]_{\text{tot}}}{[S]_{\text{tot}}}, \tag{68}$$

respectively.

#### SIX. Validation of the two-state dwell time formula with the MFPT calculation.

For a monomeric protein binding to DNA both specifically and nonspecifically, as illustrated in Fig. 1 of the main text, we derive the dwell time here using the survival time distribution $S(t)$, which describes the fraction of protein staying on DNA after binding at $t = 0$. This is also known as the mean first passage time (MFPT), or $\tau = \int_0^\infty t \frac{-dS(t)}{dt} dt$. For the differential, the initial condition is that the protein has just bound to the DNA from solution, and it survives until it returns to solution, so $S(0) = 1$ and $S(t \to \infty) = 0$, and we are ignoring rebinding events because they represent new events. Therefore, integration by parts gives $\tau = \int_0^\infty S(t)dt$. The survival probability can be related to the time-evolution of formation of $[P(t)]$, or free protein in solution, such that $S(t) = 1 - \frac{[P(t)]}{P_{\text{tot}}}$, where initially all proteins are DNA bound, $[P(0)] = 0$, and proteins cannot rebind to the DNA once they return to solution, so $[P(t \to \infty)] = P_{\text{tot}}$. Then $\frac{-dS(t)}{dt} = \frac{1}{P_{\text{tot}}} \frac{d[P(t)]}{dt}$. The system of coupled ODEs defining the time evolution of $\frac{d[PN(t)]}{dt}$, $\frac{d[PS(t)]}{dt}$ and $\frac{d[PSN(t)]}{dt}$ as defined by the states illustrated in Figure 1 (excluding protein binding to DNA) is a nonlinear system of the unknowns $[PS(t)]$, $[PN(t)]$, $[PSN(t)]$, $[N(t)]$, and $[S(t)]$, due to bimolecular reactions. Specifically,

$$\begin{aligned} \frac{d[PN(t)]}{dt} &= -\gamma k_{\text{on}}^S[S][PN(t)] - k_{\text{off}}^N[PN(t)] + k_{\text{off}}^S[PSN(t)] \\ \frac{d[PS(t)]}{dt} &= -\gamma k_{\text{on}}^N[N][PS(t)] - k_{\text{off}}^S[PS(t)] + k_{\text{off}}^N[PSN(t)] \\ \frac{d[PSN(t)]}{dt} &= \gamma k_{\text{on}}^S[S][PN(t)] + \gamma k_{\text{on}}^N[N][PS(t)] - (k_{\text{off}}^N + k_{\text{off}}^S)[PNS(t)] \end{aligned} \tag{69}$$

We compute $\frac{d[P(t)]}{dt}$ either from $\frac{d[P(t)]}{dt} = k_{\text{off}}^S[PS(t)] + k_{\text{off}}^N[PN(t)]$, or mass conservation, $P_{tot} = [P(t)] + [PS(t)] + [PN(t)] + [PSN(t)]$.

To solve, we assume an equilibrium steady-state for free DNA sites $[S]$ and $[N]$: they are not time-dependent, e.g. $[N] = [N]_{eq}$, then the system of ODEs becomes linear. We can then directly solve for all unknowns including $[P(t)]$ as a function of all variables and rate constants from the eigenvalue problem, and therefore solve for $\tau$ using $\tau = \int_0^\infty (1 - \frac{[P(t)]}{P_{\text{tot}}}) dt$. We solve for $[P(t)]$ based on the two different initial conditions possible for a protein that just bound the DNA, either $[PS, PN, PSN] = [1,0,0]$ or $[PS, PN, PSN] = [0,1,0]$. The corresponding dwell times $\tau_{PS,0}$ and $\tau_{PN,0}$ are given by:

$$\tau_{PN,0} = \frac{\left(\gamma k_{\text{on}}^{S}[S]_{\text{eq}} + k_{\text{off}}^{S}\right)\left(k_{\text{off}}^{N} + k_{\text{off}}^{S} + \gamma k_{\text{on}}^{N}[N]_{\text{eq}}\right)}{k_{\text{off}}^{S}k_{\text{off}}^{N}\left(\gamma k_{\text{on}}^{S}[S]_{\text{eq}} + k_{\text{off}}^{N} + k_{\text{off}}^{S} + \gamma k_{\text{on}}^{N}[N]_{\text{eq}}\right)} \tag{70}$$

$$\tau_{PS,0} = \frac{\left(\gamma k_{\text{on}}^{N}[N]_{\text{eq}} + k_{\text{off}}^{N}\right)\left(k_{\text{off}}^{N} + k_{\text{off}}^{S} + \gamma k_{\text{on}}^{S}[S]_{\text{eq}}\right)}{k_{\text{off}}^{S}k_{\text{off}}^{N}\left(\gamma k_{\text{on}}^{S}[S]_{\text{eq}} + k_{\text{off}}^{N} + k_{\text{off}}^{S} + \gamma k_{\text{on}}^{N}[N]_{\text{eq}}\right)}$$

And the total dwell time is weighted by the flux into either the $PS\ or\ PN$ starting state, since one cannot enter $PSN$ directly from solution, $p = k_{\text{on}}^{N}[N]_{\text{eq}}/\left(k_{\text{on}}^{N}[N]_{\text{eq}} + k_{\text{on}}^{S}[S]_{\text{eq}}\right)$,

$$\tau_{\text{monomer}}^{\text{DNA+targ}} = p\tau_{PN,0} + (1-p)\tau_{PS,0}. \tag{71}$$

It can be simplified as,

$$\tau_{\text{monomer}}^{\text{DNA+targ}} = \frac{K_{\text{A}}^{N}[N]_{\text{eq}} + K_{\text{A}}^{S}[S]_{\text{eq}} + \gamma K_{\text{A}}^{N}[N]_{\text{eq}}K_{\text{A}}^{S}[S]_{\text{eq}}}{k_{\text{on}}^{N}[N]_{\text{eq}} + k_{\text{on}}^{S}[S]_{\text{eq}}}. \tag{72}$$

which equals Eq. SVI.B.1.1. To conclude, the MFPT calculated here from the survival time distribution is identical to the dwell time calculated from the two-state model of Eq. SVI.B.1.1. The survival time distribution contains more detail than the MFPT, as it predicts the full time-dependence. This makes it more challenging to calculate, and if the number of unknowns in the linear system produces an eigenvalue equation that is higher than 5$^{\text{th}}$ order, the eigenvalues must be solved for numerically, whereas Eq. SVI.B.1.1 can always be written down symbolically.

### SX. Parameter Range Justifications

#### SX.A Species concentration in each microenvironment

We have broken up a nucleus into possible microenvironments to solve for the equilibrium steady-state behavior in each of them separately (first). Thus, for any species $A$, we use local concentrations for all models, namely $[N] = N/\left(\frac{V}{L}\cdot l\right)$. For $N$ sites, the local concentration is the same as the global concentration in the nucleus, since $N \propto l$. However, in our model, $S$ specific sites vary by segment (from 0 to 4), thus the local concentration $[S]$ in the microenvironment need not equal the global concentration across the genome.

#### SX.B Estimate volume-to-length ratio in *D. Melanogaster*

Most of the nuclei of *D. Melanogaster* is smaller than $V{\sim}25 \times 10^{9}\ \text{nm}^3$ [9]. Their genome has about 120 Mb euchromatin that is accessible to all types of transcription factors [10]. Since 3 base pairs is 1 nm long, the contour length of the DNA in euchromatin is $L{\sim}\frac{120\ Mb}{2}\cdot\frac{1nm}{3bp} = 2\times 10^{7}\ \text{nm}$. Thus, we have $V/L \approx 1000\ \mathrm{nm}^2$. We always keep this ratio a constant in our model such that the volume of our models is $(V/L) \cdot l$, where $l$ is the length of the DNA segment. According to $V/L$, we chose $h^2$ from $1\ \mathrm{nm}^2$ to $1000\ \mathrm{nm}^2$ such that $\gamma = \frac{V}{L}/h^2$ ranges from 1 to 1000, which means dimensional reduction enhances (or if $\gamma$=1 does not change) each pairwise reaction to form contacts.

#### SX.C Estimate length of DNA segments between two nucleosomes

For most eukaryotic cells, the average nucleosome spacing is about 190 bp where 147 bp wrap around nucleosomes[11]. This results in around $\frac{L}{190\ \mathrm{nm}} = \frac{2\times10^7}{190} \approx 10^5$ continuous DNA segments in *Drosophila* genome. Since the entry and exit 10 bp of nucleosomes are available to proteins[12], we assert 63 bp (21 nm) as the shortest length of DNA segments that is accessible to proteins. The space between two nucleosomes can be as long as thousands bp[13]. However, since the persistence length of ds-DNA is about 50 nm [14], we limit the length of DNA segment to be less than 500 nm.

#### SX.D Definition of nonspecific sites and the number of nonspecific $N$ and $S$ sites

The total number of sites (both $N$ and $S$) on a DNA segment is the maximum number of proteins that can bind to this DNA. Suppose a DNA has $N_{\mathrm{bp}}$ basepairs and the footprint length (or the sequence length of targets) of the protein is $l_{\mathrm{fp}}$. Then, the maximum number of proteins that can bind is $\lfloor N_{\mathrm{bp}}/l_{\mathrm{fp}} \rfloor$ (Fig-S1). Here we take GAF as an example. Its DNA binding domain is a zinc finger and binds to GAGAG motifs on DNA [15] and thus $l_{\mathrm{fp}} = 5$. When a DNA has $N_{\mathrm{bp}} = 63$, $\lfloor N_{\mathrm{bp}}/l_{\mathrm{fp}} \rfloor = 12$. For environment $\mathrm{DNA}$, it means that there are 12 $N$ sites. For environment $\mathrm{DNA + targ}$ and $\mathrm{DNA + clusTarg}$, for the purpose of dimer binding, we assume there are always 2 $S$ sites, and there are 10 $N$ sites.

#### SX.E Kinetic rates and association constants for pairwise reactions between protein and DNA

To maintain consistency with the kinetics of GAF, we use values representative of GAF. Based on the searching time $\tau_{\mathrm{search}} = 150\ \mathrm{s}$ and copy numbers (14288 targets for GAF) given in Tang et al. [15], the apparent association rate for targets is in the scale of $10^4\ \mathrm{nm}^3/s$. However, since facilitated diffusion can enhance the association rate by several magnitudes [1], here we assume $k_{\mathrm{on}}^{S} = 10^3\ \mathrm{nm}^3/s = 602.2\ (\mathrm{M}\cdot\mathrm{s})^{-1}$. Shown in *in-vitro* experiments.[16], the apparent binding frequency to specific sites is about 10-fold as to nonspecific sites. Also to approximately decouple the effect of facilitated diffusion, here we assume $k_{\mathrm{on}}^{N} = 0.2 \cdot k_{\mathrm{on}}^{S} = 120.44\ (\mathrm{M}\cdot\mathrm{s})^{-1}$. The *in-vivo* measurement finds a transient dwell time of 3.7 s and a stable dwell time of 130 s [15], according to which we assume the lower limit of $k_{\mathrm{off}}^{N}$ is in the scale of seconds and assume $k_{\mathrm{off}}^{S} < 0.1 \cdot k_{\mathrm{off}}^{N}$. Marklund et al. [17] showed that the sequence specificity of protein-DNA binding is mainly influenced by association rate and the association constants varied by up to two orders of magnitude. Therefore, we assume the association rates are constants and dissociation rates can change as salt concentration changes and we vary $K_{\mathrm{A}}^{N}$ from $1\ \mathrm{M}^{-1}$ to $10^3\ \mathrm{M}^{-1}$. For $K_{\mathrm{A}}^{S}$, since Le et al. [18] reported that sequence variation can result in dissociation constant change of up to 3 magnitudes, we set $K_{\mathrm{A}}^{S} \le 1000 \cdot K_{\mathrm{A}}^{N}$.

#### SX.F Dynamical parameters and association constants for protein dimerization

Both the association rate and dissociation rate of protein dimerization can vary substantially. For simplicity, we always fix the dissociation rate $k_{\rm off}^{P}$ and change the association rate $k_{\rm on}^{P}$ according to different association constant $K_{\rm A}^{P}$. We vary $K_{\rm A}^{P}$ from 0 for monomers to $10^9\ {\rm M}^{-1}$ for strong dimers and to ∞ for obligate dimers. The 3D and 1D diffusion constants are $D_{\rm 3d} = 1.5\ {\rm \mu m}^2/s$ in solution and $D_{\rm 1d} = 0.6\ {\rm \mu m}^2/s$ on DNA representing the dynamics of GAF [15, 16].

#### SX.G The local concentration of proteins

For GAF, the concentration of proteins in the nucleus is in the scale of $1\ {\rm \mu M}$ (56683 monomers [15]). However, in each microenvironment, we consider local variations of equilibrium protein concentrations are possible. With $[P]_{\rm tot} = 7.908\ {\rm \mu M}$ and the length of DNA is $l = 21\ {\rm nm}$, there are 0.2 protein copies in the micro environment. When targets are present and proteins are fixed on DNA, other in-solution proteins can diffuse to fill the "depleted" region and raise the local concentration. With 2 targets on DNA, in some cases we will therefore assume that 2 protein copies are present, which gives $[P]_{\rm tot} = 158.15\ {\rm \mu M}$ with $l = 21\ {\rm nm}$. Although this concentration is much higher than the overall nucleic concentration of any transcription factors, at the nonequilibrium steady-state of the nucleus, it could reach this high due to the inhomogeneities we are studying here.

### SXI. Theoretical extensions to measure selectivity arising from competition between multiple microenvironments.

To calculate the equilibrium for studying the selectivity between adjacent DNA segments with different environments, we implement an algorithm to solve two ODE systems together while keeping the concentration of proteins in solution ($[P]_{\rm eq} + [PP]_{\rm eq}$) to be the same. Denote two environments as $E_1$ and $E_2$, and the total numbers of proteins at equilibrium are $N_P^1$ and $N_P^2$, respectively. The total number in the whole system is $N_P^{\rm tot} = N_P^1 + N_P^2$. Suppose the recruitment fraction in environment $E$ with total number of proteins $N_P$ is $\theta(E, N_P)$, which can be numerically solved. Then, for $E_1$ and $E_2$, the numbers of proteins in solution are $N_P^1(1 - \theta(E_1, N_P^1)\ )$ and $N_P^2(1 - \theta(E_2, N_P^2))$, respectively. Now, the equilibrium solution for these connected environments is,

$$\begin{cases} N_P^1\left(1 - \theta(E_1, N_P^1)\right) = N_P^2\left(1 - \theta(E_2, N_P^2)\right) \\ N_P^1 + N_P^2 = N_P^{\rm tot} \end{cases}, \tag{73}$$

where $N_P^{\rm tot}$, $E_1$, and $E_2$ are given by user. The selectivity for $E_1$ is defined as,

$$\text{Selectivity}_1 = \frac{N_P^1 \cdot \theta(E_1, N_P^1)}{N_P^2 \cdot \theta(E_2, N_P^2)}. \tag{74}$$

To numerically calculate the selectivity in Fig. 7b, the parameters used for $N_P^{\rm tot} = 2$ ($[P]_{\rm tot} = 79.075\ {\rm \mu M}$) and $N_P^{\rm tot} = 4$ ($[P]_{\rm tot} = 158.15\ {\rm \mu M}$) are listed in Dataset S1.

To calculate the partitioning of proteins in Fig. S10-f,g, this method is generalized to include the population of DNA segments. Denote $N_{\mathrm{DNA}}^{i}$ as the population of DNA segments in environment $E_i$. Then, Eq. 73 becomes,

$$\begin{cases} N_P^i\left(1-\theta\left(E_i, N_P^i\right)\right) = const. \\ \sum_{i\in\{\mathrm{Environments}\}} N_{\mathrm{DNA}}^{i} N_P^i = N_P^{\mathrm{tot}}, \end{cases} \tag{75}$$

The parameters used are listed in Dataset S1.

Here we analytically estimate the partitioning of proteins between DNA with single targets (as $E_1$) and DNA with two-target clusters (as $E_2$). Eq. 75 gives,

$$\begin{cases} N_P^1\left(1-\theta(E_1, N_P^1)\right) = N_P^2\left(1-\theta(E_2, N_P^2)\right) \\ N_{\mathrm{DNA}}^{1} N_P^1 + N_{\mathrm{DNA}}^{2} N_P^2 = N_P^{\mathrm{tot}} \end{cases} \tag{76}$$

and then the number of proteins bound at equilibrium is,

$$\begin{cases} N_P^1\theta(E_1, N_P^1) = \dfrac{N_P^{\mathrm{tot}}}{N_{\mathrm{DNA}}^{1}+N_{\mathrm{DNA}}^{2}} \cdot \dfrac{\theta(E_1, N_P^1)\left(1-\theta(E_2, N_P^2)\right)}{2-\theta(E_1, N_P^1)-\theta(E_2, N_P^2)} \\ N_P^2\theta(E_2, N_P^2) = \dfrac{N_P^{\mathrm{tot}}}{N_{\mathrm{DNA}}^{1}+N_{\mathrm{DNA}}^{2}} \cdot \dfrac{\theta(E_2, N_P^2)\left(1-\theta(E_1, N_P^1)\right)}{2-\theta(E_1, N_P^1)-\theta(E_2, N_P^2)} \end{cases} \tag{77}$$

and the selectivity is (using Eq. 74),

$$\mathrm{Selectivity}_2 = \frac{\theta(E_2, N_P^2)\left(1-\theta(E_1, N_P^1)\right)}{\theta(E_1, N_P^1)\left(1-\theta(E_2, N_P^2)\right)} \tag{78}$$

For monomers, Eq. 42 gives the recruitment fraction, $\frac{\chi_N+\chi_S(1+\gamma\chi_N)}{1+\chi_N+\chi_S(1+\gamma\chi_N)}$. Assuming target binding is much stronger than nonspecific binding, namely $\chi_S(1+\gamma\chi_N) \gg \chi_N$, the recruitment fraction derives $\frac{\chi_S(1+\gamma\chi_N)}{1+\chi_S(1+\gamma\chi_N)}$. Since there are two S sites in $E_2$ and 1 S site in $E_1$, we have and denote $\chi_S^{E_2} = \chi_S^{E_1} = \chi_S$. Assuming and denoting $\chi_N^{E_2} = \chi_N^{E_1} = \chi_N$ and noting that $\gamma$ should be the same for two environments, we have,

$$\begin{cases} \theta(E_1, N_P^1) = \dfrac{\chi_S(1+\gamma\chi_N)}{1+\chi_S(1+\gamma\chi_N)} \\ \theta(E_2, N_P^2) = \dfrac{2\chi_S(1+\gamma\chi_N)}{1+2\chi_S(1+\gamma\chi_N)} \end{cases}. \tag{79}$$

Therefore, like Eq. 77, the per-target bound proteins are the same (However, one still has $\mathrm{Selectivity}_2 = 2$ by Eq. 78), $\frac{N_P^1\theta(E_1, N_P^1)}{1} = \frac{N_P^2\theta(E_2, N_P^2)}{2}$. For irreversible dimers, adopt the assumptions for $\chi_S^{E_2} = \chi_S^{E_1} = \chi_S$ and $\chi_N^{E_2} = \chi_N^{E_1} = \chi_N$, Eq. 46 (for $E_1$) and Eq. 52 (for $E_2$) derive,

$$\begin{cases} \theta(E_1, N_P^1) = \dfrac{2\chi_S(1+\gamma\chi_N)^2}{1+2\chi_S(1+\gamma\chi_N)^2} \\ \theta(E_2, N_P^2) = \dfrac{2\chi_S(1+\gamma\chi_N)^2 + C_0 K_A^S \chi_S\left(1+\gamma\chi_N\right)^2}{1+2\chi_S(1+\gamma\chi_N)^2 + C_0 K_A^S \chi_S\left(1+\gamma\chi_N\right)^2} \end{cases}. \tag{80}$$

Plugging this into Eq. 78 gives the maximum selectivity for $E_2$,

$$\text{Selectivity}_2 = 1 + \frac{C_0 K_A^S \chi_S}{2}. \tag{81}$$

When dimerization is reversible, a higher local concentration results in a higher dimer-to-monomer ratio at equilibrium and shift the selectivity from the lower limit of *2* to the upper limit of $1 + C_0 K_A^S \chi_S / 2$.

### SXII. Numerical Methods

#### SXII.A Solving rate equations for equilibria.

The true equilibria without approximation are obtained by numerically solving ODEs (Eqs. SV.B.2, SV.C.2, and SV.D.2) using the SciPy package solve_ivp with the 'LSODA' integration routines[19] in Python3. For systems that considers monomers or reversible dimers ($K_{\text{eq}}^P < \infty$), initial conditions are $[P] = [P]_{\text{tot}}$, $[S] = [S]_{\text{tot}}$, and $[N] = [N]_{\text{tot}}$. For systems considering irreversible dimers, initial conditions use $[PP] = [P]_{\text{tot}}/2$. Note that for environment $\text{DNA} + \text{clusTarg}$, targets are considered as clusters ($[S_2] = [S]/2$), thus the initial condition use $[S_2] = [S]_{\text{tot}}/2$. The ODEs are solved from time 0 to $t_e$. The concentrations at $t_e$ is considered as at equilibrium. Instead of using a fixed large, $t_e$, we adjust $t_e$ by $\boldsymbol{x}(t_e)$ (as defined in SV.A.1). We enlarge $t_e$ until $\boldsymbol{x}(t_e)$ satisfies two equilibrium conditions:

1. $\boldsymbol{M} \cdot \boldsymbol{p}(\boldsymbol{x}(t_e)) < \epsilon \cdot \boldsymbol{p}(\boldsymbol{x}(t_e))$ (as defined in SV.A.1): The relative change of each species is smaller than $\epsilon$. We use $\epsilon = 0.01$ for environment $\text{DNA} + \text{clusTarg}$ and $\epsilon = 0.001$ for environments DNA and $\text{DNA} + \text{targ}$.

2. $\left|K_{\text{A}}^{obs}/K_{\text{A}} - 1\right| < \epsilon$: for all reactions: Each reaction is at equilibrium within error of $\epsilon$ (same as in item condition 1). Observed equilibrium ($K_{\text{eq}}^{obs}$) are calculated for each reaction (Using Table S2, S3, and S4, for a reaction $A + B \leftrightharpoons C$, $K_{\text{A}}^{obs} = [C]/[A][B]$) and compared with true association constants ($K_{\text{A}}$'s listed in Table S2, S3, and S4).

The equilibria are then used to calculate dwell times (SIV.A), recruitment fractions (SIV.B), and target occupancies (SIV.C). Codes are available on GitHub (https://github.com/sangmk/dimerEnhanceProteinDNA). The parameters used for each plot are listed SI Dataset S1: "Parameters_rateEquations.xlsx".

#### SXII.B. Gillespie simulation

We use the algorithm described by Gillespie 1976 [20, 21] to simulate the stochastic dynamics of our reaction network. We use the algorithm described by Gillespie 1976 [20, 21] to simulate the stochastic dynamics of our reaction network. The propensity functions are described in Table S2, S3, and S4. To calculate the dwell time, we labeled all proteins such that the binding condition of each protein at each timepoint is tracked. For better sampling, we terminate simulations when each protein has at least 10 dissociation events from DNA and more than $1000 \cdot N_P$ total dissociation events happen where $N_P$ is the number of protein monomers. Each simulation is initialized at equilibrium. The numbers of molecules in each species (protein in all binding states, S sites, and N sites) are rounded to the nearest integer by numpy.round()[22], determined by equilibrium concentration with the constraint of $N_P = 200$. $N_P = 200$ is ensured by the number of protein monomers. After rounding the number of all species and adjusting the number of protein monomers without having negative values, we accept $N_P$ even when $N_P > 200$. Codes and parameters used for each simulation are available on GitHub (https://github.com/sangmk/dimerEnhanceProteinDNA).

#### SXII.C Single-particle spatiotemporal stochastic simulations (NERDSS)

We use the nonequilibrium reaction-diffusion self-assembly simulator (NERDSS)[23] to perform particle-based stochastic simulations, available open source at https://github.com/mjohn218/NERDSS. Proteins are designed with three interfaces, one to form protein dimers and one each to bind DNA nonspecifically and/or specifically. Thus, as in Figure 1, all sites can bind simultaneously. In environment $\mathrm{DNA} + \mathrm{clusTarg}$ Two clustered targets ($S$ molecules) are bound together to enable the loop closure reaction, which maps a bimolecular reaction to a co-localized unimolecular reaction while preserving detailed balance. We consider following molecules and reactions to build our system:

Molecules:

- $S$: Specific binding sites.
- $N$: Nonspecific binding sites.
- $P$: Proteins.
- $nuc$: Nucleosomes.

We set $S$ and $nuc$ as stationary molecules with $D = 0$. $N$ molecules move along the 1-D DNA with $D_N = 1.0\ \mu\mathrm{m}^2/s$. $P$ molecules move in the 3-D solution with $D_P = 1.5\ \mu\mathrm{m}^2/s$.

When $P$ and $N$ associates, the protein moves along the 1-D DNA, and the complex $PN$ has diffusion constant $D_{PN} = (D_P^{-1} + D_N^{-1})^{-1} = 0.6\ \mu\text{m}^2/\text{s}$. The conversion from 3D macroscopic rates to 3D microscopic kinetic rates can be found in Fu's work [24].

##### SXII.D Cross validation between spatial simulations and nonspatial solutions.

For Figure S2, each condition reports an average over 48 trajectories. 20 proteins are initialized randomly in the system and are let to bind on DNA. Then, simulations are restarted. Molecules and kinetic rates are set up as described in SXII.C. Parameters are listed in table S5.

##### SXII.E Selectivity between two DNA segments with $\text{DNA} + \text{clusTarg}$ and $\text{DNA} + \text{targ}$ separated by a nucleosome.

As shown in Figure 7a, DNA is a straight fiber along the x-axis. One nucleosome (nucleosomes have 5.5 nm radius) is placed in the middle of DNA fiber to block 1D diffusions and two nucleosomes are placed at two ends of the DNA fiber to make the whole system periodic. Proteins are initialized in solution randomly. For each condition, 48 trajectories were simulated with wall time 2000 s. To avoid correlation between data points, the coordinates of proteins were collected every 60 s. The burn-in time is 120 s. Standard errors are calculated by 100 bootstrap samplings. Molecules and kinetic rates are set up as described in SXII.C. Proteins are considered bound to DNA when its center of mass is within 5.5 nm of the DNA fiber. See Table S6 for parameters.

##### SXII.F Partitioning of proteins among four DNA molecules with 1~4 clustered targets.

As shown in Figure S10a, four DNA molecules are placed at four 1/4 centers in a water box. Four proteins are initialized in the solution randomly. Four DNA molecules have 1 to 4 targets clustered at the center. For monomer and dimer simulations, the protein structure is the same as above. For the linear tetramer structure, an additional protein-protein binding domain is added such that one protein can bind two other proteins (Figure S10b). Since there are only four proteins in the system, the largest multimer is tetramer. There is no nucleosome in the system. 30 trajectories were simulated with wall time 2000 s. To avoid correlation between data points, the coordinates of proteins were collected every 80 s. The burn-in time is 20 s. Standard errors are calculated by 100 bootstrap samplings over all trajectories. Molecules and kinetic rates are set up as described in SXII.C. Proteins are considered bound to DNA when its center of mass is within 5.5 nm of the DNA fiber. See Table S7 for parameters.

#### SXIII. The distribution of the size of target clusters

We describe here how the CHiP-seq data on GAF occupancy is extracted for Figure 5d in the main text. The ChIP-seq peaks and intensities for GAF are obtained from NCBI GSE58957 [25], Table S3 of the publication, filtered for high confidence peaks whose "Jaspar_hcGAF=Yes". These peaks were grouped into intensities based on the GAF motif clusters in Fig 5 of a previous publication [15]. We performed the same grouping here and added in an analysis for error bars. We briefly describe here the protocol for establishing the motif cluster sizes. For each peak in the CHiP-Seq data, we count how many GAGAG motifs are within the sequence. Only full-length motifs are considered as targets. For example, GAF's full sequence motif is GAGAG, therefore both GAGAG and GAGAGAG have one target and GAGAGAGAGAG has two targets. This is consistent with the space taken up by a dimer of proteins that bind to the targets, which requires two full motifs. So for each peak intensity, we know the number of target motifs. To calculate the standard error on the median intensity for each cluster size *n,* we perform bootstrapping. We sample with replacement from the distribution of intensities for each cluster size. We repeat this 100 times to get a standard error on the intensity. For reference, for a cluster size of 6 there are 303 peak intensities in the dataset, and for a cluster size of 1 there are 505 peaks in the dataset.

To compute the relative frequency of all GAGAG clusters across the genome, as plotted in Fig S10, we do not use the CHiP-seq data. Here, we instead calculate the number *n* of GAGAG motifs that are between two adjacent nucleosomes, and assign this to be a cluster of size *n*. Even if there is some separation between the motifs, if they are between adjacent nucleosomes we assume they are part of a single cluster. This is because GAF has intrinsically disordered regions that can support protein-protein interactions across several or even tens of nm [26]. Considering that the edge of nucleosomes are accessible for proteins dynamically [16], we use the center of each nucleosome as the barrier. The nucleosome positions are obtained from NucMap [27] dmNuc0210501 [13]. The GAGAG motif positions within the drosophila genome are located by using the software tool Integrative Genomics Viewer (IGV_2.18.4) [28].

### Table S1 Parameter range used to represent physiological conditions

From Fig.1 to Fig. 6, Fig. S2, Fig. S6 to Fig. S8, and Fig. S10, we use parameters in physiological regions discussed above (Table S1). However, limited by the timestep of single-particle reaction-diffusion simulations, we use higher rates for protein-DNA reactions such that proteins can dynamically explore among DNA sites. For Fig 7 and Fig S10, these rates are magnified: $k_{\mathrm{on}}^{N} = 12044\ \mathrm{M}^{-1}s^{-1}$, $k_{\mathrm{off}}^{N} = 1204.4\ s^{-1}$, $k_{\mathrm{on}}^{S} = 60220\ \mathrm{M}^{-1}s^{-1}$, and $k_{\mathrm{off}}^{S} = 6.022\ s^{-1}$.

| Parameter | Unit | Range or value | Parameter | Unit | Range or value |
|---|---|---|---|---|---|
| $K_{\mathrm{A}}^{N}$ | $\mathrm{M}^{-1}$ | $1\text{~}2 \times 10^{3}$ | $K_{\mathrm{A}}^{S}$ | $\mathrm{M}^{-1}$ | $1 \times 10^{3}\text{~}2 \times 10^{6}$ |
| $k_{\mathrm{on}}^{N}$ | $\mathrm{M}^{-1}s^{-1}$ | 120.4 | $k_{\mathrm{on}}^{S}$ | $\mathrm{M}^{-1}s^{-1}$ | 602.2 |
| $k_{\mathrm{off}}^{N}$ | $s^{-1}$ | 0.120~120.4 | $k_{\mathrm{off}}^{S}$ | $s^{-1}$ | $3.01 \times 10^{-4}$<br>$\text{~}6.02 \times 10^{-2}$ |
| $[N]_{\mathrm{tot}}$ | μM | 948.9 (0 target)<br>790.8 (2 targets) | $[S]_{\mathrm{tot}}$ | μM | 0 (0 target)<br>158.2 (2 targets) |
| $\chi_N$ | 1 | $1.6 \times 10^{-2}\text{~}1.9$ | $\chi_S$ | 1 | $0.16\text{~}3.2 \times 10^{2}$ |
| $K_{\mathrm{A}}^{P}$ | $\mathrm{M}^{-1}$ | 0, ∞,<br>$10^{1}\text{~}10^{9}$ | $[P]_{\mathrm{tot}}$ | μM | $1.6\text{~}1.6 \times 10^{4}$ |
| $k_{\mathrm{on}}^{P}$ | $\mathrm{M}^{-1}s^{-1}$ | $10^{2}\text{~}10^{7}$ | $K_{\mathrm{A}}^{P}[P]_{\mathrm{tot}}$ | 1 | 0, ∞,<br>$1.6\times 10^{1}\text{~}1.6 \times 10^{9}$ |
| $k_{\mathrm{off}}^{P}$ | $s^{-1}$ | $10^{-3}\text{~}10^{1}$ | $\gamma$ | 1 | 1~1000 |

*Table S1.*

### Table S2 Reactions with their association constants and propensities for environment DNA

1D reactions are colored red and their $K_{\mathrm{A}}$'s follow the definition of Eq. SI. 5.

| Reaction | $K_A$ | Forward reaction propensity / Backward reaction propensity |
|---|---|---|
| $P + P \leftrightharpoons PP$ | $K_A^P$ | $p_{P+P\to PP} = k_{\mathrm{on}}^P[P]^2$ |
| | | $p_{PP\to P+P} = k_{\mathrm{off}}^P[PP]$ |
| $PN + P \leftrightharpoons PPN$ | $2K_A^P$ | $p_{P+PN\to PPN} = 2k_{\mathrm{on}}^P[P][PN]$ |
| | | $p_{PPN\to P+PN} = 2k_{\mathrm{off}}^P[PPN]$ |
| $P + N \leftrightharpoons PN$ | $K_A^N$ | $p_{P+N\to PN} = k_{\mathrm{on}}^N[P][N]$ |
| | | $p_{PN\to P+N} = k_{\mathrm{off}}^N[PN]$ |
| $PP + N \leftrightharpoons PPN$ | $2K_A^N$ | $p_{PP+N\to PPN} = 2k_{\mathrm{on}}^N[PP][N]$ |
| | | $p_{PPN\to PP+N} = k_{\mathrm{off}}^N[PPN]$ |
| $PPN + N \leftrightharpoons PNPN$ | $\gamma \frac{K_A^N}{2}$ | $p_{PPN+N\to NPPN} = \gamma k_{\mathrm{on}}^N[PPN][N]$ |
| | | $p_{NPPN\to PPN+N} = 2k_{\mathrm{off}}^N[NPPN]$ |
| $PN + PN \leftrightharpoons PNPN$ | $\gamma K_A^P$ | $p_{PN+PN\to NPPN} = \gamma k_{\mathrm{on}}^P[PN][PN]$ |
| | | $p_{NPPN\to PN+PN} = k_{\mathrm{off}}^P[NPPN]$ |

*Table S2.*

### Table S3 Reactions with their association constants and propensities for environment DNA + targ

1D reactions are colored red and their $K_{\mathrm{A}}$'s follow the definition of Eq. SI. 5. $PNPS$ and $PPSN$ are two species depending on whether two proteins form contact with DNA or one protein is co-localized to DNA.

| Reaction | $K_A$ | Forward reaction propensity<br>Backward reaction propensity |
|---|---|---|
| $P + P \leftrightharpoons PP$ | $K_A^P$ | $p_{P+P\to PP} = k_{\mathrm{on}}^P[P]^2$<br>$p_{PP\to P+P} = k_{\mathrm{off}}^P[PP]$ |
| $PN + P \leftrightharpoons PPN$ | $2K_A^P$ | $p_{P+PN\to PPN} = 2k_{\mathrm{on}}^P[P][PN]$<br>$p_{PPN\to P+PN} = 2k_{\mathrm{off}}^P[PPN]$ |
| $P + N \leftrightharpoons PN$ | $K_A^N$ | $p_{P+N\to PN} = k_{\mathrm{on}}^N[P][N]$<br>$p_{PN\to P+N} = k_{\mathrm{off}}^N[PN]$ |
| $PP + N \leftrightharpoons PPN$ | $2K_A^N$ | $p_{PP+N\to PPN} = 2k_{\mathrm{on}}^N[PP][N]$<br>$p_{PPN\to PP+N} = k_{\mathrm{off}}^N[PPN]$ |
| $PPN + N \leftrightharpoons PNPN$ | $\gamma\frac{K_A^N}{2}$ | $p_{PPN+N\to NPPN} = \gamma k_{\mathrm{on}}^N[PPN][N]$<br>$p_{NPPN\to PPN+N} = 2k_{\mathrm{off}}^N[NPPN]$ |
| $PN + PN \leftrightharpoons PNPN$ | $\gamma K_A^P$ | $p_{PN+PN\to NPPN} = \gamma k_{\mathrm{on}}^P[PN][PN]$<br>$p_{NPPN\to PN+PN} = k_{\mathrm{off}}^P[NPPN]$ |
| $P + S \leftrightharpoons PS$ | $K_A^S$ | $p_{P+S\to PS} = k_{\mathrm{on}}^S[P][S]$<br>$p_{PS\to P+S} = k_{\mathrm{off}}^S[PS]$ |
| $PP + S \leftrightharpoons PPS$ | $2K_A^S$ | $p_{PP+S\to PPS} = 2k_{\mathrm{on}}^S[PP][S]$<br>$p_{PPS\to PP+S} = k_{\mathrm{off}}^S[PPS]$ |
| $PS + P \leftrightharpoons PPS$ | $2K_A^P$ | $p_{PS+P\to PPS} = 2k_{\mathrm{on}}^P[PS][P]$<br>$p_{PPS\to PS+P} = k_{\mathrm{off}}^P[PPS]$ |
| $PSN + P \leftrightharpoons PPSN$ | $2K_A^P$ | $p_{PSN+P\to PPSN} = 2k_{\mathrm{on}}^P[PSN][P]$<br>$p_{PPSN\to PSN+P} = k_{\mathrm{off}}^P[PPSN]$ |
| $PS + N \leftrightharpoons PSN$ | $\gamma K_A^N$ | $p_{PS+N\to PSN} = \gamma k_{\mathrm{on}}^N[PS][N]$<br>$p_{PSN\to PS+N} = k_{\mathrm{off}}^N[PSN]$ |
| $PPS + N \leftrightharpoons PNPS$ | $\gamma K_A^N$ | $p_{PPS+N\to PNPS} = \gamma k_{\mathrm{on}}^N[PPS][N]$<br>$p_{PNPS\to PPS+N} = k_{\mathrm{off}}^N[PNPS]$ |
| $PPS + N \leftrightharpoons PPSN$ | $\gamma K_A^N$ | $p_{PPS+N\to PPSN} = \gamma k_{\mathrm{on}}^N[PPS][N]$<br>$p_{PPSN\to PPS+N} = k_{\mathrm{off}}^N[PPSN]$ |
| $PNPS + N \leftrightharpoons PNPSN$ | $\gamma K_A^N$ | $p_{PNPS+N\to PNPSN} = \gamma k_{\mathrm{on}}^N[PNPS][N]$<br>$p_{PNPSN\to PNPS+N} = k_{\mathrm{off}}^N[PNPSN]$ |
| $PPSN + N \leftrightharpoons PNPSN$ | $\gamma K_A^N$ | $p_{PPSN+N\to PNPSN} = \gamma k_{\mathrm{on}}^N[PPSN][N]$<br>$p_{PNPSN\to PPSN+N} = k_{\mathrm{off}}^N[PNPSN]$ |
| $PN + S \leftrightharpoons PSN$ | $\gamma K_A^S$ | $p_{PN+S\to PSN} = \gamma k_{\mathrm{on}}^S[PN][S]$<br>$p_{PSN\to PN+S} = k_{\mathrm{off}}^S[PSN]$ |

| | | |
|---|---|---|
| $PPN + S \leftrightharpoons PNPS$ | $\gamma K_A^S$ | $p_{PPN+S \to PNPS} = \gamma k_{\text{on}}^S [PPN][S]$ |
| | | $p_{PNPS \to PPN+S} = k_{\text{off}}^S [PNPS]$ |
| $PPN + S \leftrightharpoons PPSN$ | $\gamma K_A^S$ | $p_{PPN+S \to PPSN} = \gamma k_{\text{on}}^S [PPN][S]$ |
| | | $p_{PPSN \to PPN+S} = k_{\text{off}}^S [PPSN]$ |
| $PNPN + S \leftrightharpoons PNPSN$ | $2\gamma K_A^S$ | $p_{PNPN+S \to PNPSN} = 2\gamma k_{\text{on}}^S [PNPN][S]$ |
| | | $p_{PNPSN \to PNPN+S} = k_{\text{off}}^S [PNPSN]$ |
| $PN + PS \leftrightharpoons PNPS$ | $2\gamma K_A^P$ | $p_{PN+PS \to PNPS} = 2\gamma k_{\text{on}}^P [PN][PS]$ |
| | | $p_{PNPS \to PN+PS} = k_{\text{off}}^P [PNPS]$ |
| $PSN + PN \leftrightharpoons PNPSN$ | $2\gamma K_A^P$ | $p_{PSN+PN \to PNPSN} = 2\gamma k_{\text{on}}^P [PSN][PN]$ |
| | | $p_{PNPSN \to PSN+PN} = k_{\text{off}}^P [PNPSN]$ |

*Table S3*

#### Table S4 Reactions with their association constants and propensities for environment DNA + clusTarg

1D reactions are colored red and their $K_A$'s follow the definition of Eq. SI. 5. The blue equations are unimolecular or can be thought of as effectively conformational changes. Importantly, the species $PPS_2$ is distinct from $PSPS$; in $PSPS$, each protein has bound specifically to the DNA, whereas in $PPS_2$, only one protein has bound to the DNA. They are not bimolecular reactions as they do not involve diffusion/search to contact, and the rates must therefore have units of $s^{-1}$, where $c_0 = 1M$ is the standard state concentration. The use of $c_0$ ensures that the binding between P and S has the same free energy barrier as it would for a bimolecular reaction and will give the same equilibrium as if both proteins bound both specific sites simultaneously (e.g. $PP + S_2 \leftrightharpoons SPPS$, with $K_A^{PSPS} = \frac{[PSPS]}{[PP][S_2]} = 2\exp\left(-\frac{\Delta G^{PSPS}}{k_B T}\right)/c_0 = 2\exp\left(-\frac{2\Delta G^{PS}}{k_B T}\right)/c_0$), rather than using a two-step scheme (e.g. $PP + S_2 \leftrightharpoons PPS_2 \leftrightharpoons SPPS$), as we do here.

Since $S_2$ has two targets, reactions involving binding to $S_2$ always have the concentration of $[S_2]$ multiplied by 2. These factors are colored green.

| Reaction | $K_A$ | Forward reaction propensity / Backward reaction propensity |
|---|---|---|
| $P + P \leftrightharpoons PP$ | $K_A^P$ | $p_{P+P\to PP} = k_{\text{on}}^P[P]^2$ |
| | | $p_{PP\to P+P} = k_{\text{off}}^P[PP]$ |
| $PN + P \leftrightharpoons PPN$ | $2K_A^P$ | $p_{P+PN\to PPN} = 2k_{\text{on}}^P[P][PN]$ |
| | | $p_{PPN\to P+PN} = 2k_{\text{off}}^P[PPN]$ |
| $P + N \leftrightharpoons PN$ | $K_A^N$ | $p_{P+N\to PN} = k_{\text{on}}^N[P][N]$ |
| | | $p_{PN\to P+N} = k_{\text{off}}^N[PN]$ |
| $PP + N \leftrightharpoons PPN$ | $2K_A^N$ | $p_{PP+N\to PPN} = 2k_{\text{on}}^N[PP][N]$ |
| | | $p_{PPN\to PP+N} = k_{\text{off}}^N[PPN]$ |
| $PPN + N \leftrightharpoons PNPN$ | $\gamma\frac{K_A^N}{2}$ | $p_{PPN+N\to NPPN} = \gamma k_{\text{on}}^N[PPN][N]$ |
| | | $p_{NPPN\to PPN+N} = 2k_{\text{off}}^N[NPPN]$ |
| $PN + PN \leftrightharpoons PNPN$ | $\gamma K_A^P$ | $p_{PN+PN\to NPPN} = \gamma k_{\text{on}}^P[PN][PN]$ |
| | | $p_{NPPN\to PN+PN} = k_{\text{off}}^P[NPPN]$ |
| $P + S_2 \leftrightharpoons PS_2$ | $2K_A^S$ | $p_{P+S_2\to PS_2} = 2\cdot k_{\text{on}}^S[P][S_2]$ |
| | | $p_{PS_2\to P+S_2} = k_{\text{off}}^S[PS_2]$ |
| $PP + S_2 \leftrightharpoons PPS_2$ | $2\cdot 2K_A^S$ | $p_{PP+S_2\to PPS_2} = 2\cdot 2k_{\text{on}}^S[PP][S_2]$ |
| | | $p_{PPS_2\to PP+S_2} = k_{\text{off}}^S[PPS_2]$ |
| $PS_2 + P \leftrightharpoons PPS_2$ | $2\cdot 2K_A^P$ | $p_{PS_2+P\to PPS_2} = 2\cdot 2k_{\text{on}}^P[PS_2][P]$ |
| | | $p_{PPS_2\to PS_2+P} = k_{\text{off}}^P[PPS_2]$ |
| $PS_2N + P \leftrightharpoons PPS_2N$ | $2K_A^P$ | $p_{PS_2N+P\to PPS_2N} = 2k_{\text{on}}^P[PS_2N][P]$ |

| | | |
|---|---|---|
| | | $p_{PPS_2N\to PS_2N+P} = k_{\text{off}}^P[PPS_2N]$ |
| $PS_2 + N \leftrightharpoons PS_2N$ | $\gamma K_A^N$ | $p_{PS_2+N\to PS_2N} = \gamma k_{\text{on}}^N[PS_2][N]$ |
| | | $p_{PS_2N\to PS_2+N} = k_{\text{off}}^N[PS_2N]$ |
| $PPS_2 + N \leftrightharpoons PNPS_2$ | $\gamma K_A^N$ | $p_{PPS_2+N\to PNPS_2} = \gamma k_{\text{on}}^N[PPS_2][N]$ |
| | | $p_{PNPS_2\to PPS_2+N} = k_{\text{off}}^N[PNPS_2]$ |
| $PPS_2 + N \leftrightharpoons PPS_2N$ | $\gamma K_A^N$ | $p_{PPS_2+N\to PPS_2N} = \gamma k_{\text{on}}^N[PPS_2][N]$ |
| | | $p_{PPS_2N\to PPS_2+N} = k_{\text{off}}^N[PPS_2N]$ |
| $PNPS_2 + N \leftrightharpoons PNPS_2N$ | $\gamma K_A^N$ | $p_{PNPS_2+N\to PNPS_2N} = \gamma k_{\text{on}}^N[PNPS_2][N]$ |
| | | $p_{PNPS_2N\to PNPS_2+N} = k_{\text{off}}^N[PNPS_2N]$ |
| $PPS_2N + N \leftrightharpoons PNPS_2N$ | $\gamma K_A^N$ | $p_{PPS_2N+N\to PNPS_2N} = \gamma k_{\text{on}}^N[PPS_2N][N]$ |
| | | $p_{PNPS_2N\to PPS_2N+N} = k_{\text{off}}^N[PNPS_2N]$ |
| $PSPS + N \leftrightharpoons PSPSN$ | $2\gamma K_A^N$ | $p_{PSPS+N\to PSPSN} = 2\gamma k_{\text{on}}^N[PSPS][N]$ |
| | | $p_{PSPSN\to PSPS+N} = k_{\text{off}}^N[PSPSN]$ |
| $PSPSN + N \leftrightharpoons PSNPSN$ | $\gamma K_A^N/2$ | $p_{PSPSN+N\to PSNPSN} = \gamma k_{\text{on}}^N[PSPSN][N]$ |
| | | $p_{PSNPSN\to PSPSN+N} = 2k_{\text{off}}^N[PSNPSN]$ |
| $PN + S_2 \leftrightharpoons PS_2N$ | $2\gamma K_A^S$ | $p_{PN+S_2\to PS_2N} = 2\gamma k_{\text{on}}^S[PN][S_2]$ |
| | | $p_{PS_2N\to PN+S_2} = k_{\text{off}}^S[PS_2N]$ |
| $PPN + S_2 \leftrightharpoons PNPS$ | $2\gamma K_A^S$ | $p_{PPN+S_2\to PNPS_2} = 2\gamma k_{\text{on}}^S[PPN][S_2]$ |
| | | $p_{PNPS_2\to PPN+S_2} = k_{\text{off}}^S[PNPS_2]$ |
| $PPN + S_2 \leftrightharpoons PPSN$ | $2\gamma K_A^S$ | $p_{PPN+S_2\to PPSN} = 2\gamma k_{\text{on}}^S[PPN][S_2]$ |
| | | $p_{PPS_2N\to PPN+S_2} = k_{\text{off}}^S[PPS_2N]$ |
| $PNPN + S_2 \leftrightharpoons PNPSN$ | $2\cdot 2\gamma K_A^S$ | $p_{PNPN+S_2\to PNPSN} = 22\gamma k_{\text{on}}^S[PNPN][S_2]$ |
| | | $p_{PNPSN\to PNPN+S_2} = k_{\text{off}}^S[PNPS_2N]$ |
| $PN + PS_2 \leftrightharpoons PNPS_2$ | $2\gamma K_A^P$ | $p_{PN+PS_2\to PNPS_2} = 2\gamma k_{\text{on}}^P[PN][PS_2]$ |
| | | $p_{PNPS_2\to PN+PS_2} = k_{\text{off}}^P[PNPS_2]$ |
| $PS_2N + PN \leftrightharpoons PNPS_2N$ | $2\gamma K_A^P$ | $p_{PS_2N+PN\to PNPS_2N} = 2\gamma k_{\text{on}}^P[PS_2N][PN]$ |
| | | $p_{PNPS_2N\to PS_2N+PN} = k_{\text{off}}^P[PNPS_2N]$ |
| $PPS_2 \leftrightharpoons PSPS$ | $K_A^S c_0/2$ | $p_{PPS_2\to PSPS} = k_{\text{on}}^S C_0[PPS_2]$ |
| | | $p_{PSPS\to PPS_2} = 2k_{\text{off}}^S[SPPS]$ |
| $PPS_2N \leftrightharpoons PSPSN$ | $K_A^S c_0$ | $p_{PPS_2N\to PSPSN} = k_{\text{on}}^S C_0[PPS_2N]$ |
| | | $p_{PSPSN\to PPS_2N} = k_{\text{off}}^S[PSPSN]$ |
| $PNPS_2 \leftrightharpoons PSPSN$ | $K_A^S c_0$ | $p_{PNPS_2\to PSPSN} = k_{\text{on}}^S C_0[PNPS_2]$ |
| | | $p_{PSPSN\to PNPS_2} = k_{\text{off}}^S[PSPSN]$ |
| $PNPS_2N \leftrightharpoons PSNPSN$ | $K_A^S c_0/2$ | $p_{PNPS_2N\to PSNPSN} = k_{\text{on}}^S C_0[PNPS_2N]$ |
| | | $p_{PSNPSN\to PNPS_2N} = 2k_{\text{off}}^S[PSNPSN]$ |

Table S4

### Table S5 Parameter used for cross-validation in NERDSS simulation

| Parameter | Unit | Value or range | Parameter | Unit | Value or range |
| --- | --- | --- | --- | --- | --- |
| Water box - x | nm | 1050 | $k_{\text{on}}^{N}$ | $\text{nm}^3/\text{s}$ | 200.00 |
| Water box - y | nm | 31.623 | $k_{\text{off}}^{N}$ | $\text{s}^{-1}$ | 1.2044 |
| Water box - z | nm | 31.623 | $\sigma_{P+N}$ | nm | 1.00 |
| $P$ copynumber | 1 | 20 | $h^2$ | $\text{nm}^2$ | 10 |
| $N$ copynumber | 1 | 600 | $k_{\text{off}}^{P}$ | $\text{s}^{-1}$ | 0.4000 |
| $k_{\text{on}}^{P}$ | $\text{nm}^3/\text{s}$ | 0, 66.423, 664.24, 6642.9 | $\sigma_{P+P}$ | nm | 2.00 |

*Table S55.*

### Table S6 Parameter used for selectivity between two segments of DNA

The association rates of dimers are: $k_{\mathrm{on}}^{P} = 0$, $k_{\mathrm{on}}^{P} = 6.6423 \times 10^{-6}\ \mathrm{nm}^3/\mathrm{s}$, $k_{\mathrm{on}}^{P} = 6.6423 \times 10^{-5}\ \mathrm{nm}^3/\mathrm{s}$, $k_{\mathrm{on}}^{P} = 6.6424 \times 10^{-4}\ \mathrm{nm}^3/\mathrm{s}$, $k_{\mathrm{on}}^{P} = 6.6431 \times 10^{-3}\ \mathrm{nm}^3/\mathrm{s}$, $k_{\mathrm{on}}^{P} = 6.6651 \times 10^{-2}\ \mathrm{nm}^3/\mathrm{s}$, $k_{\mathrm{on}}^{P} = 6.6727 \times 10^{-1}\ \mathrm{nm}^3/\mathrm{s}$. We use fixed coordinates for molecule $S$ and $nuc$, such that $nuc$ molecules creates two DNA segments and $S$ molecules are distributed as described in Figure 6-a. Molecules $P$ and molecules $N$ are initialized randomly.

| Parameter | Unit | Value or range | Parameter | Unit | Value or range |
|---|---|---|---|---|---|
| Water box - x | nm | 106 | Water box - y | nm | 31.623 |
| Water box - z | nm | 31.623 | $h^2$ | $\mathrm{nm}^2$ | 10 |
| $P$ copynumber | 1 | 2 | $S$ copynumber | 1 | 4 |
| $N$ copynumber | 1 | 44 | $nuc$ copynumber | 1 | 3 |
| $\sigma_{P+P}$ | nm | 1.40 | $k_{\mathrm{on}}^{S}$ | $\mathrm{nm}^3/\mathrm{s}$ | 0.1005 |
| $\sigma_{P+N}$ | nm | 1.00 | $k_{\mathrm{off}}^{S}$ | $\mathrm{s}^{-1}$ | 6.0541 |
| $\sigma_{P+S}$ | nm | 1.00 | $k_{\mathrm{on}}^{N}$ | $\mathrm{nm}^3/\mathrm{s}$ | 200.00 |
| $\sigma_{P+nuc}$ | nm | 5.50 | $k_{\mathrm{off}}^{N}$ | $\mathrm{s}^{-1}$ | 1.2044 |
| $\sigma_{N+nuc}$ | nm | 5.50 | $k_{\mathrm{off}}^{P}$ | $\mathrm{s}^{-1}$ | 0.4000 |

*Table S66.*

#### Table S7 Parameter used for selectivity between four segments of DNA

We use fixed coordinates for molecule $S$. Four DNA segments are separated spatially far enough to avoid reactions between proteins on different DNA. $S$ molecules are distrubted as described in Figure S11-a. Molecules $P$ and molecules $N$ are initialized randomly.

| Parameter | Unit | Value or range | Parameter | Unit | Value or range |
|---|---|---|---|---|---|
| Water box - x | nm | 200 | Water box - y | nm | 63.246 |
| Water box - z | nm | 63.246 | $h^2$ | $\mathrm{nm}^2$ | 10 |
| $P$ copynumber | 1 | 2, 4 | $S$ copynumber | 1 | 10 |
| $N$ copynumber | 1 | 446 | $k_{\mathrm{on}}^{S}$ | $\mathrm{nm}^3/s$ | 0.1005 |
| $\sigma_{P+P}$ | nm | 1.40 | $k_{\mathrm{off}}^{S}$ | $\mathrm{s}^{-1}$ | 6.0541 |
| $\sigma_{P+N}$ | nm | 1.00 | $k_{\mathrm{on}}^{N}$ | $\mathrm{nm}^3/s$ | 0.0200 |
| $\sigma_{P+S}$ | nm | 1.00 | $k_{\mathrm{off}}^{N}$ | $\mathrm{s}^{-1}$ | 1205.7 |
| $k_{\mathrm{on}}^{P}$ | $\mathrm{nm}^3/s$ | 0.1665 | $k_{\mathrm{off}}^{P}$ | $\mathrm{s}^{-1}$ | 10.0316 |

*Table S77.*

### Figure S1. Definition of nonspecific binding.

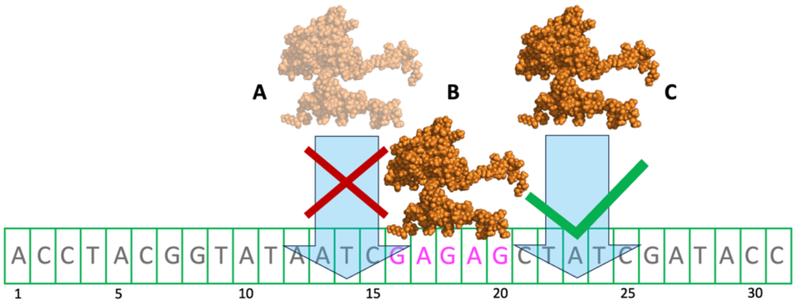


*Figure S1* We define protein binding sites on DNA as non-overlapping sequences with the length of the protein's footprint length. This segment has $N_{bp} = 31$ basepairs, and the footprint length of TF is $l_{fp} = 5$ (target site is the magenta sequences). When one protein associates to DNA, there are $N_{bp} - l_{fp} + 1 = 27$ possible positions. However, the maximum number of proteins that can bind is $\lfloor N_{bp}/l_{bp} \rfloor = 6$. Here, when protein B is bound, protein C can bind but A cannot.

### Figure S2. Spatial simulation yields same dynamics and equilibrium as nonspatial simulations.

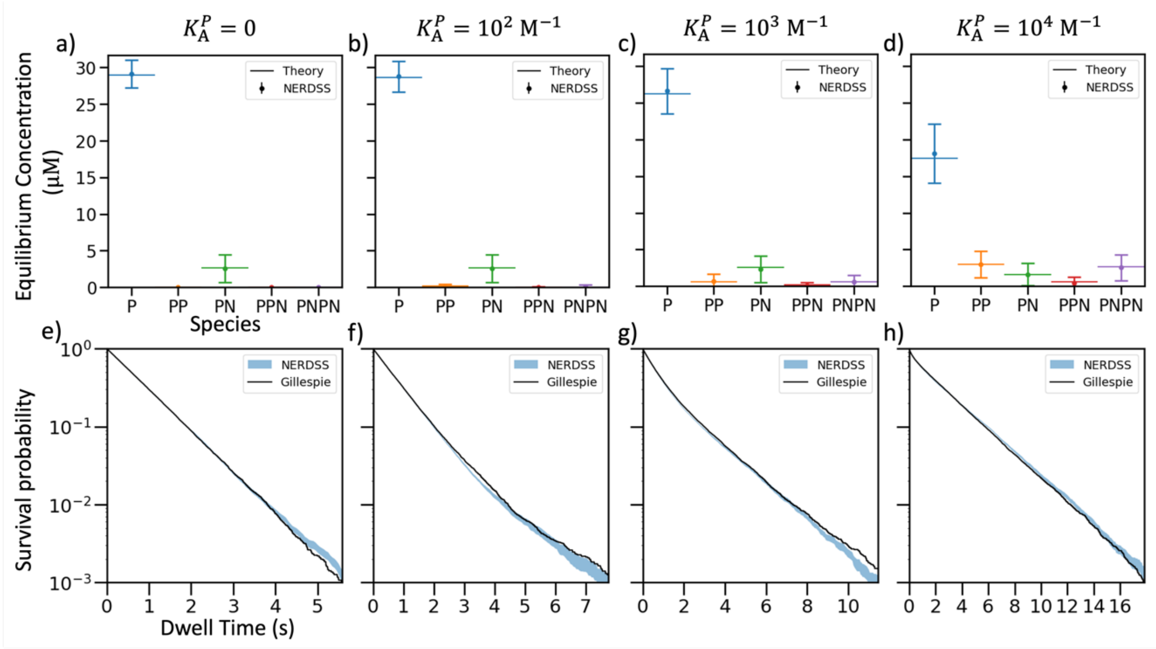


*Figure S2* NERDSS results that have spatial resolution agree with nonspatial methods. a) – d): The equilibrium concentration of proteins in different states from NERDSS simulations (scatters with error bars, see SXII.D) agree with our analytical predictions (given by SXII.A, horizontal lines). No dimer states of proteins are observed with $K_A^P = 0$. e) – h): The dwell time obtained from NERDSS (blue shaded region labels the mean $\pm$ standard error, see SXII.D) agree with Gillespie simulation's results (black curves).

### Figure S3. Environment: DNA (protein + nonspecific DNA)

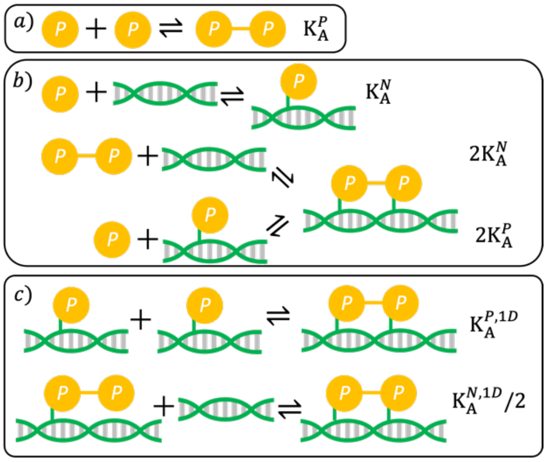


*Figure S3* A schematic graph of environment DNA. This model contains proteins $P$ (yellow circles) and random DNA sequences $N$ (green helix with gray base pairs). They can combine to form 4 additional species, for 6 species total: $P, N, PP, PN, PPN, NPPN$. The 6 possible reactions are described by S2.1~S2.6. Box a) and box b) list the 3D reaction and the 3D-1D transitions, respectively. Box c) shows the 1D reactions. For the limit of no dimerizations $K_A^P \rightarrow 0$ , only monomers exist, and 1 reaction occurs: $P + N \rightleftharpoons PN$. For the limit of irreversible dimerization, $K_A^P \rightarrow \infty$: no protein monomers exist at equilibrium and 2 reactions occur: $PP + N \rightleftharpoons PPN$, and $PPN + N \rightleftharpoons PNPN$ can occur. The association constants are listed on the right of each reaction. The green bonds represent nonspecific associations, and the yellow bonds represent protein-protein bindings.

### Figure S4. Environment: DNA+targ (protein + nonspecific DNA and two separated targets)

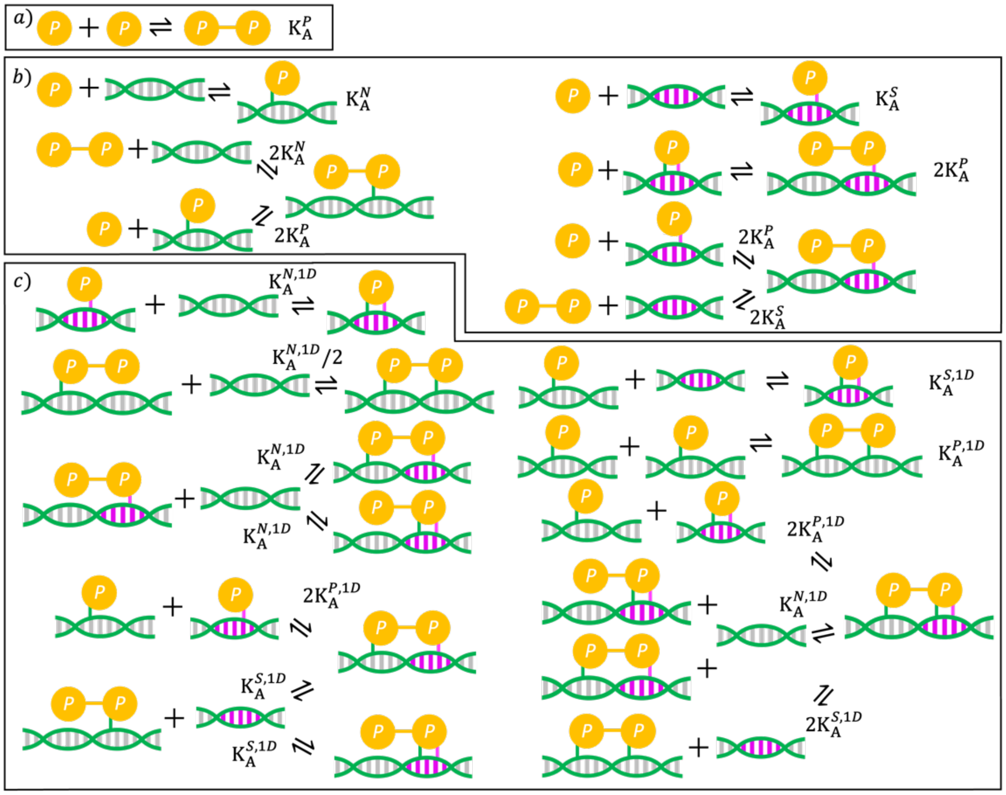


Figure S4 *A schematic graph of environment DNA+targ*. This model contains protein (yellow *circles*), random DNA sequence (gray base), and target DNA sequence (magenta base pairs). We show all 16 possible reactions that occur between proteins and between protein and DNA. Box a) shows 3D reactions, box b) shows 3D-1D transitions, and box c) shows 1D reactions. Compared to model DNA, the additional reactions are: S3.7 ~ S3.21. The association constants are labeled next to each reaction. The green bonds represent nonspecific associations, the magenta bonds represent protein-target bindings, and the yellow bond represent protein-protein bindings.

### Figure S5. Environment: DNA+clusTarg (protein + nonspecific DNA and a cluster of two targets)

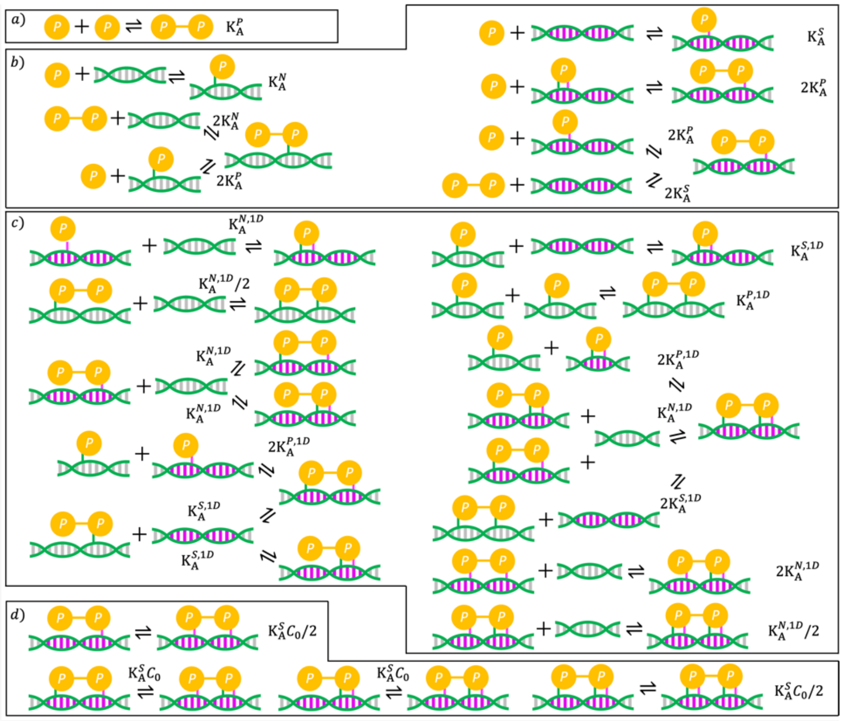

*Figure S5* A schematic graph of environment DNA+clusTarg. Different from model B (Figure ), the bindings of one target ($S$) are replaced by bindings of two clustered targets ($S_2$) (e.g. $P + S \rightleftharpoons PS$ is replaced with $P + S_2 \rightleftharpoons PS_2$). Besides, there are 6 more reactions. 2 are 1D nonspecific bindings ($SPPS + N \rightleftharpoons SPPSN$ and $SPPSN + N \rightleftharpoons NSPPSN$) in box c) and 4 are unimolecular reactions ($PPS_2 \rightleftharpoons SPPS$, $NPPS_2 \rightleftharpoons SPPSN$, $PPS_2N \rightleftharpoons SPPSN$, and $NPPS_2N \rightleftharpoons NSPPSN$) in box d). Association constants are labeled next to each reaction.

### Figure S6. The distribution of dwell times when $k_{\text{off}}^{P} = k^*$

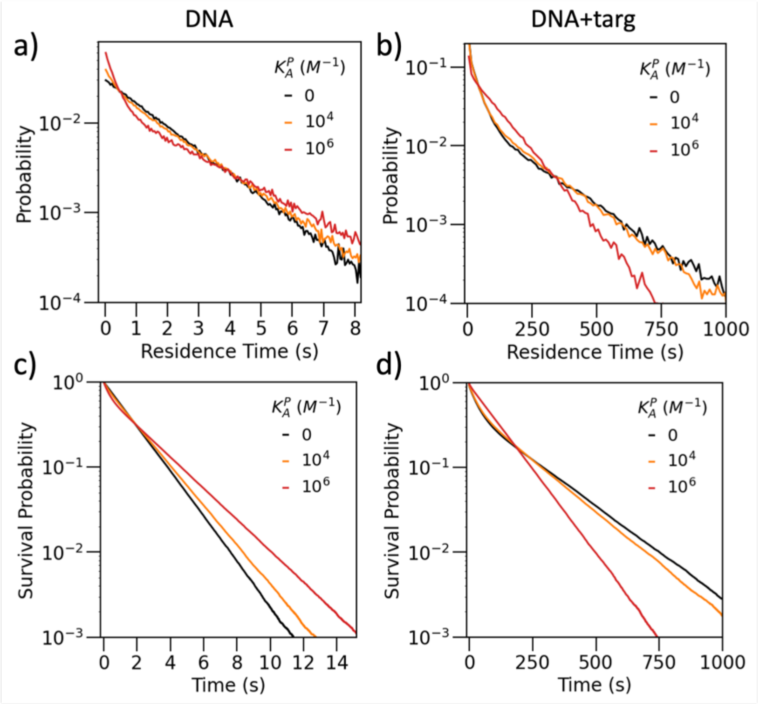


*Figure S6* a) and b) are the distribution of dwell times collected from Gillespie simulations of environment $DNA$ and $DNA + targ$, respectively. $k_{on}^{P}$'s are varied between different curves, and the red curves are for monomers thus represents intrinsic distribution of dwell times. Due to the discretization of proteins, real critical rates are slower than predicted $k^*$. c) and d) are the survival probabilities corresponding to a) and b), respectively. With isolated targets, there are intrinsically two dissociation patterns featuring specific binding and nonspecific binding (b). Dimerization increases the dwell time on nonspecific sites and the difference between the intrinsic short and long dissociation patterns gets vague. However, with only random DNA (a), dimerization introduces dissociation of colocalized proteins, a new short-lived pattern. In a) and c), $k^* = 0.1\ s^{-1}$, $k_{off}^{P} = 0.08\ s^{-1}$, and $\tau = 1.71 \pm 0.05\ s$. In b) and d), $k^* = 0.1\ s^{-1}$, $k_{off}^{P} = 0.055\ s^{-1}$, and $\tau = 102.2 \pm 0.6\ s$.

### Figure S7. The binary effect of dimerization on target occupancy and protein dwell time in environment $DNA + targ$ with high protein concentration.

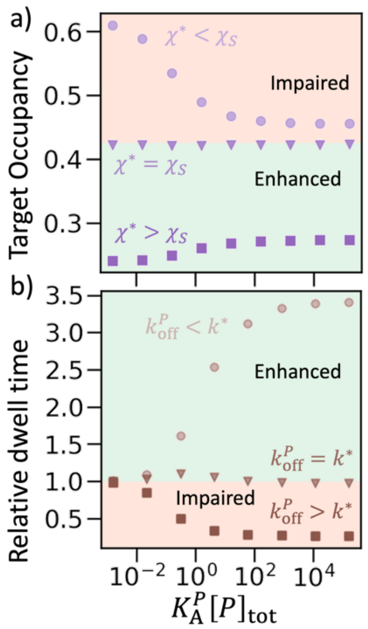


*Figure S2* With high protein concentration ($[P]_{tot} = [S]_{tot}$), dimerization may impair or enhance occupancy and dwell time in environment $DNA + targ$. a) Although Eq. 10 in main text does not apply under high $[P]_{tot}$, we manually fitted $\chi^* = \chi_S = 0.199$ is the critical partition coefficient such that dimerization does not change target occupancy. b) Eq. 15 in main text does not apply and we manually fitted $k_{off}^P = k^* = 0.25\ s^{-1}$ is the critical rate where strong reversible dimers have the same dwell time as monomers. However, with weak dimerization, $k_{off}^P = k^*$ still results in a slight enhancement since some DNA sites have not been occupied by proteins, which increases $k^*$. These unoccupied DNA sites will be occupied with stronger dimerization since dimerization can enhance the recruitment fraction. Here the dimer dissociation rates are $0.01\ s^{-1}$, $0.25\ s^{-1}$, and $5\ s^{-1}$, for circles, triangles, and squares, respectively.

### Figure S8. In environment $DNA + clusTarg$, dimerization always enhances protein-DNA binding and target occupancy is enhanced by higher protein concentration.

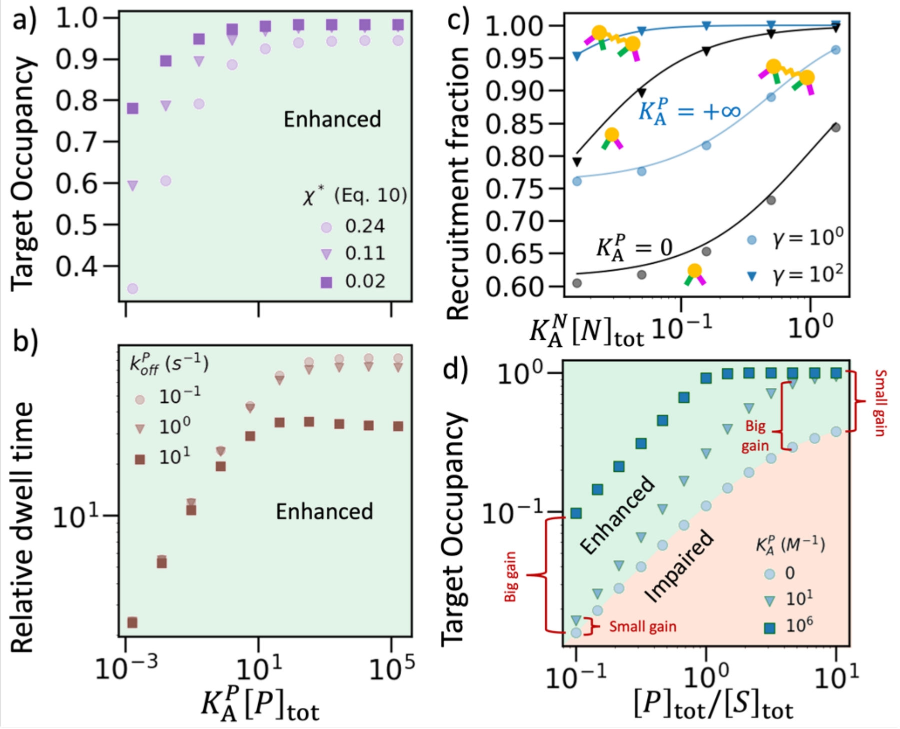


*Figure S3* In this figure, numerical results (scatters) are obtained by solving ODEs (see SXII.A). With high protein concentration ($[P]_{tot} = [S]_{tot}$), dimerization always enhance target occupancy a) and protein dwell time b) in environment $DNA + clusTarg$. c) From monomer to irreversible dimer, the recruitment fraction is hugely enhanced, even with small dimensionality enhancement, $\gamma = 1$ and $K_A^N[N]_{tot} \sim 10^{-2}$. Theoretical results (curves) are given by Eq. (53) in SI (dimer) and Eq. (43) in SI (monomer). d) Higher protein concentration cooperating with dimerization always result in higher target occupancy. The strong dimers (blue squares) are nearly linear with $[P]_{tot}/[S]_{tot}[P]_{tot}/[S]_{tot}$ until full occupancy (1.0) because they reach the maximal occupancy possible for excess target sites. For weaker dimers (green triangles) the gain compared to monomers (circles) is highest for $\frac{[P]_{tot}}{[S]_{tot}} \sim \frac{[P]_{tot}}{[S]_{tot}} \sim 2$. This non-monotonic gain where intermediate $K_A^P$ optimizes the reaction balance towards dimer formation is because at very high protein concentrations, monomers effectively occupy targets without dimerization, and at low concentrations, dimers are infrequent. Scatters are numerical results from Eq. (50) in SI.

### Figure S9. Cartoon illustrating how the intensities from ChIP-Seq measurements scale for clustered targets.

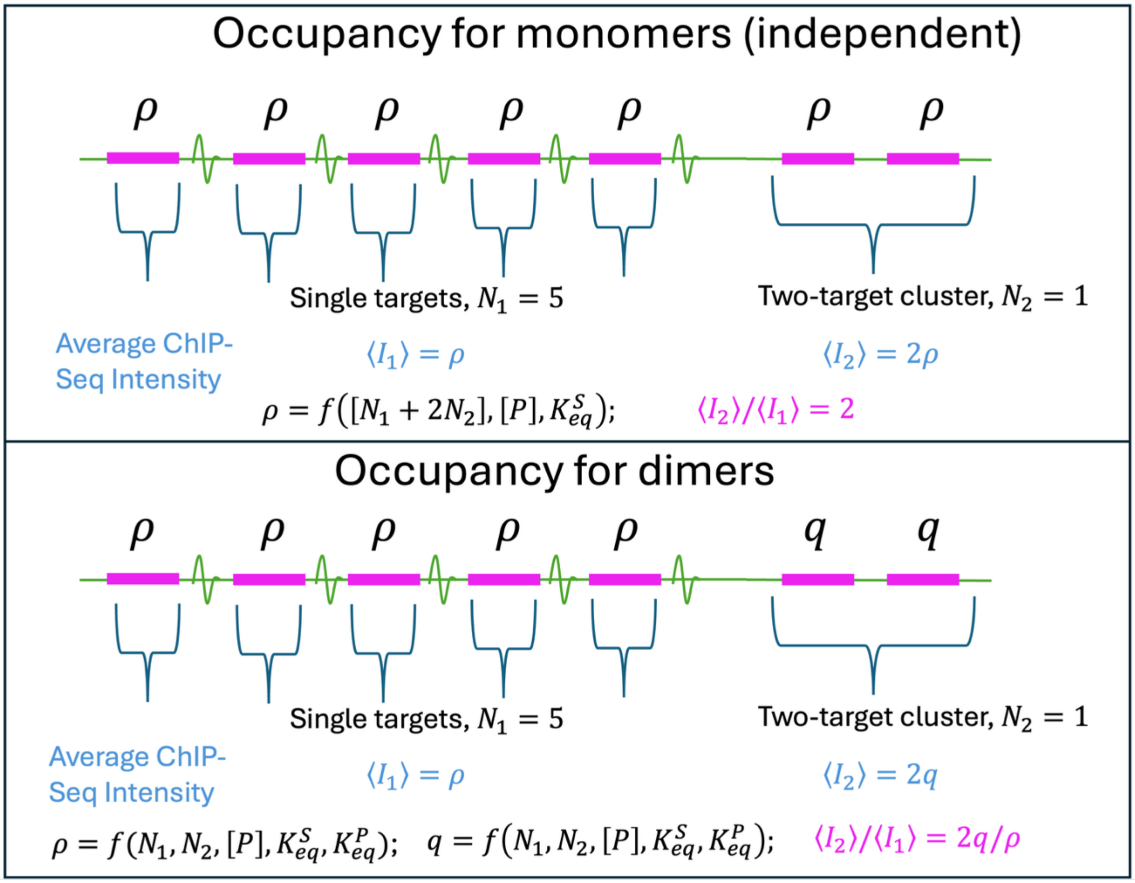


**Figure S9. Cartoon illustrating how the intensities from ChIP-Seq measurements scale for clustered targets.** In the upper panel, monomers bind independently to all sites, regardless of whether they are clustered, producing a uniform occupancy $\rho$ on all targets. Because peak intensities are integrated over both targets in a cluster of 2, the intensity of a two-target cluster, $\langle I_2 \rangle$, is double that of a single target $\langle I_1 \rangle$. Although $\rho$ will vary with the total number of target sites, proteins, and DNA affinity, the ratio of intensities is constant, $\langle I_2 \rangle / \langle I_1 \rangle = 2$. In contrast, the lower panel illustrates that for correlated interactions, such as driven by dimerization, the occupancy on two-target clusters and single targets both depend on all concentrations and binding parameters. As a result, the ratio of intensities is not a constant value, with $\langle I_2 \rangle / \langle I_1 \rangle = 2q/\rho$, and typically $q > \rho$.

### Figure S10. Partitioning of proteins on DNA segments with target clusters in various sizes

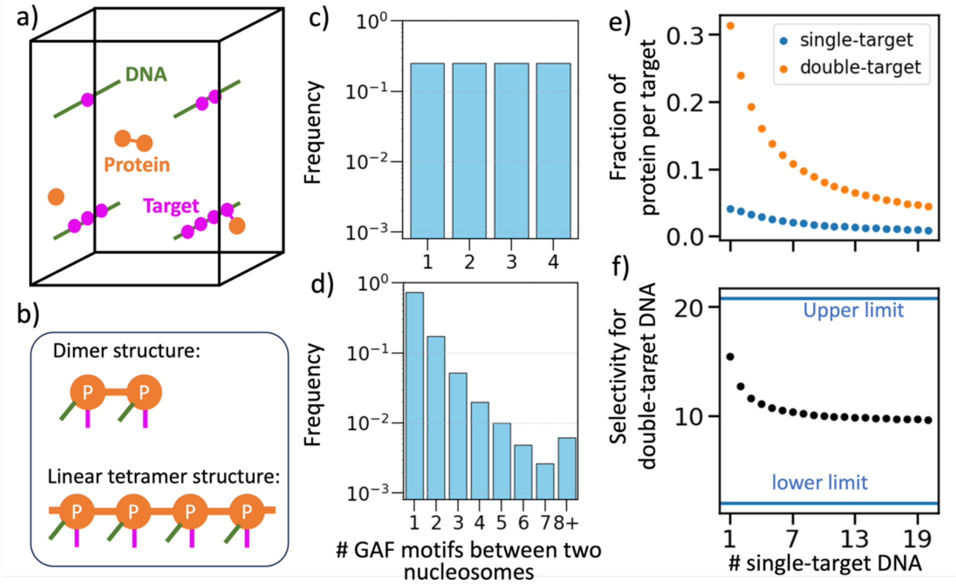


*Figure S10* Partitioning of proteins on DNA segments with target clusters in various sizes. a) In our simulation, each DNA segment is a fiber isolated from others as illustrated. These fibers are initialized at designed positions. No nucleosomes are added. Proteins are initialized randomly. b) Each protein has one interface to react with other proteins which allows the formation of dimers. For tetramers, each protein has two interfaces such that the oligomer can grow linearly. There are four total proteins in the system and thus the largest oligomer is a tetramer. C) In our simulations, we have one segment with a single target, one segment with a cluster of 2 targets, one segment with a cluster of 3, and one with a segment of 4, so they all have equal frequency. D) The frequency of target clusters for the GAF motif in vivo was calculated as described in section SXIII. Most motifs are single targets, and 90% of targets are in clusters of 3 or less. For independent monomer binding, this does not affect relative ChIP-Seq intensities and thus selectivity for clustered targets (Fig S10), but for oligomer forming proteins, it does. E) Using simulations, we show that for dimer-forming proteins, the fraction of proteins per target is higher on a cluster of two targets (orange) relative to single targets (blue), as expected. We then increase the number of single target sites $N_1$ on the x-axis, with double-target sites $N_2$ fixed, and the number of proteins is also increased to keep the ratio of proteins to target numbers constant, ($N_{pro} = N_1 + 2N_2$). We see that the distribution of proteins on the targets shifts. Occupancy per target decreases for all targets, but moreso for the double-target. F) Using the data from (E) we see that the selectivity for the double-target DNA is reduced with an excess of single-target DNA, but it is still much higher than 1. In F), the upper limit is given by Eq. 64 and the lower limit is literally 2.

### Dataset S1 (separate file)

In “Parameters_rateEquations.xlsx”, we listed all parameters used in ODE solutions and theoretical results for Figure 2~7, Figure S5~S8, and Figure S10. For parameters that have no more than 3 values, the values are listed in the spreadsheet. These parameters are for different curves in corresponding plots. Some other parameters change as the x-axis or yaxis change, and the values of these parameters aligns with the axes in corresponding plots.

#### Software S1 (separate file)

In the GitHub repo (https://github.com/sangmk/dimerEnhanceProteinDNA), you can find 1) Simulation input files and instructions on how to run simulations, and 2) Jupyter notebooks that generate figures.
